# Large parks and city-wide tree cover boost butterfly diversity across 22 major U.S. cities

**DOI:** 10.64898/2026.07.02.736135

**Authors:** Jens Ulrich, Yan Yin Jenny Cheung, Christopher T. Cosma, Heather Kharouba, Laura Melissa Guzman

**Affiliations:** Department of Biology, University of Ottawa, Ottawa, Ontario, K1N615, Canada; Department of Entomology, Cornell University, Ithaca, New York 14853, USA; Conservation Biology Institute, Corvallis, OR, 97333, USA

**Author notes:** Corresponding Author: Jens Ulrich.

**Keywords:** Urban ecology, Community science, Occupancy models, Equilibrium Theory of Island Biogeography, Insect conservation

## Abstract

Accelerating global urbanization necessitates a better understanding of how to manage cities that promote biodiversity. However, we currently lack multi-year, multi-city studies, which limits a generalizable understanding of how both within and between city differences impact the spatial and temporal dynamics of urban biodiversity. Here, we tested hypotheses about the drivers of butterfly diversity within and across urban parks by applying Bayesian occupancy models to five years of iNaturalist community science data from 2,550 parks in 22 major U.S. cities. We found that cities with bigger parks supported more species per park, including more disturbance- and edge-avoidant species. This was driven by a positive effect of park size on butterfly species colonization rates. We also found that attributes of habitat quality (plant diversity within parks and tree cover surrounding parks) contributed to butterfly species occupancy. Park connectivity increased species persistence, but the overall effects on butterfly species occupancy varied across cities. Finally, we found that the total area of tree cover throughout a city, rather than the size or connectivity of individual parks, was the primary determinant of city-wide diversity: Increasing total tree canopy cover from below-average (*∼* 6%) to above-average (*∼* 22%) increased city-wide species richness by *∼* 10%. These findings highlight the need for cities to maintain large parks while also increasing city-wide tree cover to support biodiversity across local to regional scales. By integrating high-resolution community science data across the continental U.S., this study provides mechanistic insight into how cross-scale processes shape urban biodiversity dynamics and identifies generalizable recommendations for improving urban conservation management.

**Significance statement:** Advancing urban sustainability hinges on better understanding how to promote biodiver-sity in urban landscapes. Yet, it is largely unclear how urban biodiversity is affected by both within- and between-city differences. Applying Bayesian occupancy models to 5 years of iNaturalist community science butterfly data from 22 U.S. cities, we found that cities with larger parks supported more species per park, including species of higher conservation concern. Simultaneously, increasing city-wide tree canopy cover area, including area both within and outside of parks, increased total city-wide butterfly diversity. Integrating biodi-versity conservation recommendations from this study into urban management planning will reinforce the benefits of participating in community science programs, thereby strengthening positive feedbacks between the health of people and nature.

## 1 Introduction

Global urban land cover is projected to triple by 2050, making sustainable urban planning an increasingly important lever for limiting habitat loss and biodiversity declines [1]. Simultaneously, urban biodiversity is necessary for the health, resilience, and well-being of urban communities [2]. To mitigate widespread biodiversity losses and maximize interconnected benefits to people, United Nations Global Sustainable Development Goals emphasize the urgent need to create biodiversity-rich areas in and around cities [3].

Promoting urban insect diversity is an especially important pathway towards urban sustainability. Insects provide numerous ecological, economic, and cultural services including pollination, nutrient cycling, and opportunities for people to strengthen connections with nature [4]. Unfortunately, many insect species face widespread declines [5, 6], with urbanization imposing particularly strong ecological filters that shape the composition and persistence of insect communities [7, 8]. For example, urban areas are dominated by impervious surfaces that limit the availability of food and nesting resources [8]. Urban development also intensifies thermal stress and exposure to pollution [9, 10]. Additionally, urban infrastructure and roads impede movement between fragmented habitat patches [11, 12]. Despite these hurdles, growing evidence suggests that appropriately managed urban landscapes can support rich insect diversity [13, 14]. For example, urban meadow habitat restorations sustain diverse insect communities [15]. Increasing plant diversity, implementing wildlife-friendly gardening techniques, installing green infrastructure, and increasing tree cover and/or vegetation complexity can also increase insect diversity within urban landscapes [16–18]. Yet, it remains unclear which urban habitat characteristics and management actions are broadly effective over the long-term and across cities that differ in geographic setting, climate, landscape context, and urban form, limiting our ability to make generalizable recommendations for urban biodiversity conservation [8, 19].

Understanding habitat characteristics and management actions that support insect diversity across urban park systems is particularly important. Although insects inhabit many different urban habitat areas, including residential yards and gardens, road margins, and even rooftops [8], parks are managed through coordinated city-wide efforts, meaning that improvements in park management readily scale across urban landscapes [15]. A foundational ecological theory for understanding drivers of diversity within heterogenous systems, such as urban parks, is the Equilibrium Theory of Island Biogeography (ETIB) [20]. According to ETIB, species diversity in parks is controlled by colonization-extinction dynamics, which in turn are hypothesized to be determined by park attributes such as park size and connectivity [20, 21]. Three ETIB hypotheses predict a positive effect of park size on diversity: larger parks support bigger populations that are more likely to persist (population size hypothesis); larger parks provide easier targets for potential colonists (passive-sampling hypothesis); and larger parks contain a greater diversity of resources that could be utilized by a wider range of species with distinct ecological requirements (habitat heterogeneity hypothesis) [21]. ETIB also predicts that park connectivity increases diversity, both by increasing the probability that new species colonize and also by providing a greater supply of potential immigrants that improve the persistence of established populations [21]. Although park size, connec-tivity, and habitat quality have all been linked to insect diversity in urban parks, [20, 22, 23], previous studies have largely focused on static patterns of species richness rather than the colonization-extinction processes that underlie community assembly [20]. This distinc-tion is important because communities experiencing ongoing turnover may exhibit patterns that obscure the mechanisms maintaining diversity. Consequently, studies based solely on snapshot diversity measures may under- or overestimate the importance of park attributes for supporting biodiversity over time [21]. Multi-year analyses are needed to quantify the effects of urban park features on the trajectories of urban ecological communities.

In addition, few studies have compared patterns of insect diversity across many cities, leaving two key knowledge gaps [19]. First, we lack a generalized understanding of how diver-sity in urban parks is influenced by within-city differences (i.e., variation in park attributes) [19]. Identifying park characteristics that consistently benefit insects across urban contexts would improve our ability to make broadly applicable recommendations for biodiversity conservation. Second, cities differ in city-wide management including average park size and connectivity, and/or total area of habitat. However, it is largely unclear how between-city differences impact city-wide diversity at multiple spatial scales, including diversity within-parks (α diversity), among parks (β diversity), and across parks (γ diversity) [19]. This uncertainty limits our ability to identify city-wide management practices that promote biodiversity. For example, while ecologists increasingly recognize the contributions of small habitat patches to landscape scale diversity, many conservation actors prioritize protection of a few larger habitat patches over many, smaller patches of equal cumulative area (i.e., maximizing average park size relative to total habitat area) [24]. Yet, even if larger parks support high α diversity [22], this may only translate into greater city-wide γ diversity when environmental heterogeneity or dispersal creates high species turnover (β diversity) across parks [24–26]. Multi-city studies that simultaneously examine the effect of average park size and total habitat area on γ diversity would clarify the value of prioritizing larger parks within rapidly expanding urban landscapes. Understanding the drivers of diversity across park to city-wide scales is important because these different scales have different implications for ecosystem function and biodiversity conservation goals [25]. For example, while high α diversity can maximize local ecosystem services, high γ diversity can provide spatial and temporal insurance against transient local species losses [25]. Further, to maximize the conservation value of cities, we need to identify how city-wide management impacts species of higher conservation concern, such as disturbance- or edge-avoidant insect species [27].

Previously, multi-city, multi-year urban biodiversity studies have been difficult to accomplish and, to-date, no prior studies to our knowledge have examined how both within-and between-city differences impact insect diversity. However, increasingly widespread availability of data from community science programs, which are especially abundant in urban areas, provide new opportunities to explore ecological questions across these large spatial and temporal scales [5, 28–30]. In this study, we used 5 years of iNaturalist community science data from 2,550 parks across 22 major U.S. cities to test how both within- and between-city differences impact multi-scale spatial and temporal dynamics of urban insect biodiversity (Figure 1). We focused on butterflies because, relative to other insects, their popularity in community science programs has spurred a high availability of species-level detection data [23, 29]. Butterflies are also among the insect groups facing the most severe declines, including in urban areas, underscoring the need to consider them in urban con-servation planning [5, 6]. For all analyses, we used hierarchical Bayesian occupancy models that accounted for imperfect detection of butterflies in the community science data [31, 32]. In part 1, we tested ETIB hypotheses about how park size and connectivity impact initial occupancy of butterflies within urban parks and the change of these communities over time through colonization and/or failure to persist. In part 2, we examined how within-park but-terfly occupancy was additionally impacted by measures of park quality, including tree cover and plant diversity (as a proxy for habitat heterogeneity) within parks as well as the area of woody and herbaceous vegetation cover in the surrounding landscape. Finally, in part 3, controlling for city spatial area and latitude, we tested the importance of between-city differences (median park size, park connectivity, and total habitat area - including habitat both inside and outside of parks and distinguishing between herbaceous vegetation or tree cover habitat types) for determining multi-scale patterns of butterfly diversity, including: (i) median local species richness and the proportion of disturbance or edge-avoidant species in parks (α diversity); (ii) dissimilarity in park community compositions caused by species turnover and nestedness (β diversity); and (iii) the total number of species occurring in each city (γ diversity). By using a robust modeling framework to leverage opportunistic commu-nity science data across previously inaccessible scales, we provide mechanistic insight into how within- and between-city management shapes urban biodiversity, offering generalizable recommendations for biodiversity-friendly urban development.

**Figure 1:**
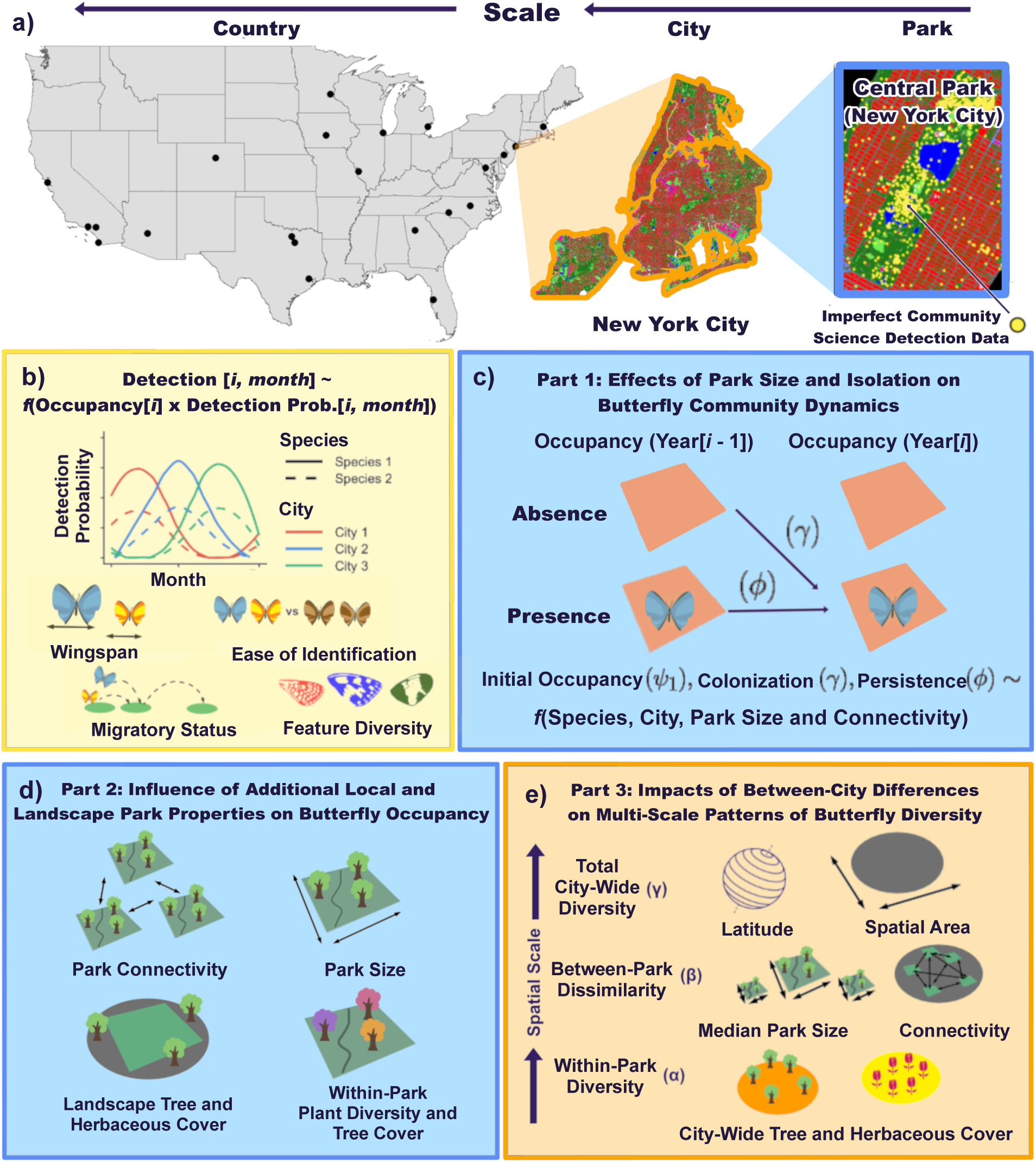
We collected community science detection data for butterflies in urban parks in 22 U.S. cities (a). We used Bayesian occupancy models that allowed butterfly detection probability to vary through time as a function of species and city (b). Using dynamic occupancy models, we examined how park size and connectivity impact colonization and persistence of butterflies at the within-park scale (blue) (c). Using static occupancy models, we considered how additional park properties contributed to butterfly occupancy within parks (blue) (d). We then used our fitted models to reconstruct diversity across all parks at the city-wide scale (orange), and tested the importance of between-city differences on within-, between, and across-park patterns of city-wide butterfly diversity (e).

## 2 Results

### 2.1 Part 1: Effects of Park Size and Connectivity on Butterfly Community Dynamics

Using dynamic multi-species occupancy models, we examined effects of park size and connectivity on initial occupancy, colonization, and persistence of butterflies within parks. We quantified park size as the total vegetated area within a park boundary and connectivity as the area-weighted distance from other parks within a 2 km buffer of the parks.

We found that park size consistently increased initial occupancy probability across all cities (mean = 0.48, 90% BCI = [0.34, 0.61]) (Figure 2a), whereas, on average, park connectivity did not have a strong association with initial occupancy (mean = *−*0.03, 90% BCI = [*−*0.17, 0.09]) (Figures 2a and S1). Overall, colonization rate for the average species in the average park was low (γ[intercept] mean = 0.36% probability, 90% BCI = [0.16%, 0.77%]), meaning that most butterfly species were unlikely to move into new parks where they were not already present. Across cities, park size increased colonization rate (mean = 1.66, 90% BCI = [1.12, 2.18]), while connectivity did not show a clear association with colonization rate (mean = *−*0.33, 90% BCI = [*−*0.77, 0.13]) (Figure 2b and S2). Once species occupied a given park, we found that they had a high rate of persistence (φ[intercept] mean = 98.9% probability, 90% BCI = [97.3%, 99.6%]) (Figure 2c), meaning that species were unlikely to be lost from parks where they were present. Connectivity increased persistence rate (mean = 1.22, 90% BCI = [0.48, 1.79]). Park size did not have a clear effect on persistence (mean = 0.56, 90% BCI = [*−*0.19, 1.41]) (Figures 2c and S3). Butterfly detection rate was significantly impacted by migratory status, feature diversity, and ease of species identification (Figure S4).

**Figure 2:**
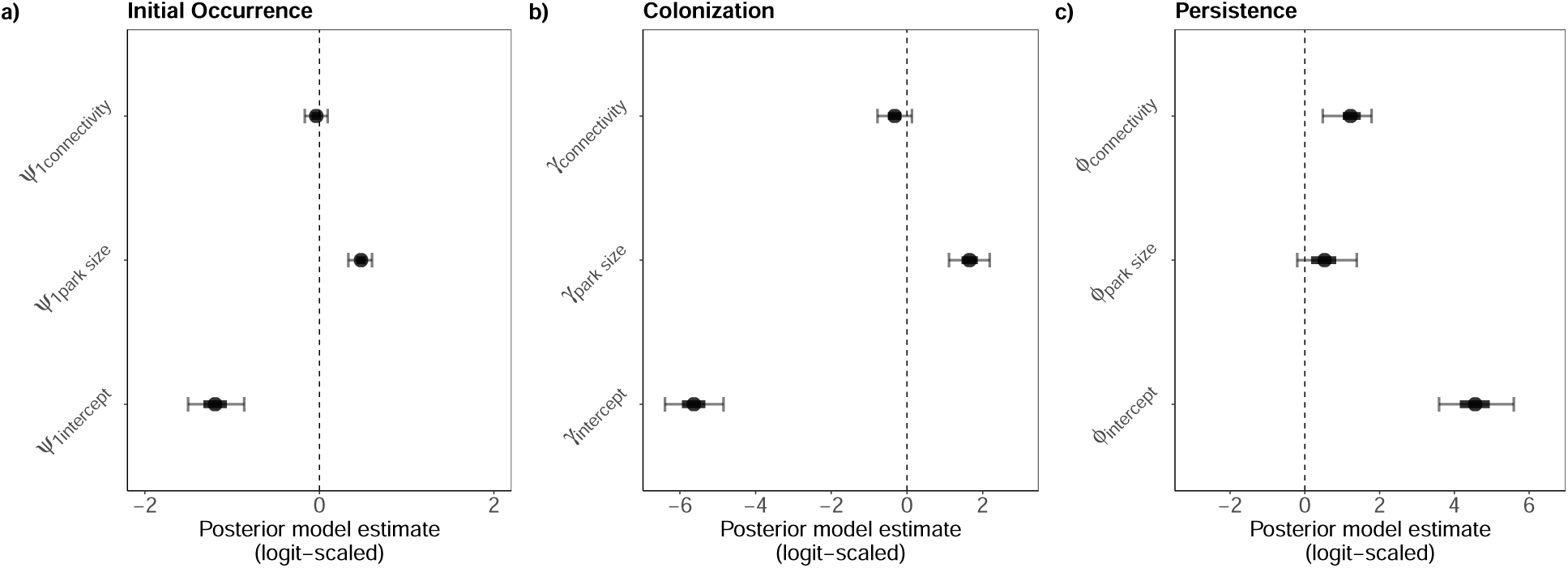
Effects of park size and connectivity on butterfly species initial occupancy (a), colonization (b), and persistence (c). Park size and connectivity were treated as city-specific random effects; posterior estimates for the mean city are displayed. Solid circles indicate posterior means while Bayesian credible intervals (50% (heavy band) and 90% (narrow band)) convey uncertainty in the estimates. Intercept terms (*ψ*1*_intercept_*, *γ_intercept_*, *ϕ_intercept_*) describe the probability of the modeled process for the average butterfly species in the average park in the average city. For park size and connectivity parameters, posterior distributions with a high density above zero suggest a positive effect while a high density below zero suggests a negative effect.

### 2.2 Part 2: Influence of Additional Local and Landscape Park Properties on Butterfly Species Occupancy

Because our dynamic models estimated that overall probability of persistence was high and overall colonization probability was low, we opted to use simplified static occupancy models to test how additional local and landscape park properties contribute to patterns of butterfly occupancy (Figure S5). We used iNaturalist detection data to quantify the diversity of flowering plant genera within parks [23], UrbanWatch’s 1 meter resolution remote sensing data to calculate the proportion of tree cover within parks [33], and NLCD 30 meter resolution remote sensing data to calculate the total area of herbaceous and woody vegetation within a 2 km buffer of the parks [34].

After incorporating these additional predictor variables (Figures S6, S7 and S8), we found that park size had the strongest direct effect on butterfly species occupancy (mean = 0.36, 90% BCI = [0.21, 0.49]) (Figure 3a). Increasing park size from one standard deviation below average to one standard deviation above average is expected to increase occupancy probability from *∼* 13% (90% BCI = [11.9%, 15.0%]) to *∼* 24% (90% BCI = [21.4%, 26.3%]) (Figure 3a). The effect of park size was strongest in San Francisco, New York City, Denton and Chicago, and weaker in San Diego, Riverside, Minneapolis, and Denver (Figures 3c and S10). Plant diversity within parks also increased the probability of butterfly occupancy (mean = 0.30, 90% BCI = [0.22, 0.39]) (Figure 3a). We found that park size had a positive association with plant diversity (Figure S9). To distinguish the direct and total effects of park size on butterfly occupancy (Figure S5), we fit a second occupancy model that did not include plant diversity. Omitting plant diversity further increased the effect of park size (Figure S11), indicating that park size has both a positive direct effect on butterfly occupancy as well as an additional positive indirect effect that is mediated by the positive impact on plant diversity [35].

**Figure 3:**
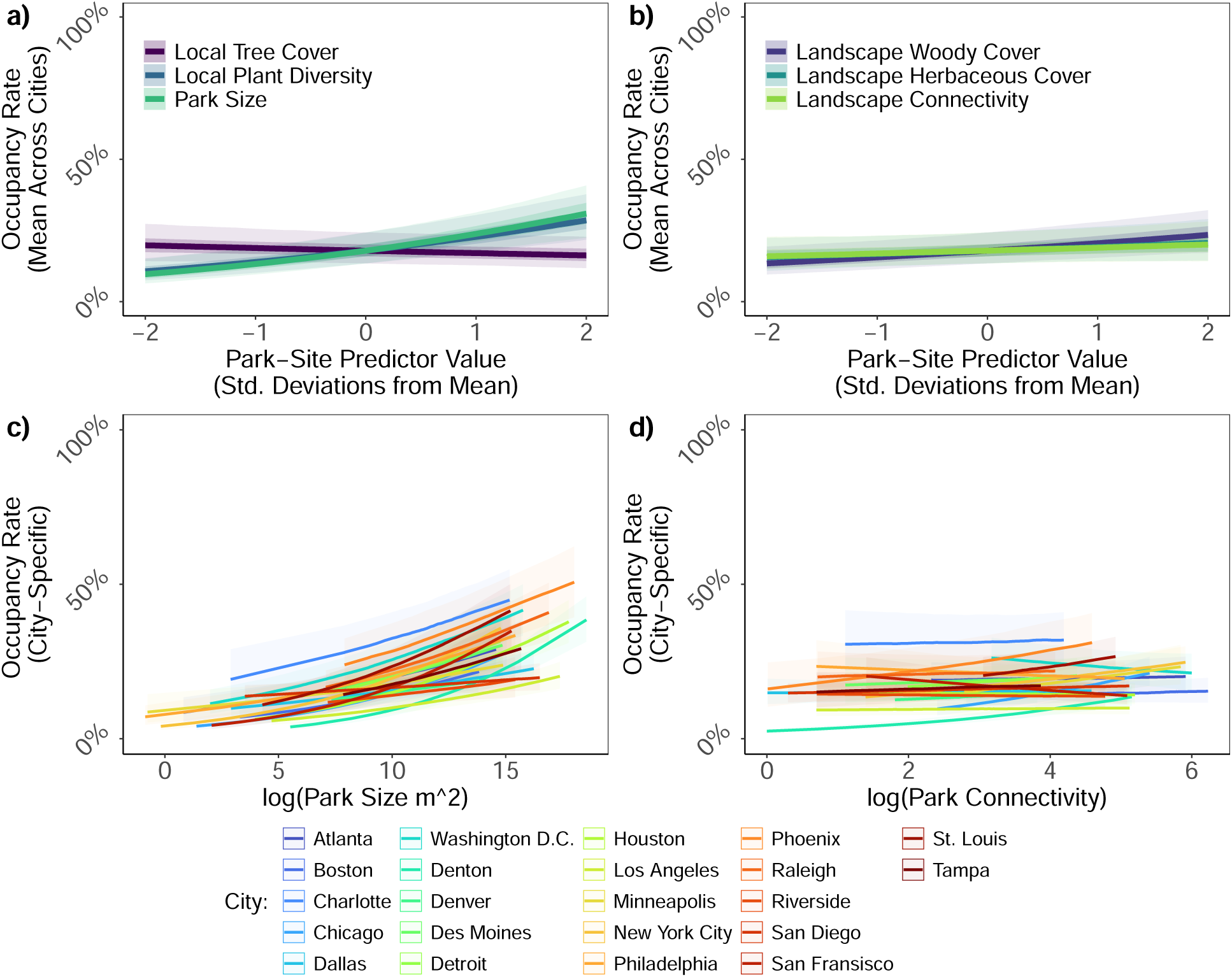
Predictions of within-park butterfly occupancy rates across a range of park-site predictor values. Top panels show the expected change in occupancy rate for the average species in the average city across the observed range of (a) local and (b) landscape park properties. Bottom panels show variation in the expected change in occupancy rate across the (c) park size and (d) connectivity values that were observed in each city while accounting for city-specific variation in occupancy rate intercepts and park-site predictor effects. 50% and 90% Bayesian credible intervals illustrate uncertainty in predictions for the average city (panels a and b). For visual simplicity, city-specific predictions include only 50% Bayesian credible intervals (panels c and d). City-specific predictions are plotted across the range of predictor values that were used to fit the occupancy models (i.e., the range of size and connectivity of parks with one or more butterfly community samples).

Across cities, area and distance weighted connectivity to other parks within a 2km landscape buffer did not have a clear effect on butterfly species occupancy (mean = 0.07, 90% BCI = [-0.04, 0.18]) (Figure 3b). However, the effect of connectivity varied among cities (Figure 3d): For example, connectivity had a more positive impact on butterfly occupancy in Denton, Chicago, Phoenix, and St. Louis versus a more neutral effect in San Francisco, Washington D.C., Philadelphia and Houston (Figures 3d and S12). Within the same 2km landscape buffer of the parks, the area of woody plant vegetation consistently increased occupancy (mean = 0.17, 90% BCI = [0.06, 0.26]) (Figure 3b). In contrast, the total area of herbaceous plant cover in the surrounding landscape did not have a clear impact on occupancy (mean = 0.08, 90% BCI = [-0.04, 0.22]) (Figure 3b).

### 2.3 Part 3: Impacts of Between-City Differences on Multi-scale Patterns of Butterfly Diversity

Cities showed differences in park and landscape features, such as the distribution of park sizes or the total area of vegetation spread across the city (Figures S13, S14) and in how these features impact butterfly species occupancy (Figure 3c and 3d). To understand how these between-city differences impact patterns of α, β, and γ diversity, we used our fitted occupancy model from part 2 to reconstruct butterfly species occupancy in all parks in each city (Figure S15). We then calculated local to landscape scale metrics of city-wide diversity and quantified relationships between diversity and city-wide features including median park size, city-wide park connectivity, and city-wide area of tree or herbaceous vegetation cover (Figures S16 and S17). We accounted for latitude as a predictor of α diversity and γ diversity measures, and total city area as a predictor of γ diversity. Given the limited sample size of 22 independent cities, our final models included only the predictors that showed at least marginal evidence of an association with the diversity outcome (i.e., at least the 50% BCI departing from 0).

Median within-park species richness (α diversity) for the average city was *∼* 23 species (Intercept mean = 23.5, 90% BCI = [20.9, 26.4]) (Figure 4a). After accounting for latitude (Figure S18), median park size was the strongest driver of within park diversity (mean = 2.77, 90% BCI = [0.1, 5.7]) (Figure 4a). Increasing city-wide median park size from one standard deviation below average (*∼* area of 0.5 soccer fields) to one standard deviation above average (*∼* area of 3 soccer fields) increased median local species richness by *∼* 5 species (mean = 5.4, 90% BCI = [0.15, 11.2]) (Figure S19). City-wide tree cover showed a weak association with within park diversity (mean = 2.11, 90% BCI = [*−*0.56, 4.7]) (Figure 4b). On average, urban parks contained *∼* 2 disturbance- or edge-avoidant species (Intercept mean = 0.78 (log-scale), 90% BCI = [0.52, 1.05]), but this was positively impacted by city median park size (mean = 0.27 (log-scale), 90% BCI = [0.02, 0.51]) (Figure S20).

**Figure 4:**
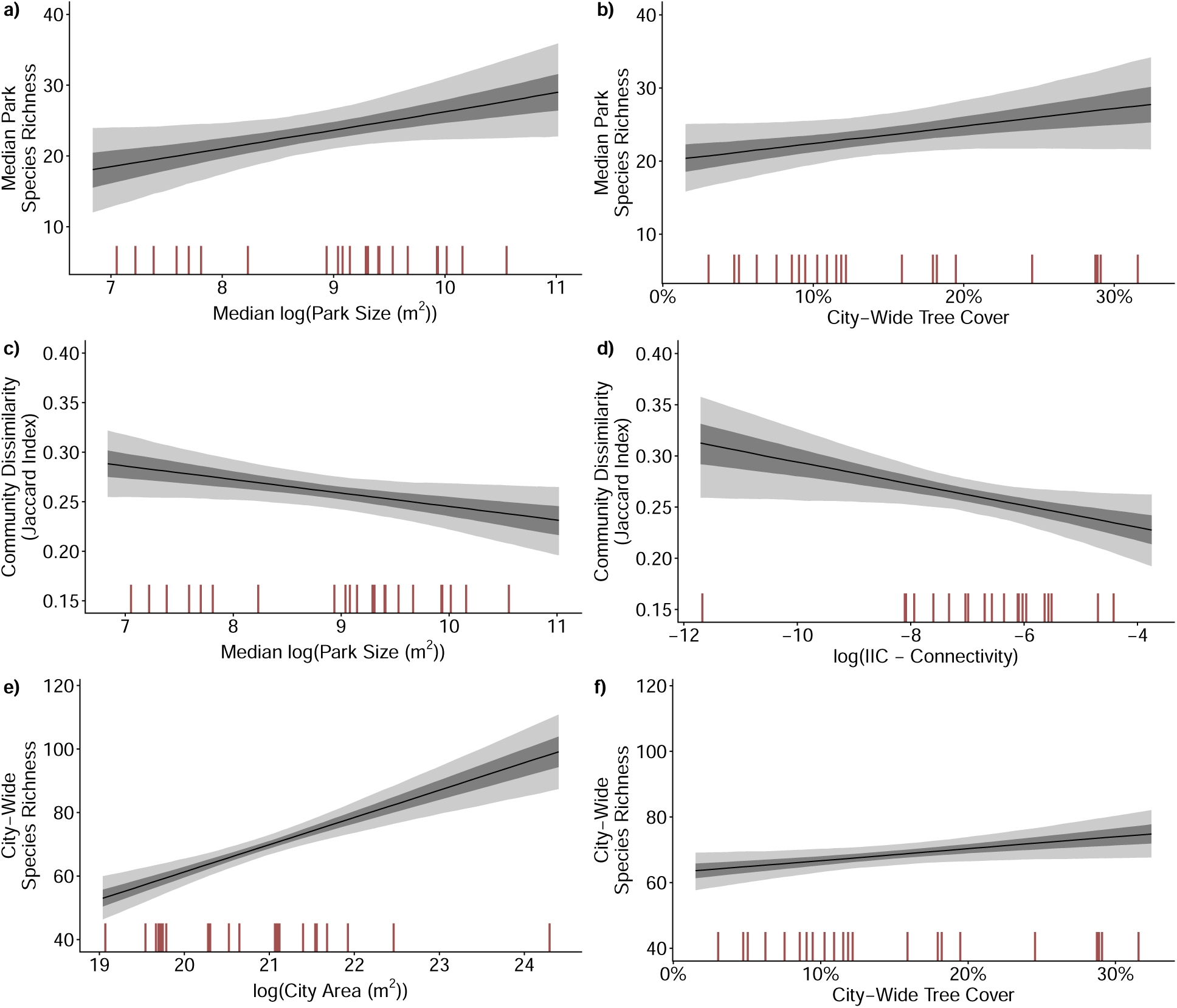
Impacts of between-city differences on butterfly diversity. Using the entire posterior distribution from our fitted occupancy model, we simulated species occupancy in all parks in each city and summarized metrics of city-wide diversity including: Median within-park species richness (a, b), Jaccard index of community dissimilarity (c, d), and total city-wide species richness (e, f). Using GLMs, we quantified effects of between-city differences on these diversity outcomes. These counterfactual prediction plots show the expected relationship for the two predictors with the largest estimated effect size. Light and dark confidence bands illustrate 50 and 90% Bayesian credible intervals around the mean expected relationship.

Average dissimilarity in species composition (Jaccard index of β diversity) (Intercept mean = 0.26, 90% BCI = [0.24, 0.28]) decreased with park size (mean = -0.01, 90% BCI = [-0.03, 0.00]) (Figures 4c and S21). We decomposed the contributions of species nestedness versus species turnover to dissimilarity, and found that park size reduced dissimilarity by decreasing species nestedness, meaning that cities with smaller parks contained many communities that were subsets of the species composition in the most diverse parks (Figure S22). City-wide park connectivity also decreased community dissimilarity (mean = -0.01, 90% BCI = [-0.03, 0.00]) (Figure 4d). However, this was driven by a negative effect on species turnover rather than nestedness, meaning that species were distributed across a greater proportion of park sites in cities with more connected park systems (Figure S23). City-wide tree cover reduced community dissimilarity (mean = -0.01, 90% BCI = [-0.02, 0.00]) (Figure S20) by decreasing nestedness (Figure S22).

Total species richness (γ diversity) for the average city was around 68 species (Intercept mean = 68.4, 90% BCI = [65.3, 71.4]) (Figure 4e). The total spatial area of a city had the strongest effect on γ diversity (mean = 10.3, 90% BCI = [6.4, 14.1]) (Figure 4e). Latitude had a weak negative association with γ diversity (mean = *−*1.8, 90% BCI = [*−*5.7, 2.2]) (Figure S24). After accounting for total area and latitude, city-wide tree cover increased γ diversity (mean = 3.24, 90% BCI = [0.1, 6.4]) (Figure 4f). Increasing city-wide tree cover from below average (6%) to above average (22%) was expected to increase total species richness by more than *∼* 6 species (mean = 6.5, 90% BCI = [0.1, 12.7]) (Figure S25).

## 3 Discussion

Synthesizing community science data with a robust occupancy modeling framework allowed us to conduct one of the largest investigations of urban biodiversity patterns to date. Looking across 22 major U.S. cities, our study had three key findings: 1) Larger parks consistently supported greater butterfly diversity by promoting species colonization. In contrast, the effects of park connectivity on butterfly communities varied across cities; 2) Flowering plant diversity within parks and woody vegetation surrounding parks consistently increased urban park butterfly occupancy; and 3) Increasing city-wide tree cover increased the total number of butterfly species found across a city’s entire network of parks. Together, these findings demonstrate that both within- and between-city differences in urban management contribute to the spatial and temporal dynamics of urban ecological communities. In application, our results emphasize that urban planners and policy makers should maintain or increase the size of individual parks to promote high levels of local biodiversity. Simultaneously, increasing plant diversity will improve the habitat quality of urban parks for butterflies. However, for promoting biodiversity at the city-wide scale, which is crucial for supporting longer-term biodiversity conservation and stable ecosystem functioning, our results suggest that increasing total city-wide habitat area may be more important than optimizing the size or connectivity of individual parks.

We found that cities with bigger parks support more diverse insect communities at local scales: Increasing city-wide median park size from an area of approximately 0.5 soccer fields to 3 soccer fields was estimated to increase butterfly species richness in the average park by *∼* 30%. Parks with high α diversity may be especially important in cities, where they provide an increased sense of engagement with nature that can benefit park users [2]. Increasing median park size also increased the average number of disturbance- or edge-avoidant species in urban park communities. Since these types of species are facing steeper declines and are often excluded by the ecological filters imposed by urbanization, these results highlight that integrating larger parks within urban landscapes is crucial for supporting species of higher conservation concern and boosting functional diversity [26, 27]. We also found that nestedness of butterfly communities was higher in cities with smaller parks. This suggests that as average park size declines, the most species rich parks (i.e., the largest parks in a city) become increasingly important for supporting city-wide γ diversity [36]. Following ETIB predictions, and similar to studies of butterflies in habitat patches embedded within mixed or agricultural mosaic landscapes [37, 38], our dynamic occupancy models suggested that larger parks primarily increase butterfly species richness by increasing colonization, supporting the ETIB passive-sampling hypothesis. In contrast, the effect of park size on species persistence was weak. These findings suggest that interventions that facilitate species recruitment could help offset lower levels of diversity in smaller parks, for example, ensuring the presence of host plants that encourage colonization of targeted species [39].

Our results suggested that plant diversity positively mediated the effect of park size on butterfly occupancy, supporting the ETIB habitat heterogeneity hypothesis. This means that bigger parks tend to have higher plant diversity, but also that having higher plant diversity increases butterfly occupancy regardless of park size. Supporting our results, Lilkendey et al. (2025) found that flowering plant diversity (especially native plant diversity) across urban green spaces in Florida positively contributed to species richness, including for other beneficial insects such as wild bees [23]. These findings are important from a management perspective because in urban landscapes that are already heavily developed, increasing plant diversity (e.g., by adding flowering meadow enhancements to turfgrass dominated spaces) may be a more feasible and immediate intervention strategy compared to expanding the spatial area of parks [15].

Unexpectedly, we did not find consistent effects of park connectivity on butterfly diversity. Instead, effects of connectivity varied among cities. For example, connectivity increased butterfly occupancy in New York City and Chicago, but had neutral to negative effects in San Francisco and Houston. This variation may be due to differences in city topology [40]. Clear positive impacts of connectivity could have occurred in cities with more inhospitable or impermeable land cover between parks, including denser infrastructure or taller buildings that may increase the cost of moving between park spaces [12]. Connectivity also had a more positive effect in Denton, the city with the greatest proportion of agricultural land cover between parks. Common agricultural practices that reduce health and survival of butterflies (e.g., pesticide use) may have increased the benefits of connectivity by amplifying the cost of traversing between parks [41]. Consistent with release from dispersal limitation [36], city-wide connectivity was negatively associated with species turnover. Importantly, reduced turnover could lead to lower city-wide (γ) diversity unless within-park diversity remains high [36]. Our dynamic occupancy model suggested that connectivity contributed to patterns of butterfly occupancy by increasing persistence rather than colonization. Proximity to other parks may not have increased colonization if unoccupied parks did not meet species ecological requirements, whereas the positive effect on persistence is consistent with immigration driven population rescue effects that buffer small populations that are already established against environmental or demographic stochasticity [42]. Given the inconsistent effects of connectivity that we observed, future research should examine how differences in city topologies impact insect movement and survival.

After controlling for other predictors including average park size and connectivity as well as intrinsic city factors of city area and latitude, we found that city-wide tree canopy cover (including canopy cover within and outside of park boundaries) was the most important predictor of γ diversity, increasing city-wide species richness by *∼* 10% in cities with above average versus below average canopy cover. This positive effect on city-wide species richness is critical because high γ diversity can provide spatial and temporal insurance against transient local species losses, translating into a city’s overall potential to support biodiversity conservation and stable ecosystem functioning [25]. Given that the total amount of habitat in an urban landscape was a more important driver of city-wide species richness than the size or connectivity of individual habitat patches, our findings suggest that a few large or many small parks (i.e., SLOSS framework dichotomies) will yield similar outcomes for urban γ diversity [24, 26]. This supports increasing recognition that preserving many small habitat patches in rapidly expanding anthropogenic landscapes is critical for advancing global biodiversity frameworks [24, 26], for example, for achieving 30 x 30 targets. In urban landscapes, small habitat patches including those embedded in private gardens or neighborhood boulevards have the potential to make meaningful contributions to a city’s overall capacity to support biodiversity [43, 44]. Further, this result highlights the value of city-wide canopy cover initiatives, for example, where the City of Montreal (Canada) recently committed to planting 500,000 new trees between 2025 and 2030 [45].

Our analyses suggested that tree cover surrounding a park (but not the proportion of tree cover within a park) drove increases in park occupancy. Complementarity in habitat use across life history stages could underlie this disconnect: Most of our detections came from adult butterflies, which are often more active in open habitat spaces [46]. Yet, woody plants surrounding parks and/or understory plants in these habitats likely provide important resources for butterfly larvae [44]. Landscape canopy cover may have also increased functional connectivity to other park spaces [47]. In addition, landscape canopy cover moderates temperature fluctuations, reducing thermal stress for butterflies in parks while simultaneously cooling cities as a whole, supporting additional benefits to human health [48]. In contrast to tree cover, we did not find consistent effects of landscape herbaceous cover on butterfly occupancy. This indicates that, on average, urban herbaceous habitat areas currently do not make strong contributions to the health of urban park butterfly populations, likely because herbaceous habitats in urban areas are often dominated by manicured turfgrass lawns with low plant diversity and frequent disturbances caused by mowing and pesticide use [15, 49]. The stronger importance of woody versus herbaceous vegetation for city-wide diversity suggests that wildlife friendly urban gardening schemes or habitat restorations should focus on integrating more trees and shrubs into planting mixes and/or improve the ways that herbaceous habitats are currently managed.

While we found that average park size predicts city-wide local diversity and that city-wide tree cover is associated with city-wide total diversity, few existing studies have examined how between-city differences impact biodiversity. For birds and mammals, intrinsic drivers such as city spatial area or city latitude were important determinants of city-wide diversity [40, 50, 51], which we also observed here for butterflies. For mammals, city-wide management impacted urban ecological communities on a local scale: local urbanization levels decreased mammal diversity, and this effect was stronger in cities with lower total vegetation cover [40]. Yet, in contrast to our results, previous studies did not find strong evidence that aspects of city-wide management impact city-wide patterns of diversity [40, 51]. We may have detected a stronger effect of city-wide tree cover on butterfly diversity given the finer resolution of our data: Our high resolution (1 meter) land cover data captured variation in tree cover contributed by small canopy patch areas, such as private yards and gardens. Alternatively, city-wide habitat quality may have had less impact on birds and mammals if urban bird and mammal communities are dominated by urban exploiter species that are less likely to respond to small scale changes in the quality or amount of semi-natural habitats [52]. The responsiveness of insects to city-wide management reinforces that urban insect conservation efforts are an especially important pathway for advancing urban sustainability. Overall, our findings highlight that city-wide management decisions, and not just intrinsic differences such as city size or geographic location, can contribute to a city’s capacity to support biodiversity conservation, especially for insects.

Reaching global Sustainable Development Goals requires promoting biodiversity in and around cities [3]. Creating urban landscapes with rich diversity of butterflies and other species groups will improve opportunities for people to interact with nature [2]. Enhancing these opportunities for human-nature interactions and nature connectedness can then strengthen support for conservation policy and initiatives [53]. Increasingly, urban communities engage with neighborhood biodiversity through community science programs [23, 28, 30, 53]. In this study, we developed a scalable methodological approach that leverages the data from community science programs to inform conservation management approaches that promote urban biodiversity, which in turn may drive positive feedbacks by reinforcing the benefits of community science participation [2, 53]. As such, beyond examining the drivers of urban biodiversity across unprecedented spatial and temporal extents, this study helps to close the loop between community science participation and conservation practice.

## 4 Materials and Methods

### 4.1 Data Collection

The UrbanWatch dataset [33] contains 1 meter resolution land cover data for 24 cities across the United States. We excluded two of these cities from our analyses: Seattle, because it did not have detections of multiple species of butterflies from any family which was necessary for our method of inferring background sampling effort; and Miami, because it had the least amount of butterfly detection data of any remaining city while also being the only city in its ecoregion area which precluded us from sharing information about species random effects from other cities (see Statistical Analyses Part 1). The remaining 22 cities included: Atlanta, Boston, Chicago, Charlotte, Washington D.C., Dallas, Denton (County), Denver, Des Moines, Detroit, Houston, Los Angeles, Minneapolis, New York City, Philadelphia, Phoenix, Raleigh, Riverside, San Diego, San Francisco, St. Louis, and Tampa. We used the National Land Trust ParkServe shapefile [54] to identify parks within the boundaries of each city, which were defined by UrbanWatch [33]. We included parks that straddled city boundaries in our analyses.

Next, we derived metrics about local and landscape park qualities that we hypothesized would affect patterns of butterfly species occupancy (Figures S13 and S14). Using UrbanWatch data, we quantified park size as log(area of herbaceous and tree cover (m^2^)) of each park [33]. We quantified plant diversity as the number of flowering plant genera within a park detected by iNaturalist users across 2020 to 2024 [55]. We grouped plant detections at the genus level because closely related plant species may provide similar functional value to butterflies and because we were more confident that genus level identification would be accurate and less affected by potential detection biases. We calculated within-park proportion of tree cover using UrbanWatch data [33].

We quantified park-level landscape connectivity as the area weighted distance to each neighboring park *n* summed across all neighbors *N* within a maximum distance of 2 km of the park boundary [56]: 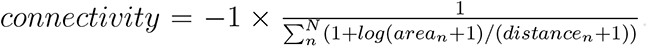.Increasing the area of neighboring parks and/or decreasing the distance of the neighboring parks increases the value of this metric. We chose a threshold of 2km based on plausible maximum inter-park movement potential for the average butterfly species [57]. We allowed parks from outside of a city’s boundary to contribute to the connectivity of parks that were located along or near a city edge. Using NLCD 30 meter resolution land cover data, we also calculated the proportion of herbaceous vegetation cover and woody (shrub or tree) vegetation cover within the 2 km buffer around each park [34]. Using this data (rather than the finer scale UrbanWatch data) allowed us to capture vegetation cover surrounding park sites that were located along or near a city edge.

We used GBIF to collect research grade detections of butterflies (Nymphalidae, Pieridae, Papilionidae, Hesperiidae, and Lycaenidae/Riodinidae families) from iNaturalist [58]. We only included data from 2020 to 2024 because this period had a high number of observations. We filtered out observations with *>* 100 m coordinate uncertainty. This allowed us to be confident that detections were actually observed within the parks to which they were assigned, in contrast to geoprivacy obscured detections or detections with low or unknown coordinate precision. We assigned species detections to park sites if the detection occurred within the park or within a 50 m buffer of the park site. Using this detection buffer, some individual detections were attributed to multiple parks if the parks were close enough that the detection was within an area of park buffer overlap. We only modeled species that were detected at least once in any park in any city. In addition, for each species, we only modeled occupancy dynamics for cities within the species geographical range [32]. We considered a city “in-range” if the species was detected at least once anywhere within the city or within a 20 km buffer area around the city. After applying our filters, our data consisted of 23, 416 unique (species by park by month by year) butterfly detections across 2, 550 park sites. Our analyses included 183 species, with a median of 21 unique detections per species (Supplemental Material: Species Detections Table).

## 4.2 Statistical Analyses

### 4.2.1 Statistical Analyses - Part 1: Effects of Park Size and Connectivity on Butterfly Community Dynamics

We used dynamic multi-species occupancy models to determine how park size and connectivity affect urban butterfly communities [31]. These models estimate initial occupancy of species (whether species were present or absent in parks in the first year they were sampled) and how occupancy changed over time through colonization (appearance of species into previously unoccupied parks) or failed persistence (loss of species at previously occupied parks). Simultaneously, these models explicitly estimate and account for imperfect detection (the failure to detect species that are present) [31]. For this analysis, we only included park sites that had one or more community sampling events in two or more years, which was necessary for us to estimate inter-annual temporal dynamics. The reduced dataset for the dynamic occupancy model contained 20, 550 unique detections of butterflies across 1, 178 of the park sites.

To apply these models to opportunistic community science data, we first converted our butterfly detections into binary outcomes. If a species was detected at a park site one or more times during a particular month, then we assigned a positive detection for that month (*y*[*i,* month] = 1). Following [32], we assigned non-detections by inferring community sampling events: If other species from the same butterfly family were observed at the same site during that month, then we assigned a non-detection (*y*[*i,* month] = 0); but if no other species from the same family were reported, we instead treated the data as NA’s (*y*[*i,* month] = NA)) (Table S1). We constrained our inference of non-detections by family because the taxonomic interest of some iNaturalist contributors may have been limited in scope [59]. Our approach of distinguishing between non-detections versus a lack of survey effort allowed us to account for spatial and temporal variation in sampling effort.

Under the dynamic occupancy model framework [31], we let *Z* represent an imperfectly detected occupancy state (presence or absence) for each observed park site, species, and year combination (*i*). We assumed that occupancy was Bernoulli distributed: *Z*[*i*] *∼* Bernoulli(*ψ*[*i*]). For the first year that a site was sampled for a specific butterfly family, we treated *ψ*[*i*] as equivalent to an initial occupancy probability: If year *i* = 1, then *ψ*[*i*] = *ψ*_1_[*i*]. To estimate rates of persistence (*ϕ*[*i*]) and colonization (*γ*[*i*]), we considered *ψ*[*i*] for latter years as the outcome of *ψ*[*i*] in the most recent previous year that received some survey effort for the same butterfly family multiplied by the corresponding rate: if year[*i*] *>* 1, then *ψ*[*i*] = *ϕ*[*i*] *× ψ*[year[i] *−* 1] + *γ*[*i*] *×* (1 *− ψ*[year[i] *−* 1]). With this formulation, our model heavily weighs the likelihood of persistence (*ϕ*) if a species was more likely to have occurred in the previous year (*ψ*[year[i] *−* 1] approaching 1); alternatively, our model heavily weighs the likelihood of colonization (*γ*) if a species was less likely to have occurred in the previous year (*ψ*[year[i] *−* 1] approaching 0). We then modeled hypothesized drivers of initial occurrence, colonization, and persistence as:

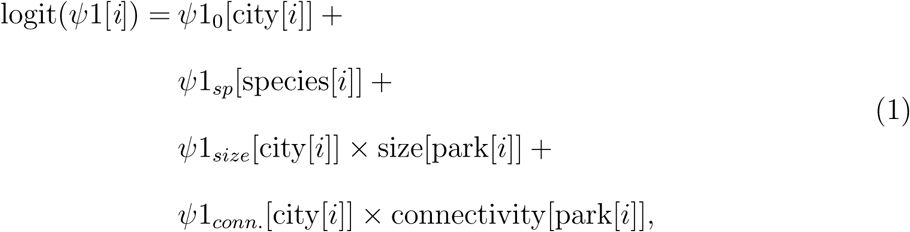

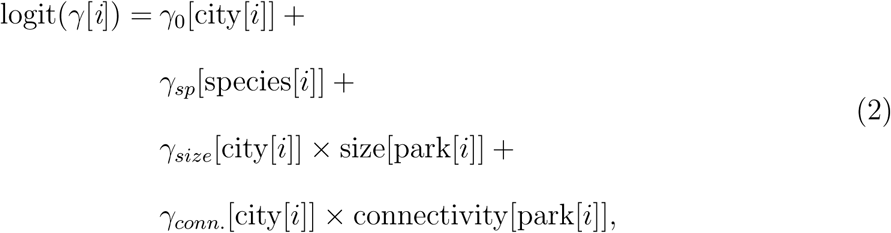

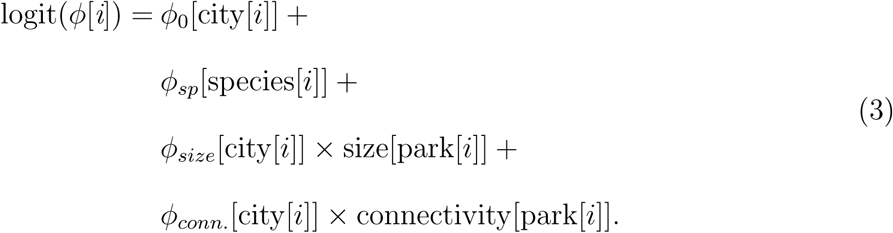

In equations 1, 2, and 3, *ψ*1_0_, *γ*_0_, and *ϕ*_0_ represent city-specific random intercepts. Equations 1, 2, and 3 included city-specific random effects of park size and connectivity (e.g., *ψ*1*_size_* and *ψ*1*_conn._*), which allowed us to test whether bigger and more connected parks support more species. *ψ*1*_sp_*, *γ_sp_*, and *ϕ_sp_* represent species random effects on the intercept. We allowed the mean of the species random effects to vary by species wingspan and migratory status, which we expected would both positively impact occupancy by increasing the ability for species to access more sites across urban landscapes [57]. The species random effects were centered on zero and therefore represent deviations from the city-specific intercept effect. For example, for initial occupancy:

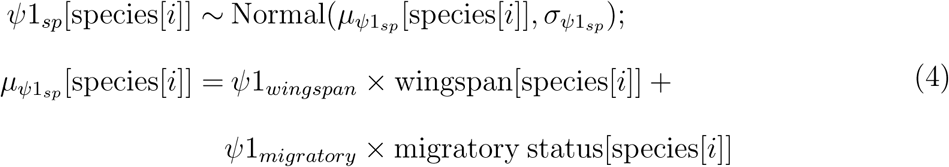

To account for imperfect detection, we assumed that occupancy of each species at each site is fixed within each year [31]. We then assumed that detection during each month was the outcome of a Bernoulli trial conditional on both occupancy and a month-specific detection rate, *p*[*i,* month]: *y*[*i,* month] *∼ Bernoulli*(*Z*[*i*]*×p*[*i,* month]) [31]. We then modeled heterogeneity in detection rate as:

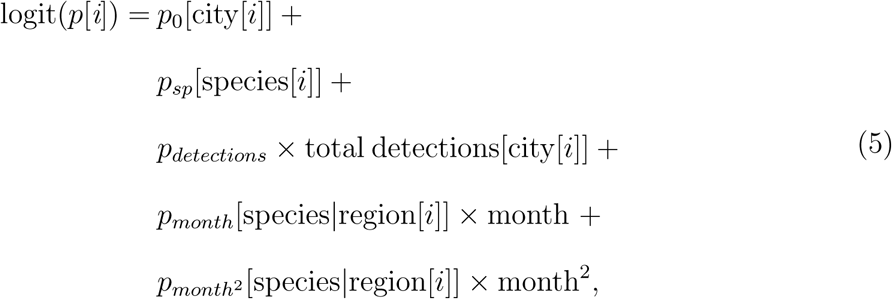

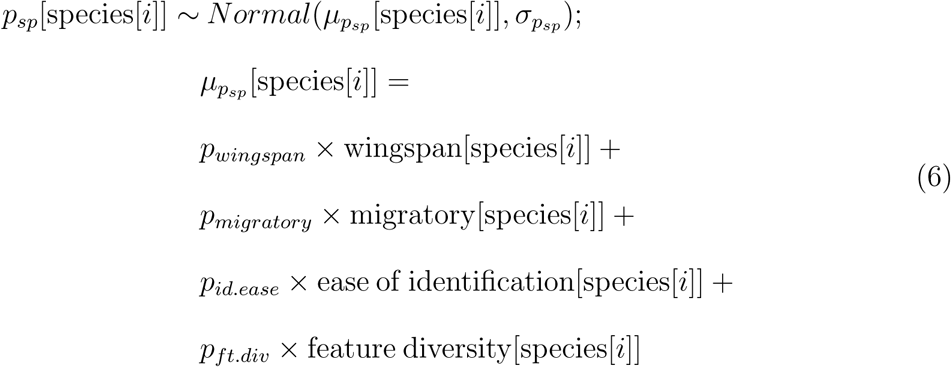

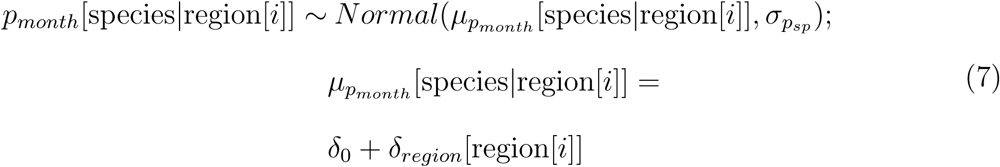

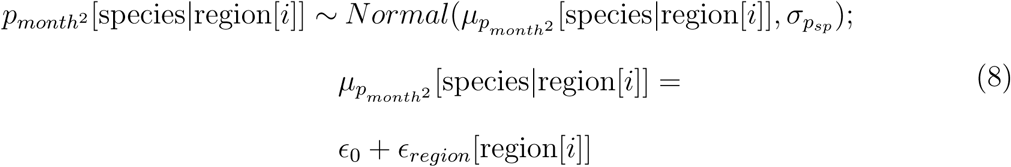

We included an effect of the total number of butterfly detections per city (*p_detections_*) to account for city differences in overall iNaturalist participation. We included species traits that we hypothesized would impact detection rate as predictors of species random intercept effects, *p_sp_*. Based on previous studies, we expected that detection would be positively impacted by: species size (wingspan), the number of identifiable features (e.g, tail, eyespot, checker patterns), and how easy a species is to identify [59]. We also expected that migratory species would be more likely to be detected given that many of them are highly charismatic and are popular focal points in monitoring programs (e.g., Monarch, Red Admiral, Gulf Fritillary, Buckeye). We used publicly shared data from [59] to assign butterfly wingspan and feature diversity. For species not already included in that study, we added the species trait data using the same methodology: we calculated wingspan as the average length of the forewing and hindwing as listed by LepTraits v1.0 [60], and we calculated feature diversity by manually parsing the species descriptions from the Butterflies and Moths of North America database [61]. As a proxy for ease of identification (calculated by genus), we divided the number of research grade iNaturalist detections by the total number of iNaturalist detections within in the United States [59]. We obtained migratory status from a previous synthesis [62], and assumed that species with no evidence of migration were non-migratory.

We treated species random effects for initial occupancy as distinct (statistically unrelated) within each city. However, for colonization, persistence, and detection we assigned the same random effect for all species across all cities within a regional cluster. We identified regional clusters based on Level 1 U.S. EPA Ecoregion groupings and city latitude: North-east (Boston, NYC, Washington D.C., and Philadelphia), Midwest (Chicago, Detroit, St. Louis, Minneapolis), Central Plains (Denver and Des Moines), Southeast (Atlanta, Charlotte, Raleigh, and Tampa), Texas (Dallas, Denton, and Houston), Southern California (San Diego, Los Angeles, and Riverside), Northern California (San Francisco), and Interior South-west (Phoenix). Grouping species random effects by regional cluster allowed us to reduce the number of parameters in our model (and increase the amount of data informing each of those parameters) while retaining flexibility in species ecology and community science detection processes across the continental spatial extent of our study.

Finally, as in [15], we included random effects of month on detection rate to capture phenology driven detection heterogeneity: *p_month_* describes the month with the highest probability of detection, where more positive values indicate a later annual phenological peak; and *p_month_*_2_ describes the shape of the annual phenology, where more negative values indicate more attenuation in detectability around the peak month. Here, including effects of region (*δ_region_* and *ɛ_region_*) allowed the community mean peak and attenuation strength to vary by region.

### 4.2.2 Statistical Analyses - Part 2: Influence of Additional Local and Landscape Park Properties on Butterfly Species Occupancy

We used static multi-species occupancy models to test hypotheses about how additional park properties contribute to butterfly occupancy. The static occupancy models included a single ecological submodel that describes the probability of occupancy of a species at site in a year. This simpler structure allowed us to include additional city-specific random effects of park properties. For this model, we included all parks with at least one community sampling event.

Using a causal inference approach, we diagrammed directed acyclic graphs (DAGs) that outlined our hypotheses about the drivers of butterfly occupancy in parks (Figure S5). We fit two static occupancy models to determine both the direct and total causal effects of some park properties. With the first model (*m*2.1), we quantified the direct effects of park size, local plant genera diversity, local tree cover, landscape connectivity, landscape herbaceous vegetation cover, and landscape woody vegetation cover on occupancy (*ψ*):

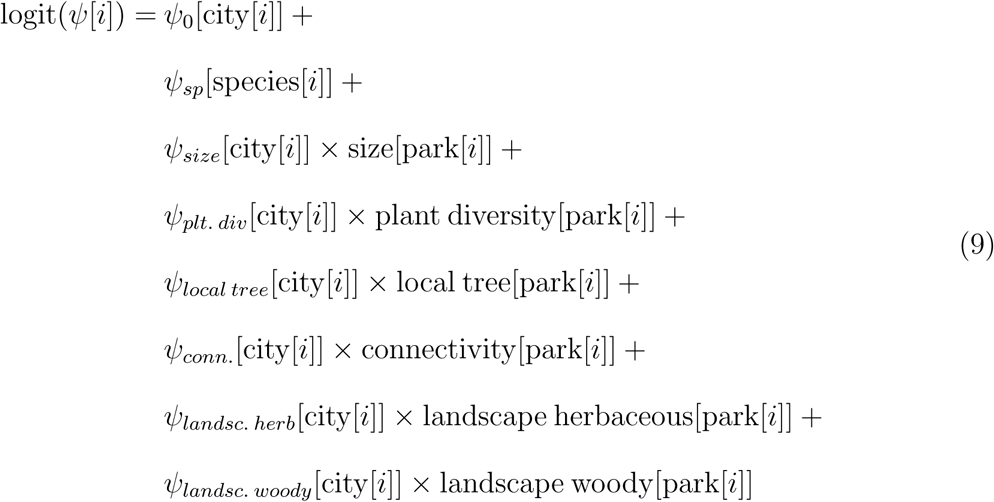

Following our DAG (Figure S5), we fit a reduced model (*m*2.2) that allowed us to quantify the total effects of park size, local tree cover, and connectivity on butterfly occupancy [35]. We hypothesized that both park size and local tree cover would indirectly effect occupancy by increasing plant diversity. We also hypothesized that landscape connectivity would indirectly effect occupancy by increasing landscape herbaceous and woody vegetation. After removing mediating effects of plant diversity and landscape herbaceous and woody vegetation, an increase in the effect of the retained predictors would provide evidence that they have additional indirect effects on butterfly occupancy [35]. To account for detection biases, both *m*2.1 and *m*2.2 included the same detection submodel outlined in Part 1.

We used a Bayesian approach to fit the occupancy models, with models written in Stan, implemented in R with cmdstanr [63]. We employed best practices for model development and fitting by: using weakly-informative priors to discourage unrealistic parameter values (Figures S26 and S27); confirming sufficient mixing of chains (Gelman-Rubin R-hat values *<* 1.1) (Figures S28 and S29), minimal within-chain autocorrelation (effective sample size/steps greater than 0.1) and no divergent transitions; and running each chain for 500 warmup steps followed by 1000 sampling steps (Figures S30 and S31) [63, 64]. We z-score standardized continuous covariates for model fitting. We assessed model fit using visual posterior predictive checks that compared real number of observed detections in each city to model-based predictions of the number of detections in each city (Figures S32 and S33).

### 4.2.3 Statistical Analyses - Part 3: Impacts of Between-City Differences on Multi-scale Patterns of Butterfly Diversity

We used our fitted occupancy model from Part 2 to reconstruct patterns of species occupancy in all parks in each of the twenty-two cities, sampling a set of 200 parameter values from the posterior distribution of *m*2.1. For each sample from the posterior distribution, we simulated species occupancy across all parks (within species geographic range) as a random draw from a Bernoulli distribution where the expected value was a linear combination of the sampled parameter values and the park and species covariates. This included simulating occupancy at parks that did not receive any sampling effort. Next, we used the reconstructed patterns of species occupancy to calculate 4 metrics of city-wide biodiversity: (1) median number of species in each park; (2) mean proportion of disturbance- or edge-avoidant species in each park; (3) mean Jaccard index of dissimilarity between all parks (and contributions of species nestedness versus turnover to this metric); and (4) total number of species occurring at least once in any park in the city. We classified species as disturbance- or edge-avoidant if LepTraits v1.0 listed them with a “strong” or “weak” avoidance affinity [60]. We calculated the Jaccard index and its decomposition into nestedness and turnover using the *adespatial* package in R [65]. Repeating this simulation procedure across the 200 randomly sampled sets of parameters yielded a distribution of 200 plausible sets of city-wide biodiversity outcomes (Figure S15).

We applied generalized linear models (GLMs) to determine how variation in city-wide management impact these city-wide biodiversity outcomes (Figure S16). We hypothesized that these city-wide biodiversity responses would be affected by city-wide median *log* (park size), percent tree cover, percent herbaceous vegetation cover, and overall park connectivity. We quantified city-wide park connectivity as the (*log* (Integral Index of Connectivity (IIC))), implemented with the *lconnect* package in R [66], using a maximum interpark connectivity threshold of 2 km [57]. We calculated city-wide tree cover and herbaceous vegetation cover using the 1 m resolution UrbanWatch data [33]. We also included latitude as a predictor of both median local species richness and total species richness, because we expected fewer species at higher latitudes [67]. Finally, we also included *log* (total city area) as a predictor of total species richness, because we expected that measuring species richness over a larger spatial area would increase the number of species that occur [50]. Given, the limited sample size of 22 independent cities, we conducted a two step analysis. First, we fit an initial model that did not propagate uncertainty, treating the mean value of each biodiversity response as a function of all predictors of interest. With the initial model, we identified predictors that showed at least marginal evidence for an association with the responses (i.e., at least the 50% BCI departing from 0). Then, using the subset of predictors identified by the initial model, we fit GLMs that fully propagated uncertainty in the biodiversity responses.

We fit the posthoc GLMs using the a Bayesian approach with the “stan glm()” function from the *rstanarm* package in R [63]. We used default prior settings, which placed weakly-informative constraints on potential parameter values [63]. To propagate uncertainty, we looped the GLM fitting procedure across the distributions of plausible biodiversity responses. For each GLM fit, we saved 40 random samples of sets of parameter values from the posterior distribution. This yielded an assembled posterior distribution of 8,000 sets of posterior parameter values that propagated the uncertainty from the outputs of our original occupancy model (40 samples per each of the 200 plausabile diversity responses) [64]. Because the number of disturbance- or edge-avoidant species was a continuous positive value that was close to zero, we used GLMs with a gamma probability distribution and a log-link function. All other biodiversity metrics were modeled as Gaussian responses.

## Supporting information

Supplemental Materials: Species Detections Table

## Acknowledgments

This research was enabled in part by support provided by the Digital Research Alliance of Canada. We thank all iNaturalist users for collecting and sharing the butterfly detection data used in this study.

## Data Availability Statement

The community science data used in the analyses are available directly from GBIF [55, 58]. Land cover data were obtained from previously published sources [33, 34, 54]. All code used to process the data and conduct the analyses are currently available through github: https://github.com/jensculrich/urban_park_community_science_project. Once the manuscript is accepted for publication, we will archive the code and data on a permanent Zenodo repository.

## Supplemental Material

**Supplemental Figure S1:**
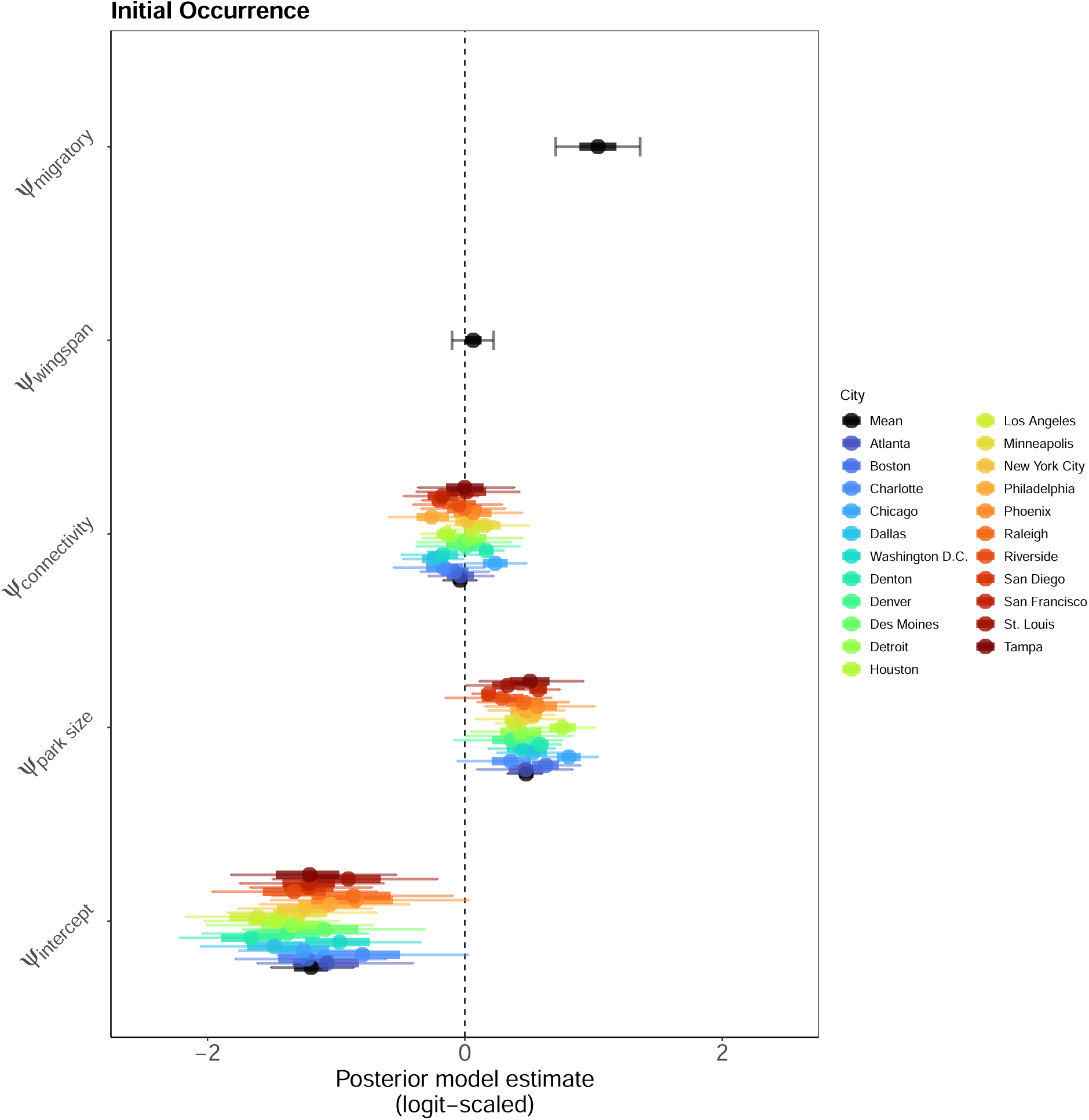
Butterfly Species Initial Occurrence - Probability of species occupancy at a site in the first year that the butterfly family was surveyed at the site. Solid circles indicate posterior estimate means. Bayesian credible intervals (50% (heavy band) and 90% (narrow band)) convey uncertainty in the magnitude of the estimates. Posterior estimates for the mean city are shown in black, while colors communicate estimates for city-specific random effects. The intercept term indicates the probability of initial occurrence for the average butterfly species at the average urban park site. For the hypothesized effect parameters (park size, connectivity, wingspan, and migratory status), posterior parameter distributions with a high density above zero suggests a positive association while a high density below zero suggests a negative association.

**Supplemental Figure S2:**
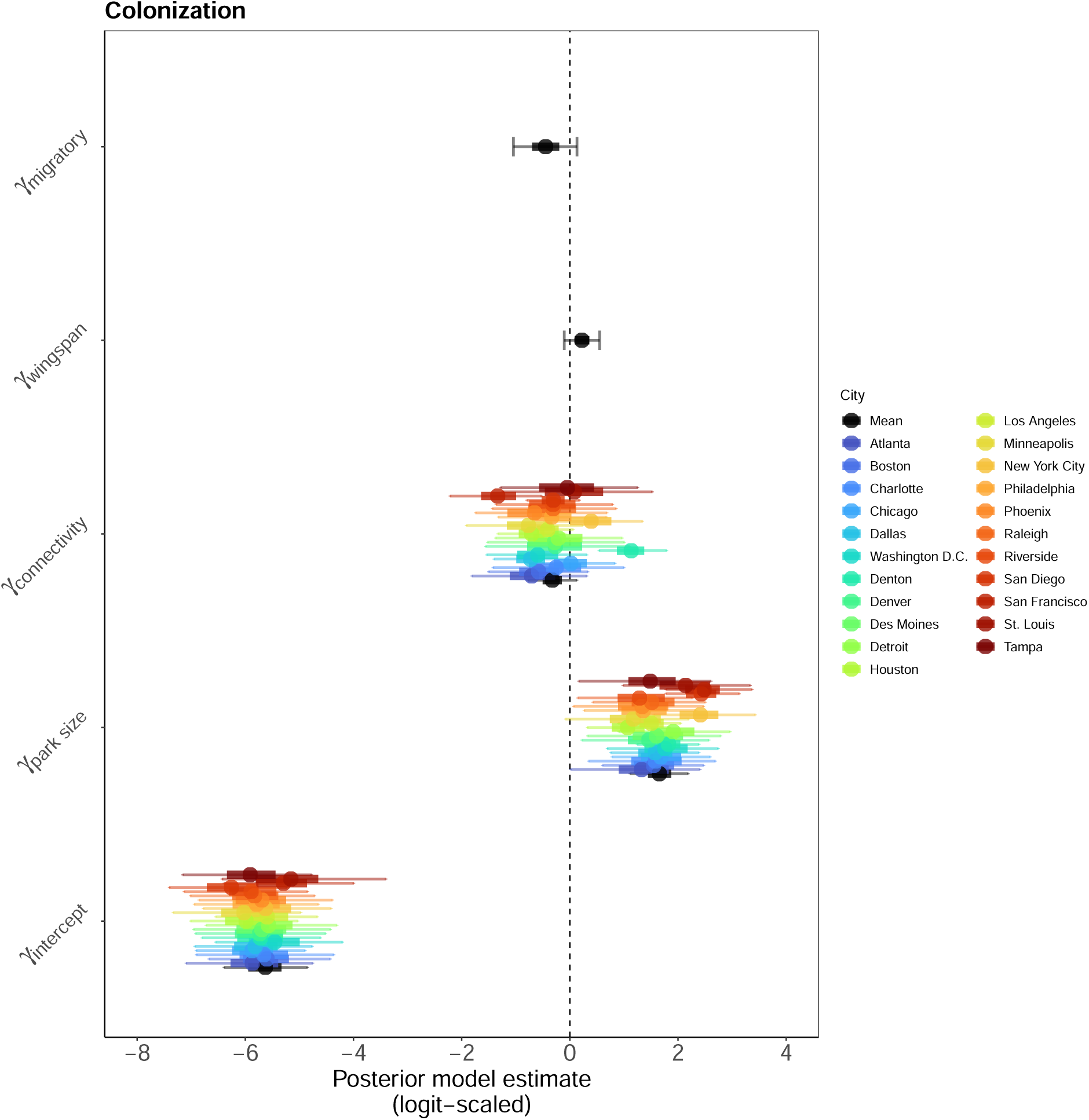
Butterfly Species Colonization - Probability of a species occurring at a site where it did not occur in a previous year. Solid circles indicate posterior estimate means. Bayesian credible intervals (50% (heavy band) and 90% (narrow band)) convey uncertainty in the magnitude of the estimates. Posterior estimates for the mean city are shown in black, while colors communicate estimates for city-specific random effects. The intercept term indicates the probability of colonization for the average butterfly species at the average urban park site. For the hypothesized effect parameters (park size, connectivity, wingspan, and migratory status), posterior parameter distributions with a high density above zero suggests a positive association while a high density below zero suggests a negative association.

**Supplemental Figure S3:**
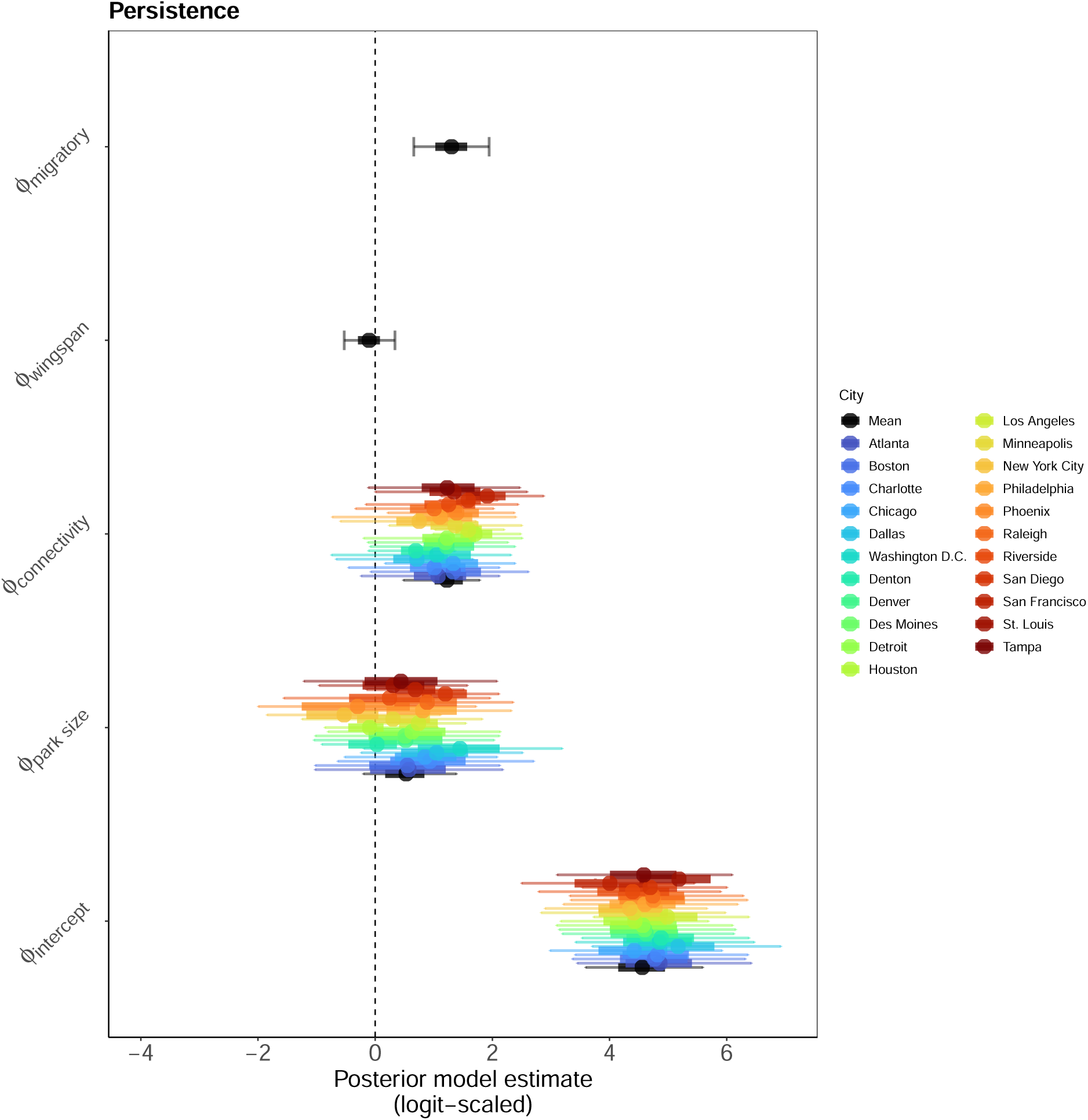
Butterfly Species Persistence - Probability of a species occurring at a site where it occurred in a previous year. Solid circles indicate posterior estimate means. Bayesian credible intervals (50% (heavy band) and 90% (narrow band)) convey uncertainty in the magnitude of the estimates. Posterior estimates for the mean city are shown in black, while colors communicate estimates for city-specific random effects. The intercept term indicates the probability of initial occurrence for the average butterfly species at the average urban park site. For the hypothesized effect parameters (park size, connectivity, wingspan, and migratory status), posterior parameter distributions with a high density above zero suggests a positive association while a high density below zero suggests a negative association.

**Supplemental Figure S4:**
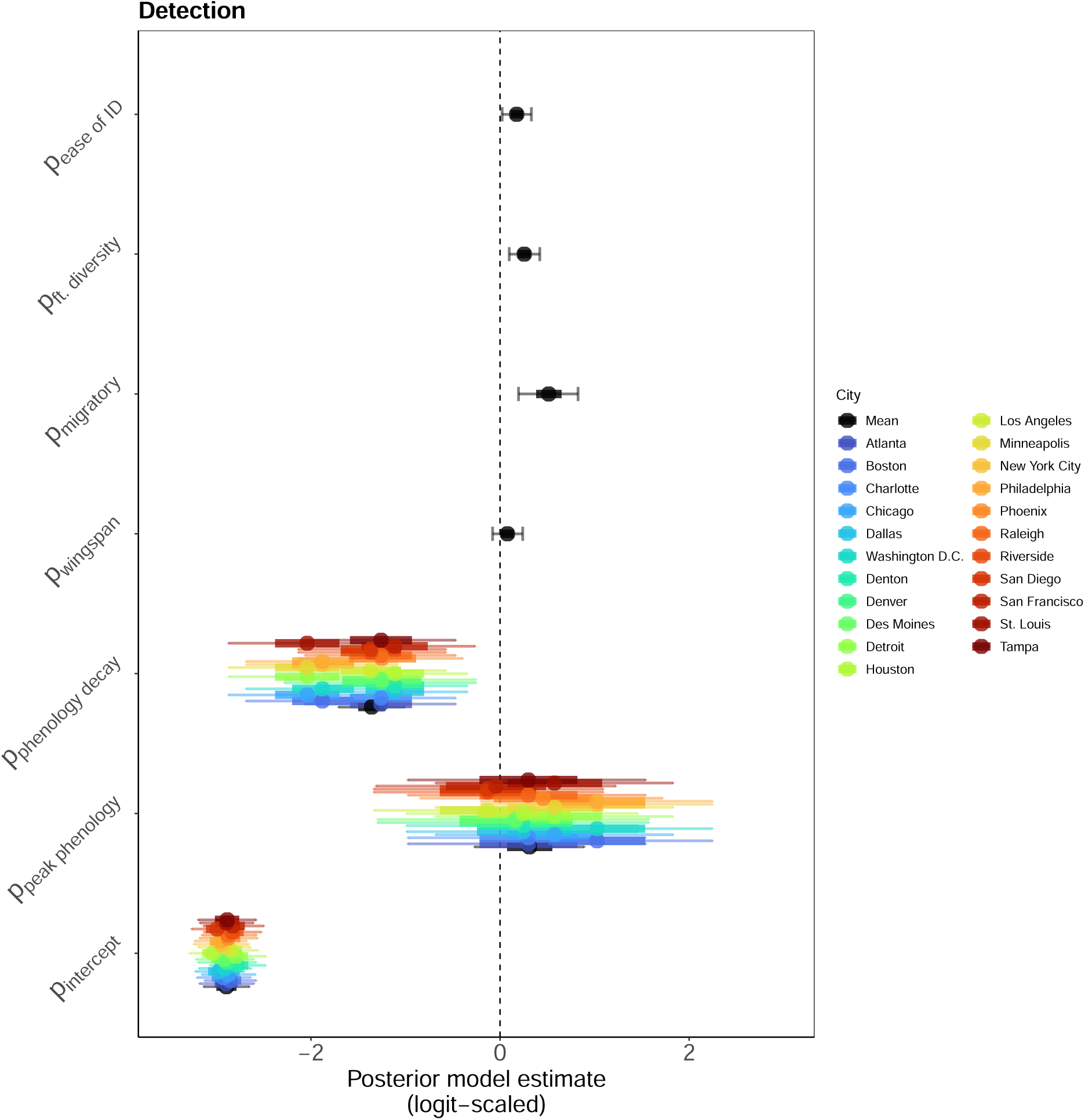
Butterfly Species Detection - Probability of detecting a species given that it occurs and that a survey effort was inferred. Solid circles indicate posterior estimate means. Bayesian credible intervals (50% (heavy band) and 90% (narrow band)) convey uncertainty in the magnitude of the estimates. Posterior estimates for the mean city are shown in black, while colors communicate estimates for city-specific random effects. The intercept term indicates the probability of detection for the average butterfly species at the average urban park site. For the hypothesized effect parameters (park size, connectivity, wingspan, and migratory status), posterior parameter distributions with a high density above zero suggests a positive association while a high density below zero suggests a negative association. Baseline detection rate (the probability of detecting the average butterfly species given that at least one other species from the same family was detected in the same park during the same time interval) was *∼* 5% (*p*[intercept] 90% BCI = [4.0%, 6.7%]) (Figure S4). Migratory species were more likely to be detected relative to non-migratory species (mean = 0.52, 90% BCI = [0.19, 0.86]) (Figure S4). Feature diversity (the number of features on the wings such as checkers, spots, tails, etc.) and ease of identification (measured as the proportion of species in a genus that are research grade [59]) also increased detection rate (mean = 0.25, 90% BCI = [0.10, 0.41]; mean = 0.17, 90% BCI = [0.03, 0.33]) (Figure S4).

**Supplemental Figure S5:**
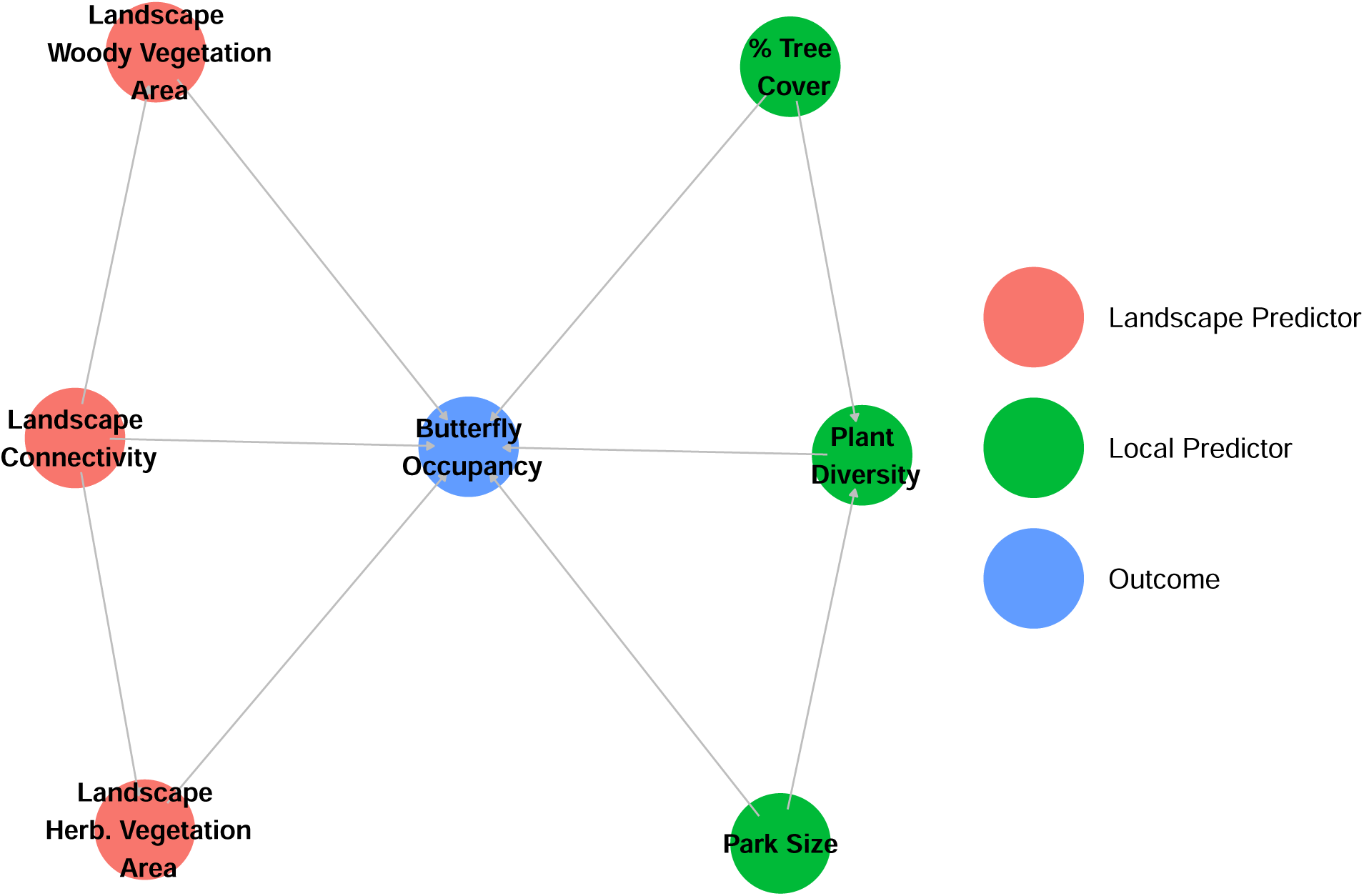
Directed Acyclic Graph (DAG) showing hypothesized causal relationships between local (green) and landscape (red) park characteristics of butterfly occupancy in urban parks. We hypothesized that park size would indirectly impact butterfly occupancy plant diversity. We also hypothesized that increased tree cover within a park would also indirectly impact butterfly occupancy by increasing the diversity of flowering plant species (i.e., more tree cover would equate to more diversity of trees that provide nectar or larval host resources). We hypothesized that spatial connectivity to nearby parks would also indirectly impact butterflies by increasing the area of woody and herbaceous vegetation surrounding the park sites. In analysis part 2, we quantified the direct effects of park size and local tree cover by stratifying by plant diversity, and the direct effect of park connectivity by stratifying by the amount of woody and herbaceous vegetation cover in the surrounding landscape. We present the results from this model, model m2.1, in the main text and figures. We then quantified the total effects of mediated occupancy predictors (park size, local tree cover, and park connectivity) by dropping mediator variables (plant diversity, landscape woody vegetation area, and landscape herbaceous vegetation area) from the model. We present the results of this second, model m2.2, in the supplemental figures.

**Supplemental Figure S6:**
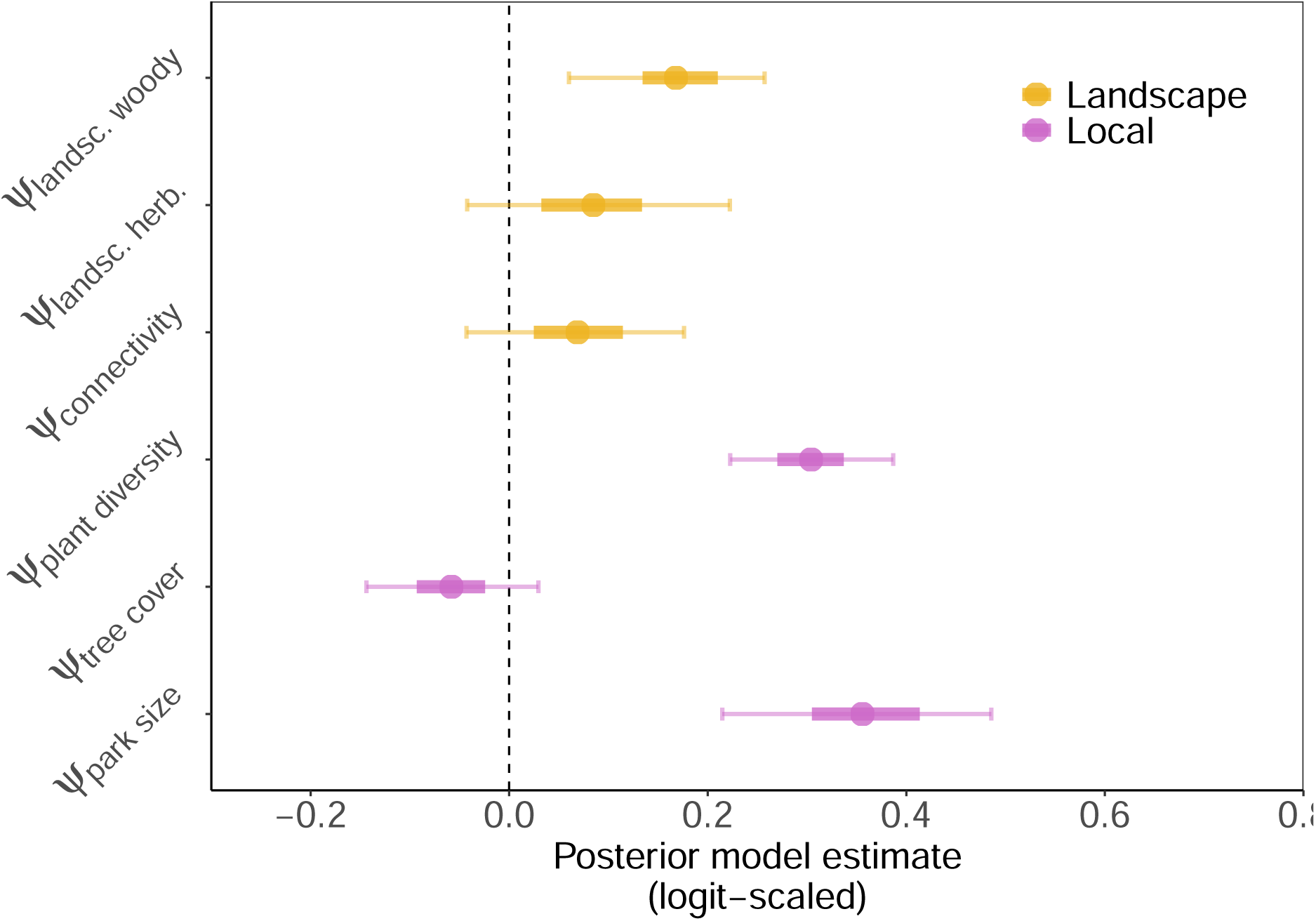
Mean-city effects of local and landscape park characteristics on static patterns of butterfly species occupancy. All park parameters were treated as city-specific random effects; posterior estimates for the mean city are displayed. Solid circles indicate posterior means while Bayesian credible intervals (50% (heavy band) and 90% (narrow band)) convey uncertainty in the estimates. Posterior estimates are colored by predictor category (local or landscape). Posterior distributions with a high density above zero suggest a positive effect while a high density below zero suggests a negative association.

**Supplemental Figure S7:**
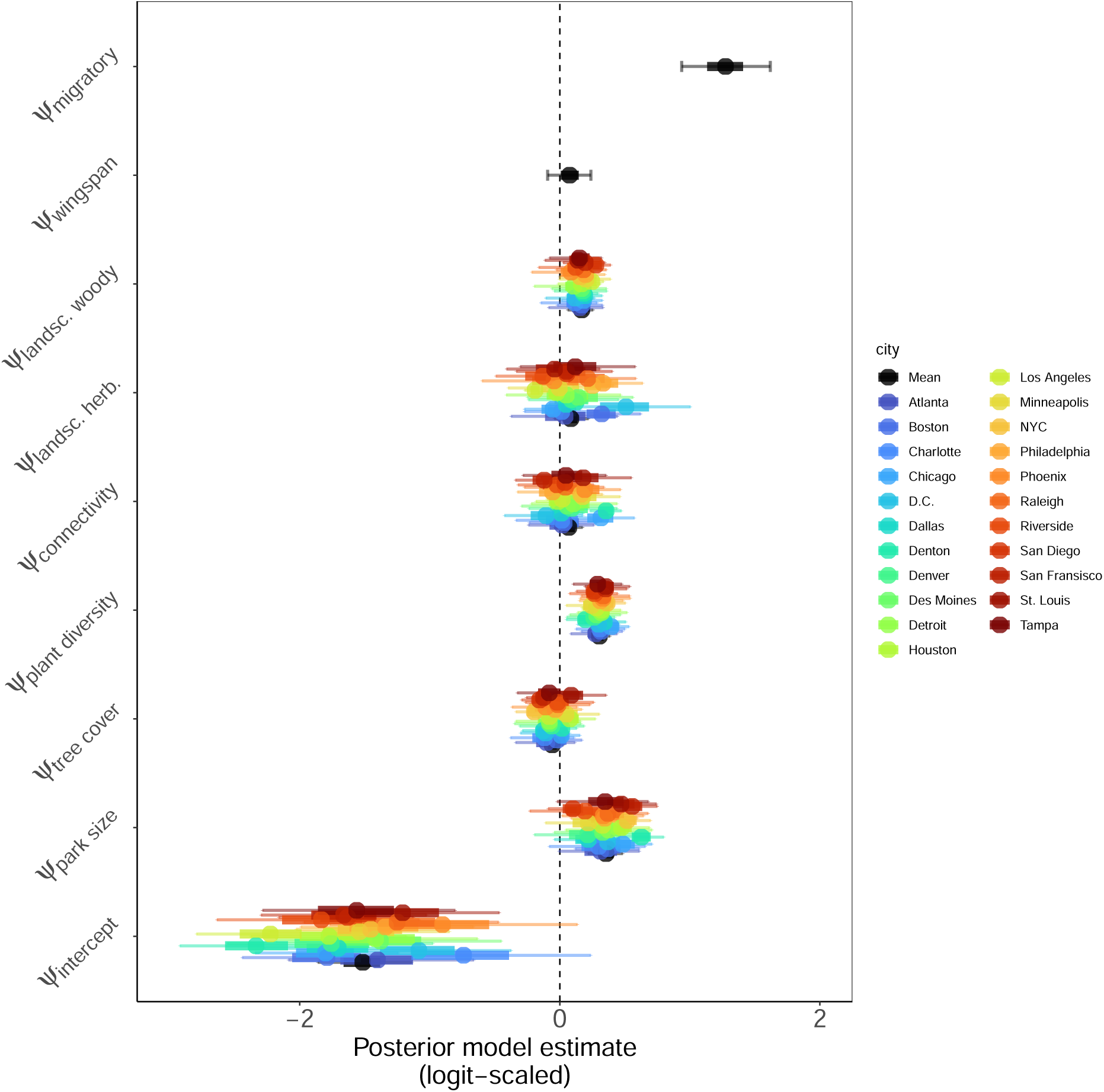
Butterfly Species Occupancy (Static Occupancy Model) - Probability of a species occurring at a site in any given year. Solid circles indicate posterior estimate means. Bayesian credible intervals (50% (heavy band) and 90% (narrow band)) convey uncertainty in the magnitude of the estimates. Posterior estimates for the mean city are shown in black, while colors communicate estimates for city-specific random effects. The intercept term indicates the probability of initial occurrence for the average butterfly species at the average urban park site. For the hypothesized effect parameters (park size, connectivity, wingspan, and migratory status), posterior parameter distributions with a high density above zero suggests a positive association while a high density below zero suggests a negative association.

**Supplemental Figure S8:**
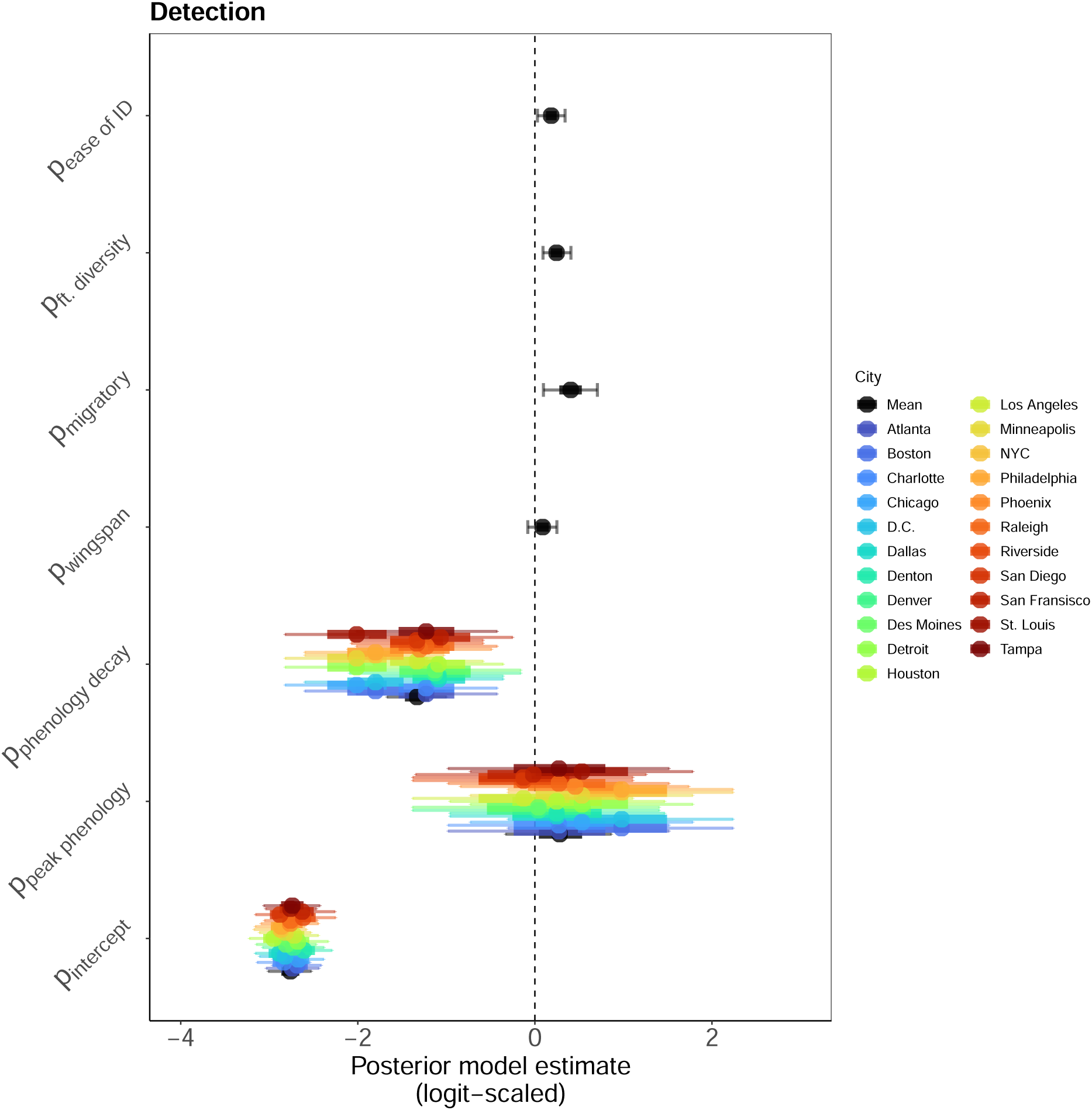
Butterfly Species Detection (Static Occupancy Model) - Probability of detecting a species given that it occurs and that a survey effort was inferred. Solid circles indicate posterior estimate means. Bayesian credible intervals (50% (heavy band) and 90% (narrow band)) convey uncertainty in the magnitude of the estimates. Posterior estimates for the mean city are shown in black, while colors communicate estimates for city-specific random effects. The intercept term indicates the probability of detection for the average butterfly species at the average urban park site. For the hypothesized effect parameters (park size, connectivity, wingspan, and migratory status), posterior parameter distributions with a high density above zero suggests a positive association while a high density below zero suggests a negative association.

**Supplemental Figure S9:**
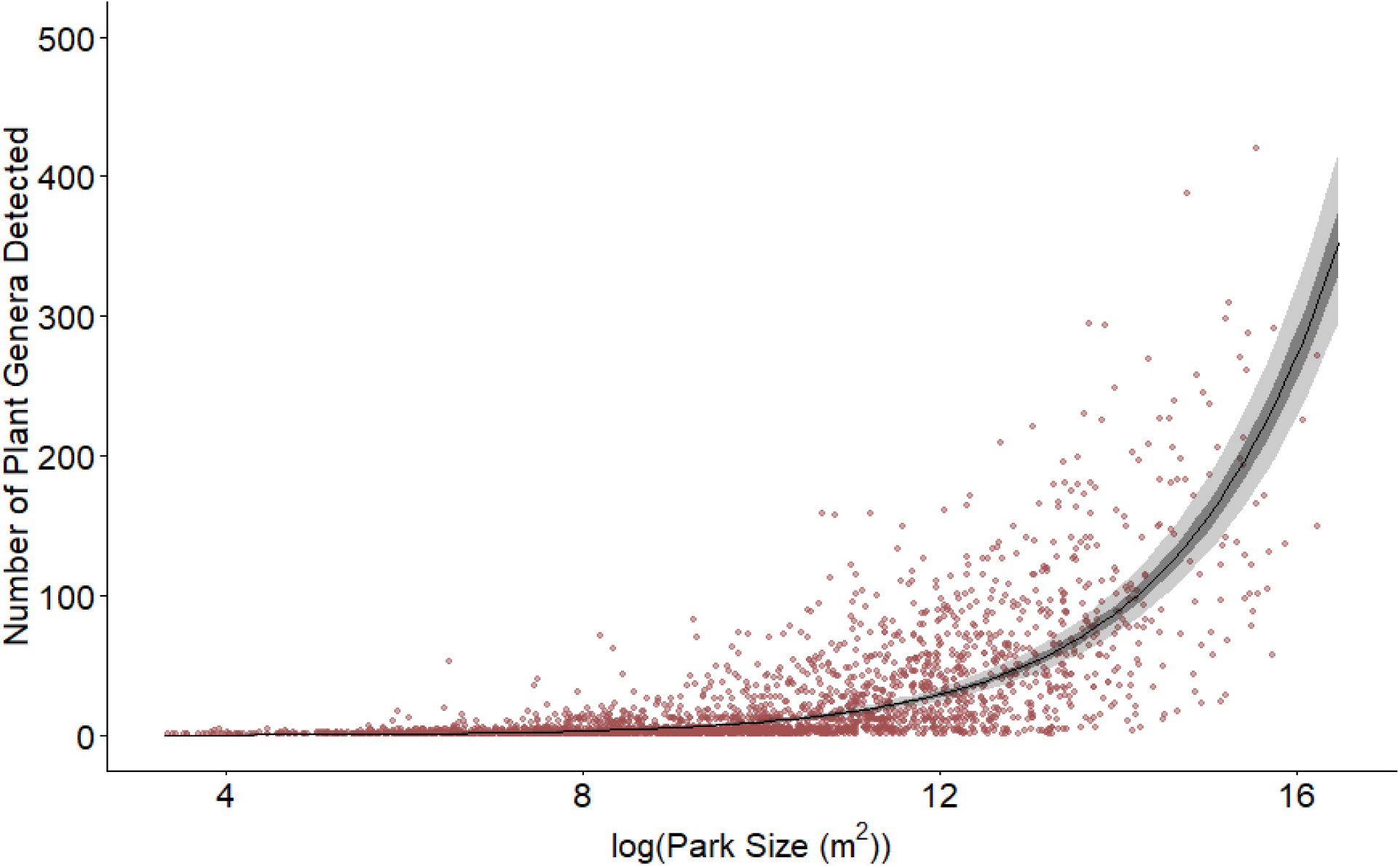
We used a Poisson GLM to describe the relationship between park size and plant diversity. Park size was positively associated with the number of flowering plant genera detected within a park (Mean estimate (log-scaled) = 1.465, 90%BCI = [1.457, 1.47]. Red points show the number of flowering plants detected in each of the 2,550 parks included in the analysis in part 2. The black line is the estimated mean relationship between park size and plant diversity. Dark and light grey bands display the 50 and 90% BCIs for the mean relationship between park size and plant diversity. The model included a city-specific random intercept effect and was fit with the ”stan glmer()” function from the R package *rstanarm* using default settings.

**Supplemental Figure S10:**
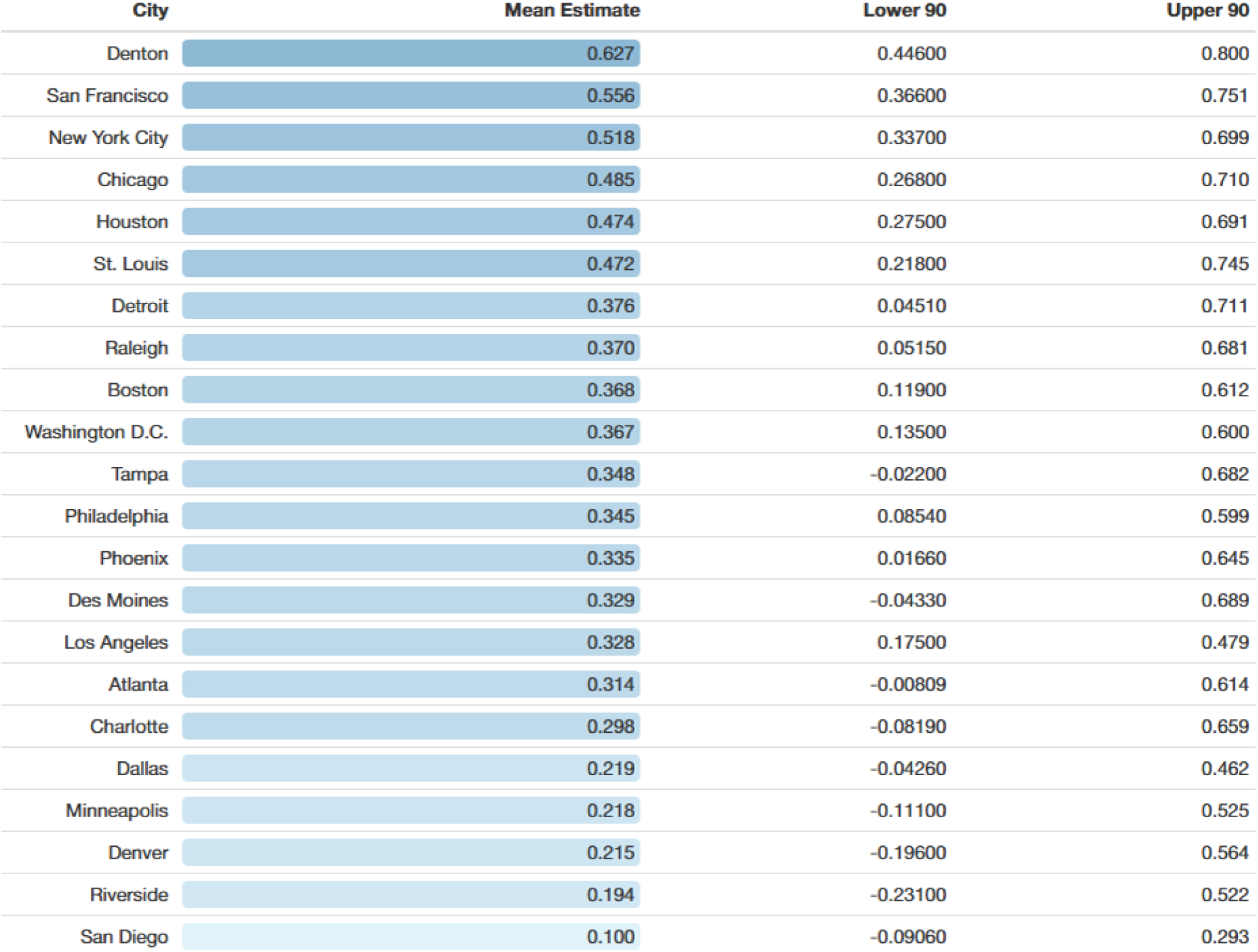
City-specific effects of park size on occupancy rate, arranged from largest (dark blue) to smallest (light blue) effect size.

**Supplemental Figure S11:**
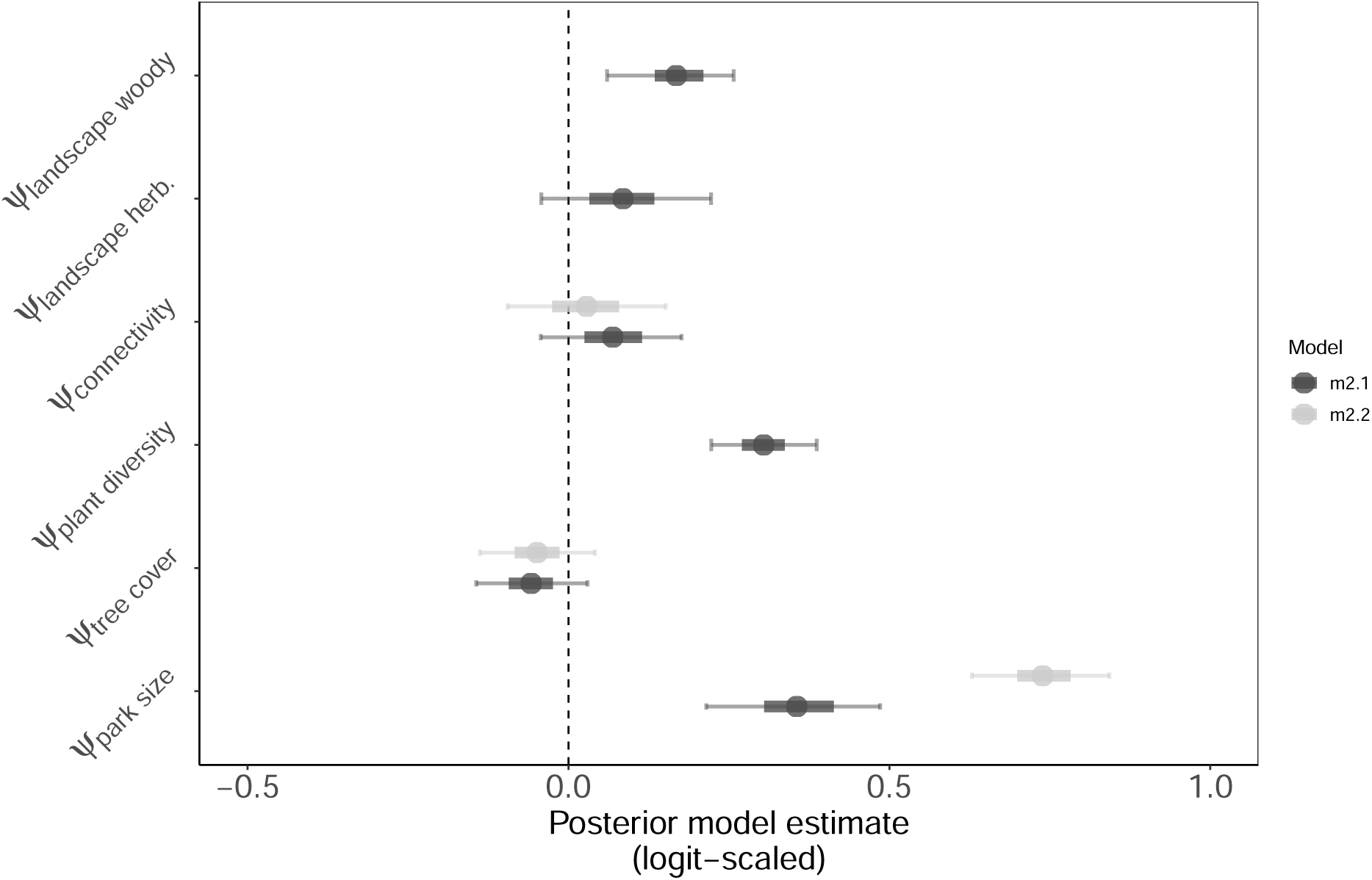
Posterior parameter estimates for static occupancy models with and without including plant diversity as a predictor of occupancy (m2.1 and m2.2, respectively). Following our DAG (supplemental figure Sx), for m2.1, the parameter estimate for park size represents the direct effect of park size on occupancy. For m2.2, the parameter estimate for park size represents the total effect of park size on occupancy, which included a potential mediating effect of plant diversity. Similarly, for m2.1, the parameter estimate for tree cover represents the direct effect of local tree cover on occupancy. For m2.2, the parameter estimate for tree cover represents the total effect of local tree cover on occupancy, which included a potential mediating effect of plant diversity. Solid circles indicate posterior estimate means. Bayesian credible intervals (50% (heavy band) and 90% (narrow band)) convey uncertainty in the magnitude of the estimates. Posterior estimates are shown for the mean city effect.

**Supplemental Figure S12:**
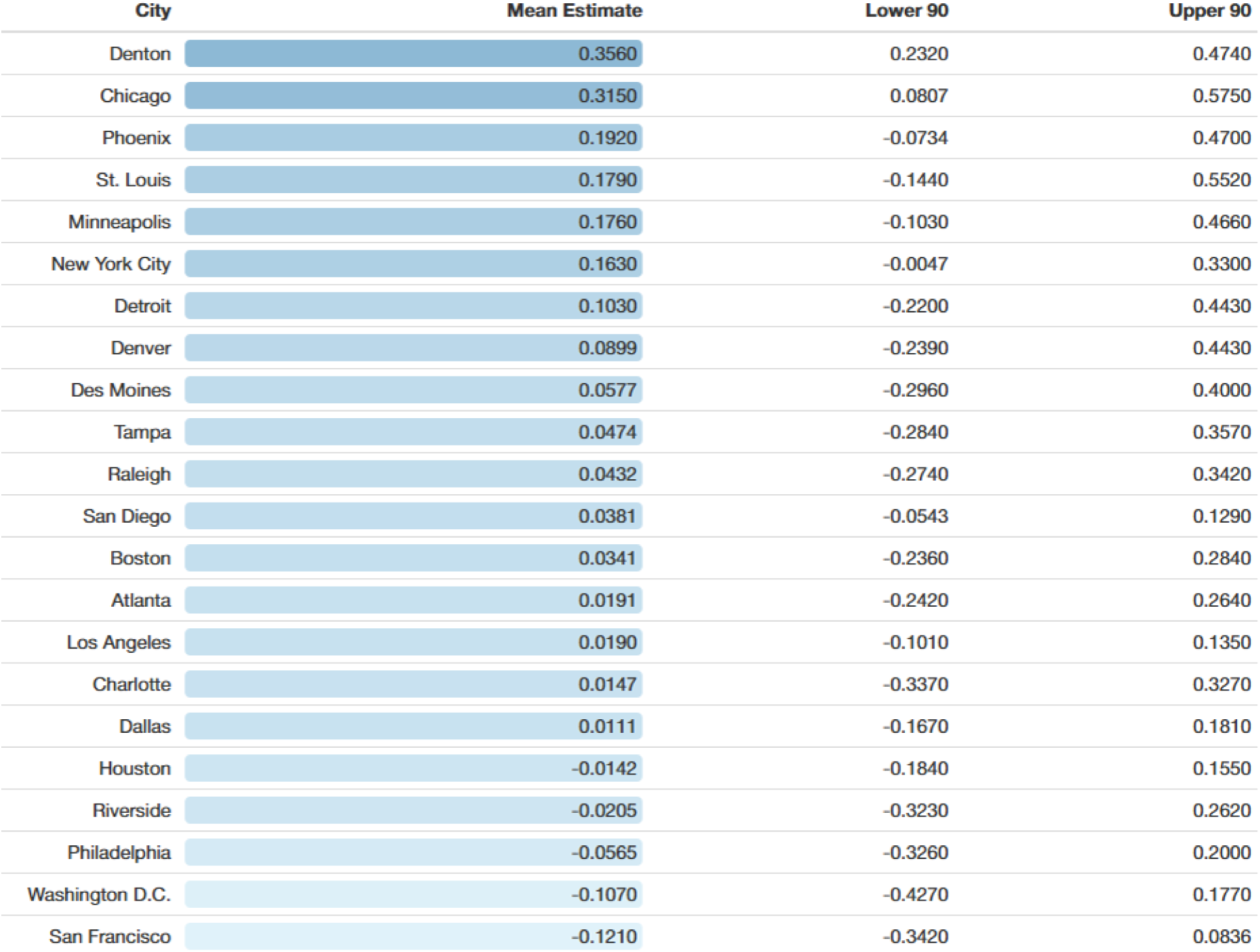
City-specific effects of park connectivity on occupancy rate, arranged from largest (dark blue) to smallest (light blue) effect size.

**Supplemental Figure S13:**
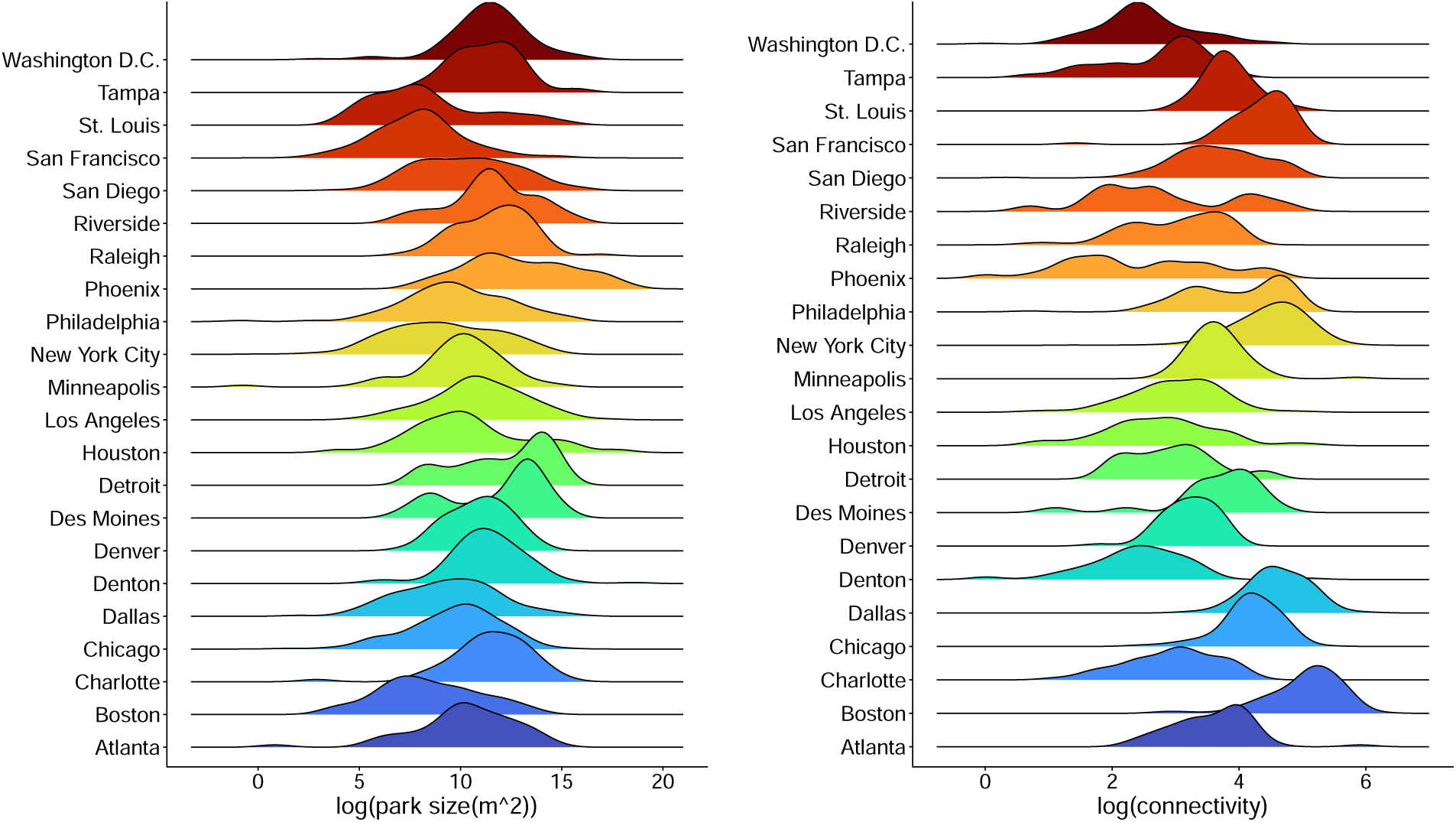
Distribution of park size and connectivity for modelled parks (parks with 1 or more survey events), coloured by city.

**Supplemental Figure S14:**
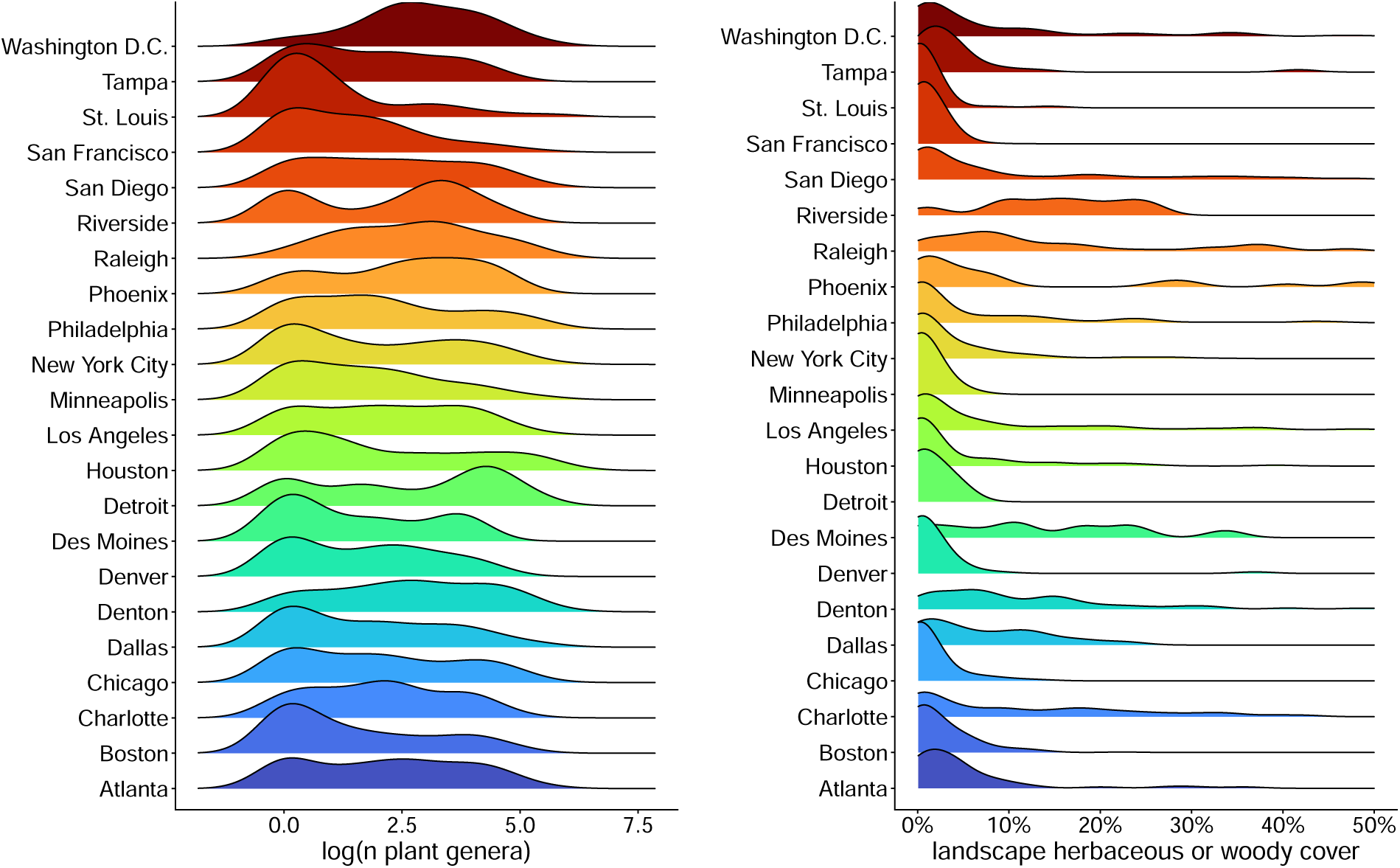
Distribution of plant diversity and landscape vegetation cover (summed herbaceous and woody cover within 2 km buffer) for modelled parks (parks with 1 or more survey events), coloured by city.

**Supplemental Figure S15:**
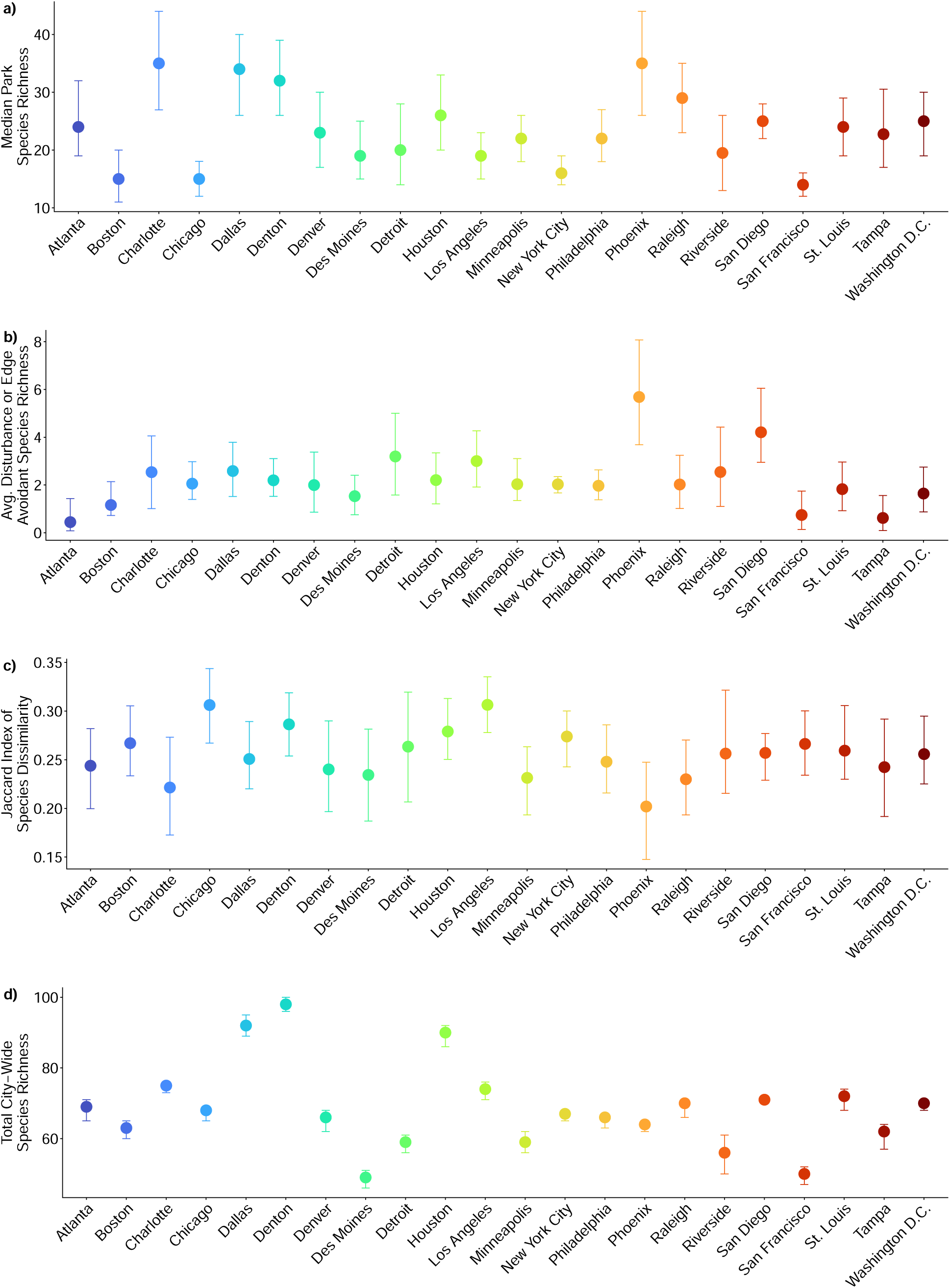
(Figure on previous page) City-wide butterfly diversity metrics including: median within-park species richness (a), average number of disturbance or edge-avoidant species in a community (b), average Jaccard index of dissimilarity in species composition (c), and total city-wide species richenss (d). Error bars display 90% Bayesian credible intervals.

**Supplemental Figure S16:**
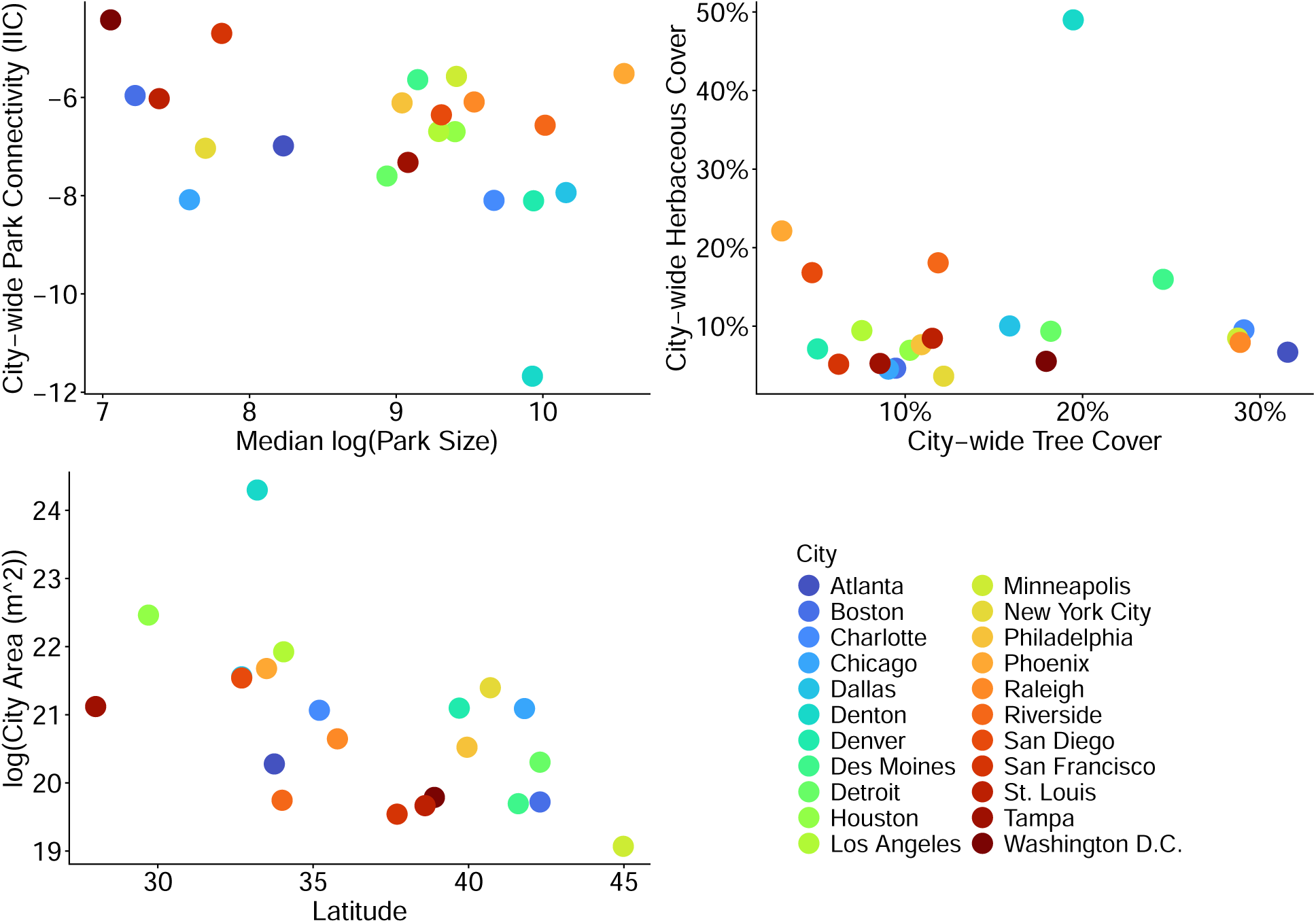
Distribution of city-wide covariate data used for Part 3 analyses, including city-wide median park size and connectivity (top-left), city-wide tree cover and herbaceous vegetation cover (top-right), and city latitude and spatial area (bottom-left). City identities are colour coded (bottom-right).

**Supplemental Figure S17:**
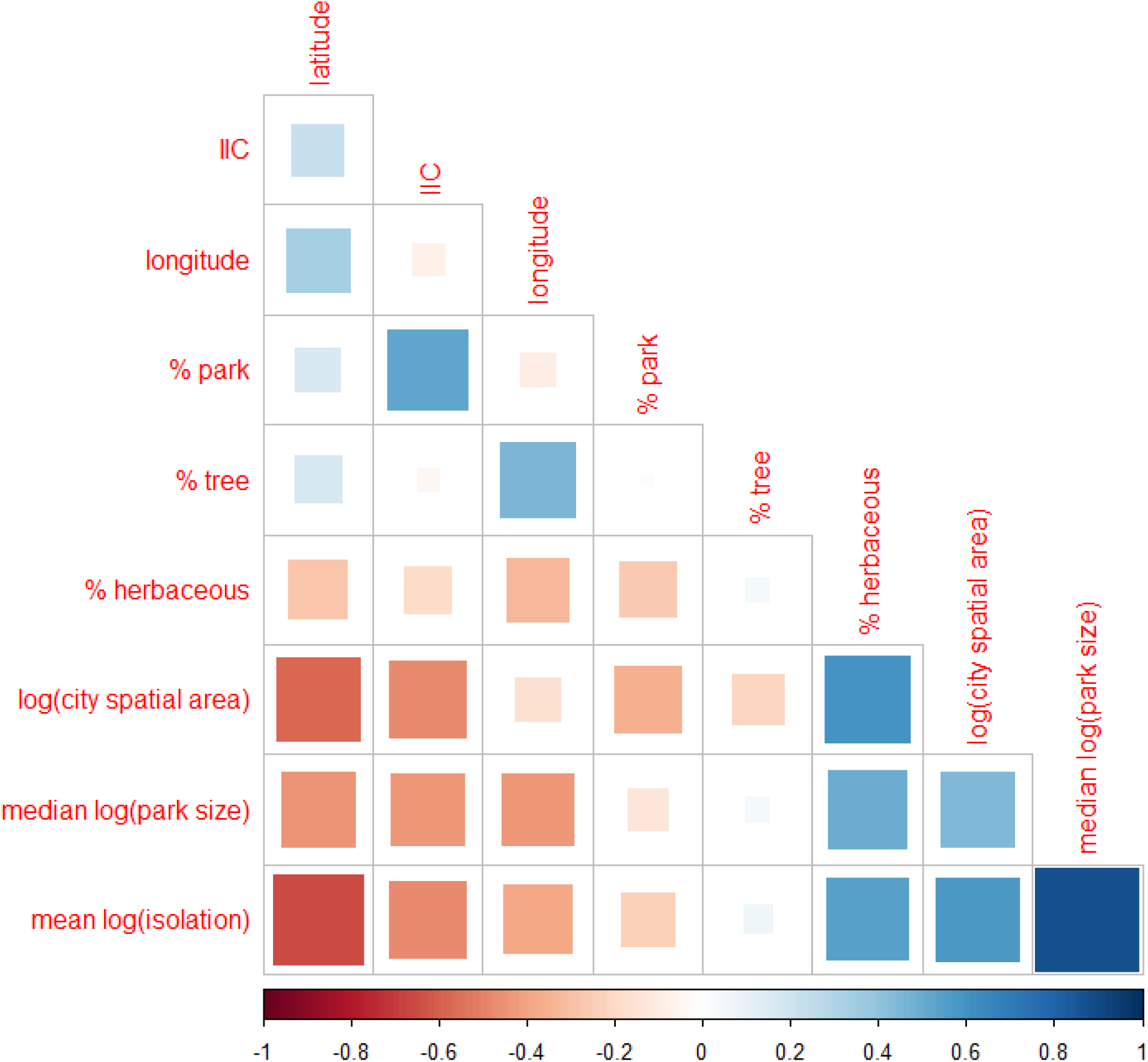
Correlations between city-wide features. In our models in part 3, we considered city spatial area, latitude, median log(park size), Integral Index of Connectivity (IIC), percentage of city area with herbaceous vegetation cover (% herbaceous), and percentage of city area with tree canopy cover (% tree). In this plot, we also show correlation values with city longitude, mean log(connectivity) (the average area- and distance-weighted connectivity metric that we computed for parts 1 and 2 and which should be inversely related to IIC connectivity), and log(total park area) (the total area of green space within parks summed across all parks within a city). Importantly, average park size is not related to % park area or % tree although to some extent positively correlated with % herbaceous. This means that the effects of average habitat patch size can be disentangled from the effects of total habitat area.

**Supplemental Figure S18:**
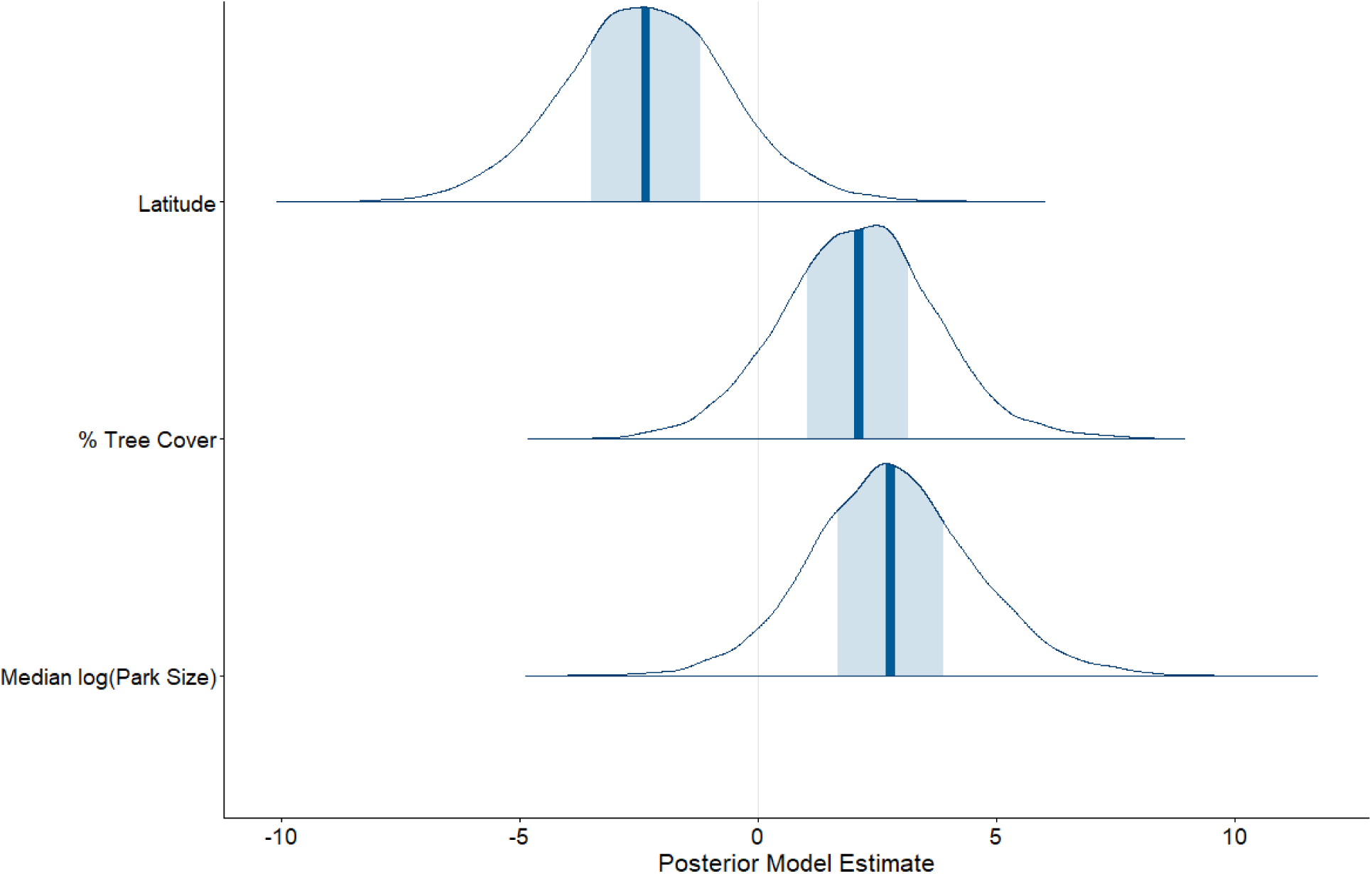
Posterior densities of parameter estimates for predictors of median park species richness. Dark blue bars indicate the median estimate, shaded blue areas indicate the 50% central quantiles of the posterior distributions and unshaded areas show the entire posterior distribution. In the final model, we only included parameters that had a clear directional effect on the mean value of the response (i.e., where at least the middle 50% BCI did not overlap with zero).

**Supplemental Figure S19:**
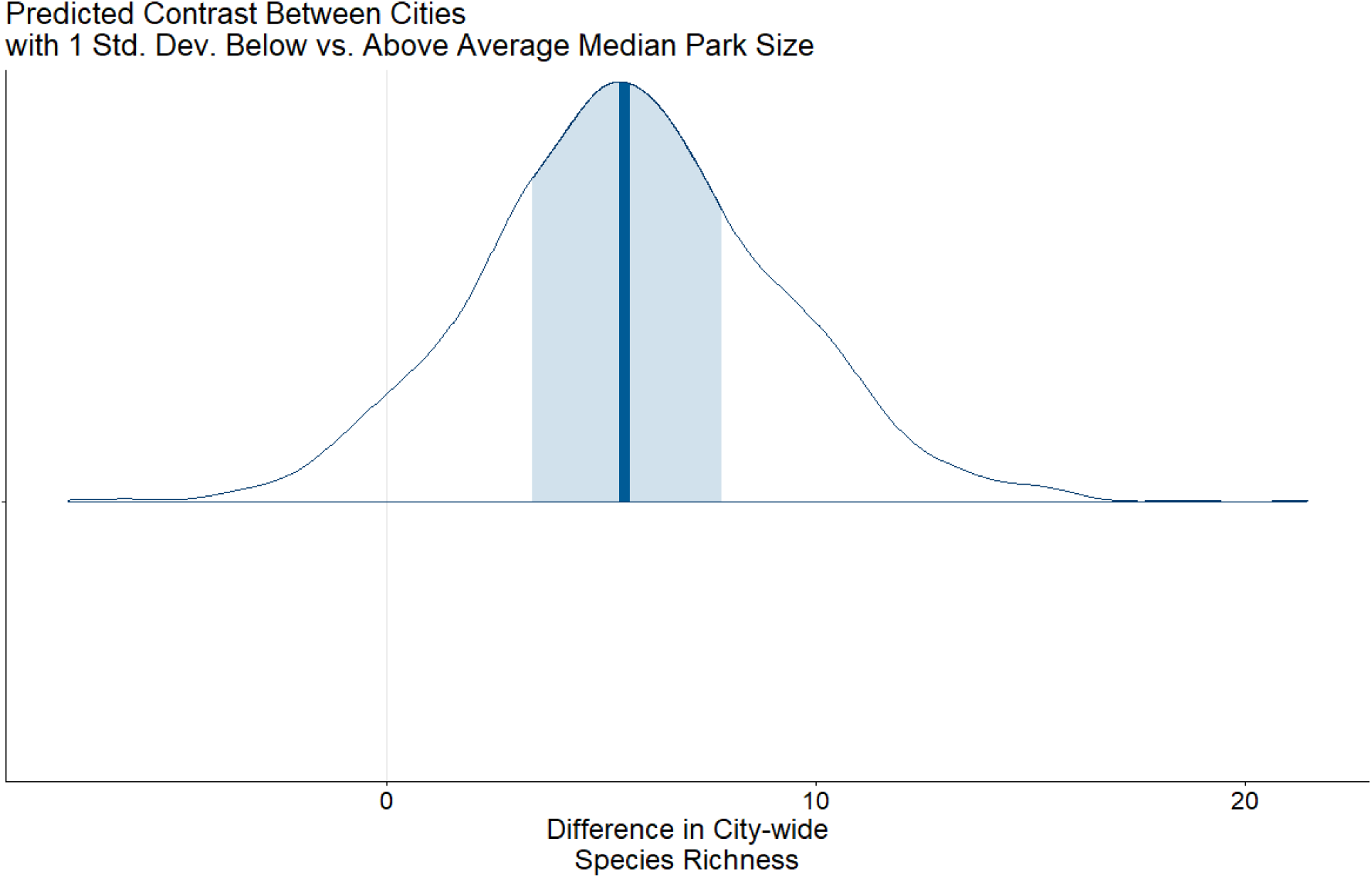
Posterior density of the predicted contrast between the median park species richness in a city with 1 standard deviation above average tree cover versus for a city with 1 standard deviation below average tree cover. Dark blue bars indicate the median estimated contrast, shaded blue areas indicate the 50% central quantiles of the posterior distribution of contrasts and unshaded areas show the entire posterior distribution of contrasts. Values greater than zero indicate a prediction of greater city-wide species richness in a city with higher than average tree cover.

**Supplemental Figure S20:**
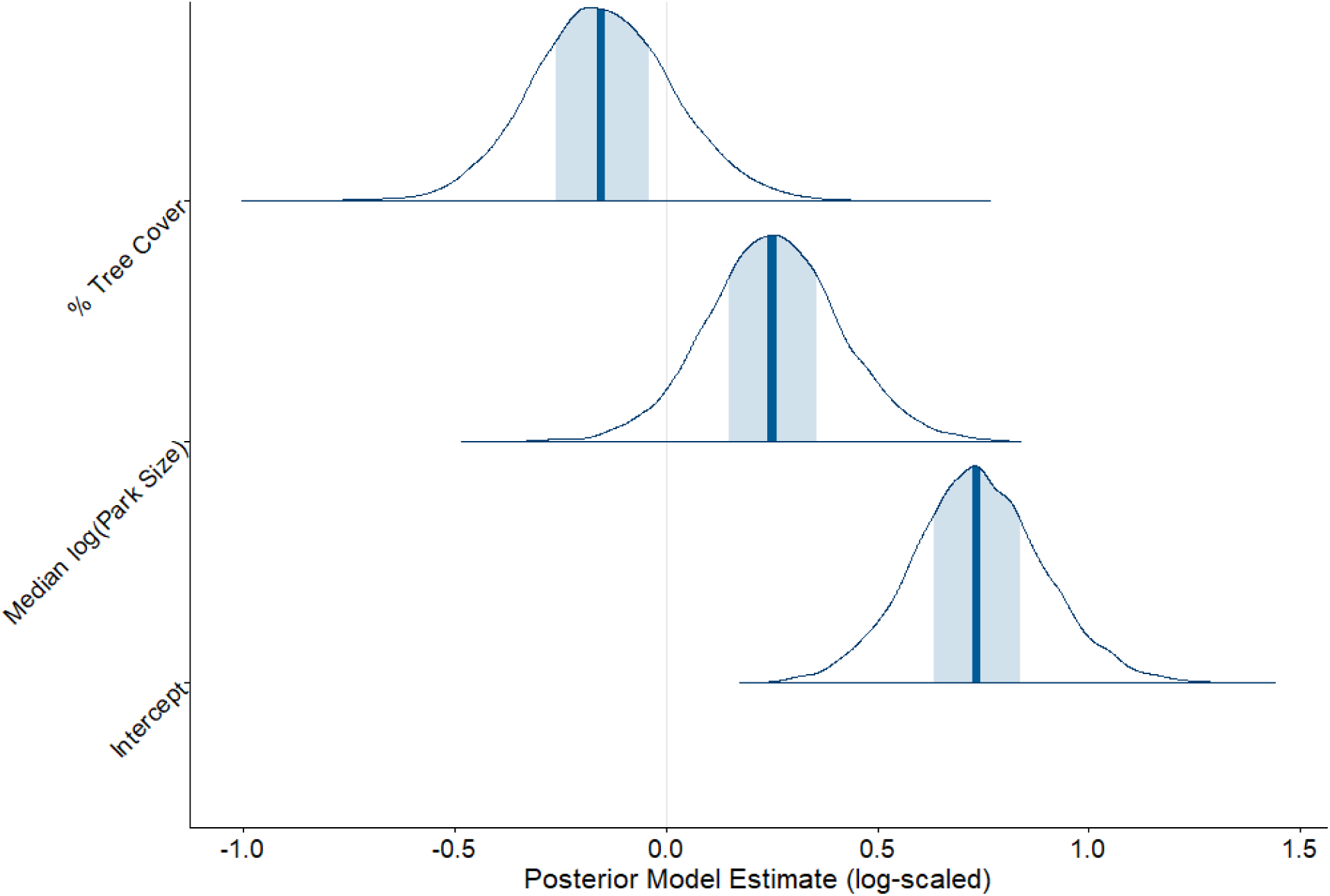
Posterior densities of parameter estimates for predictors of the number of disturbance- or edge-avoidant species in urban park communities. Dark blue bars indicate the median estimate, shaded blue areas indicate the 50% central quantiles of the posterior distributions and unshaded areas show the entire posterior distribution. In the final model, we only included parameters that had a clear directional effect on the mean value of the response (i.e., where at least the middle 50% BCI did not overlap with zero).

**Supplemental Figure S21:**
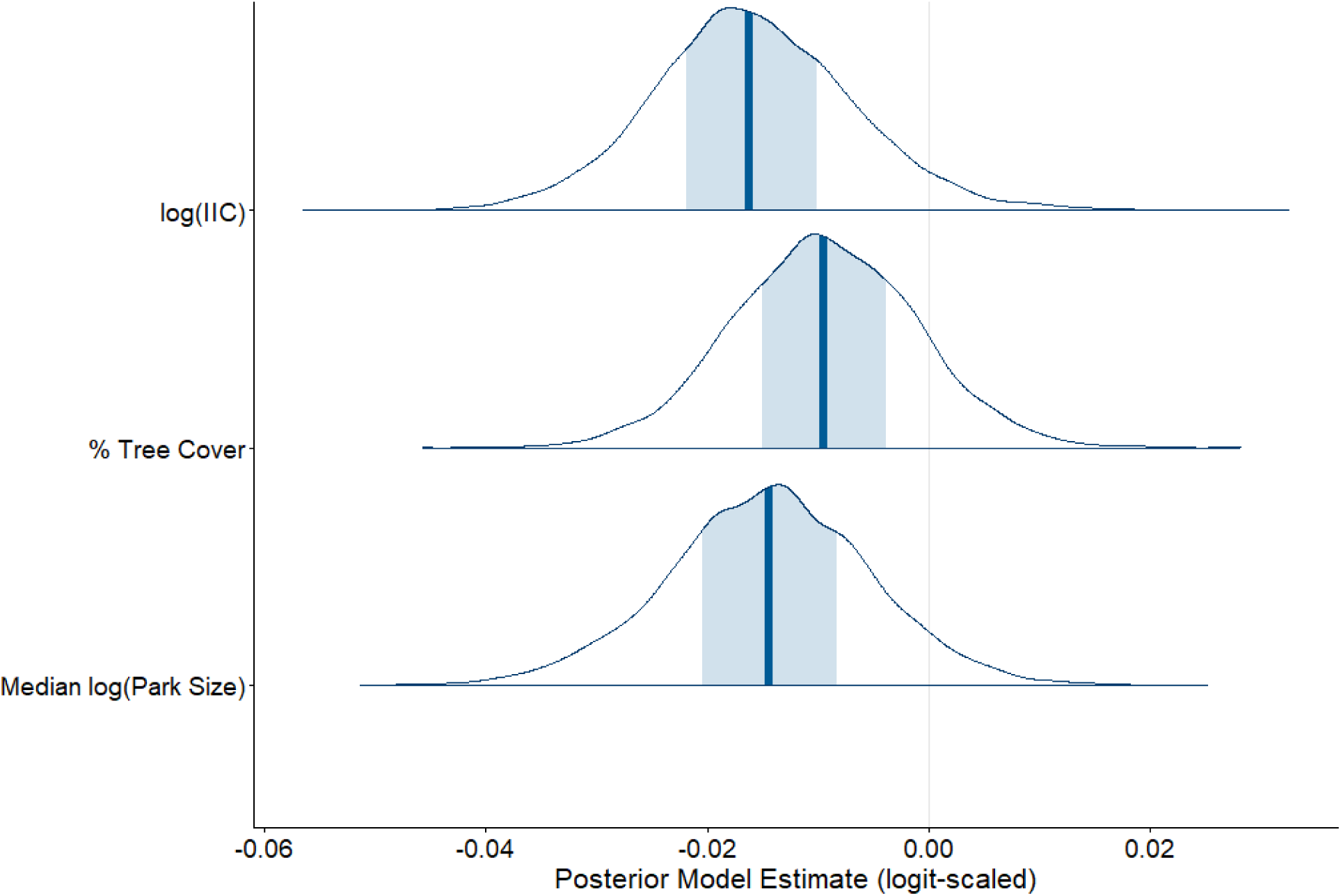
Posterior densities of parameter estimates for predictors of the Jaccard Index of Community Dissimilarity. Dark blue bars indicate the median estimate, shaded blue areas indicate the 50% central quantiles of the posterior distributions and un-shaded areas show the entire posterior distribution. In the final model, we only included parameters that had a clear directional effect on the mean value of the response (i.e., where at least the middle 50% BCI did not overlap with zero).

**Supplemental Figure S22:**
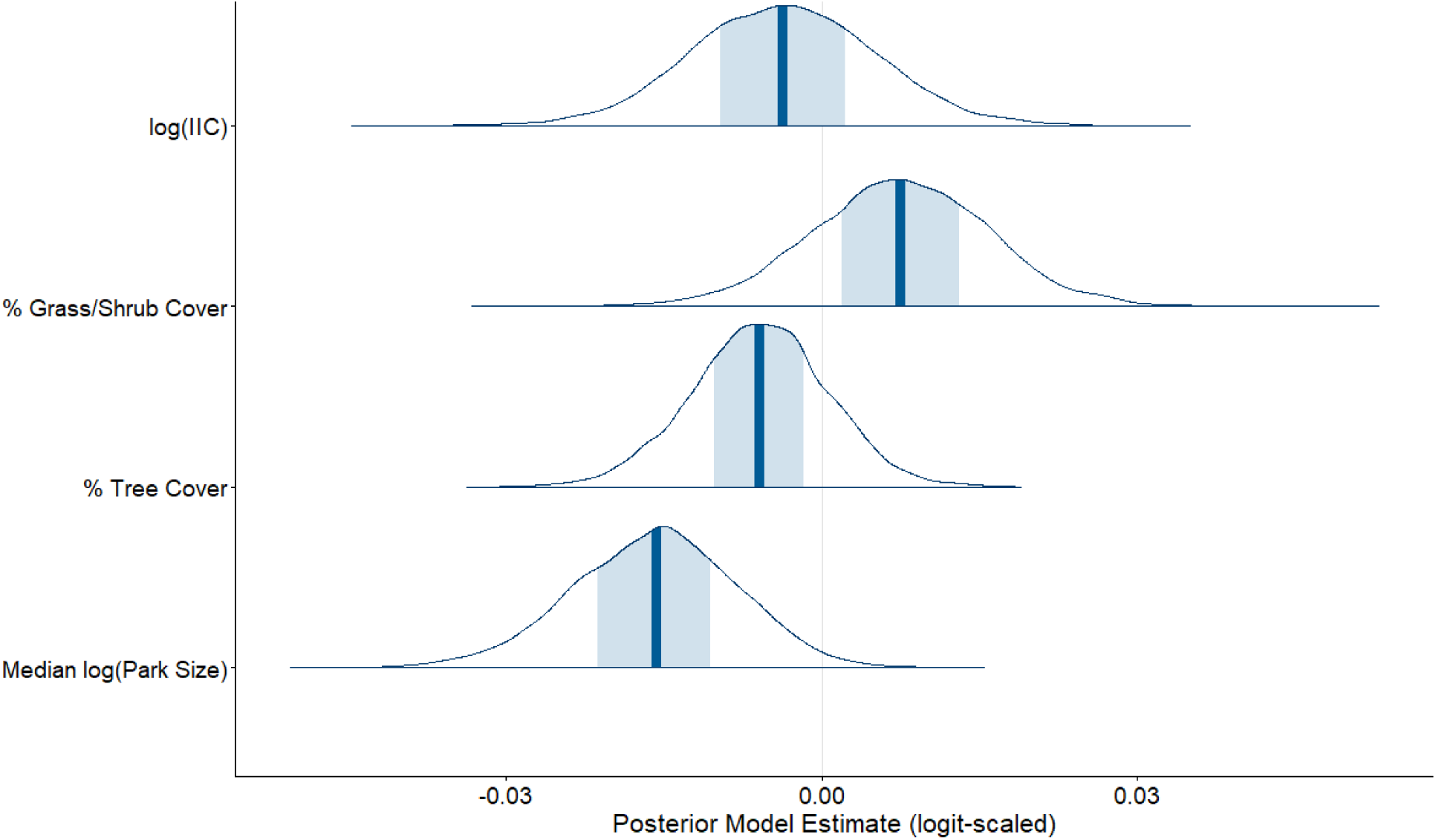
Posterior densities of parameter estimates for predictors of the contributions of species nestedness to the Jaccard Index of Community Dissimilarity. Dark blue bars indicate the median estimate, shaded blue areas indicate the 50% central quantiles of the posterior distributions and unshaded areas show the entire posterior distribution. In the final model, we only included parameters that had a clear directional effect on the mean value of the response (i.e., where at least the middle 50% BCI did not overlap with zero).

**Supplemental Figure S23:**
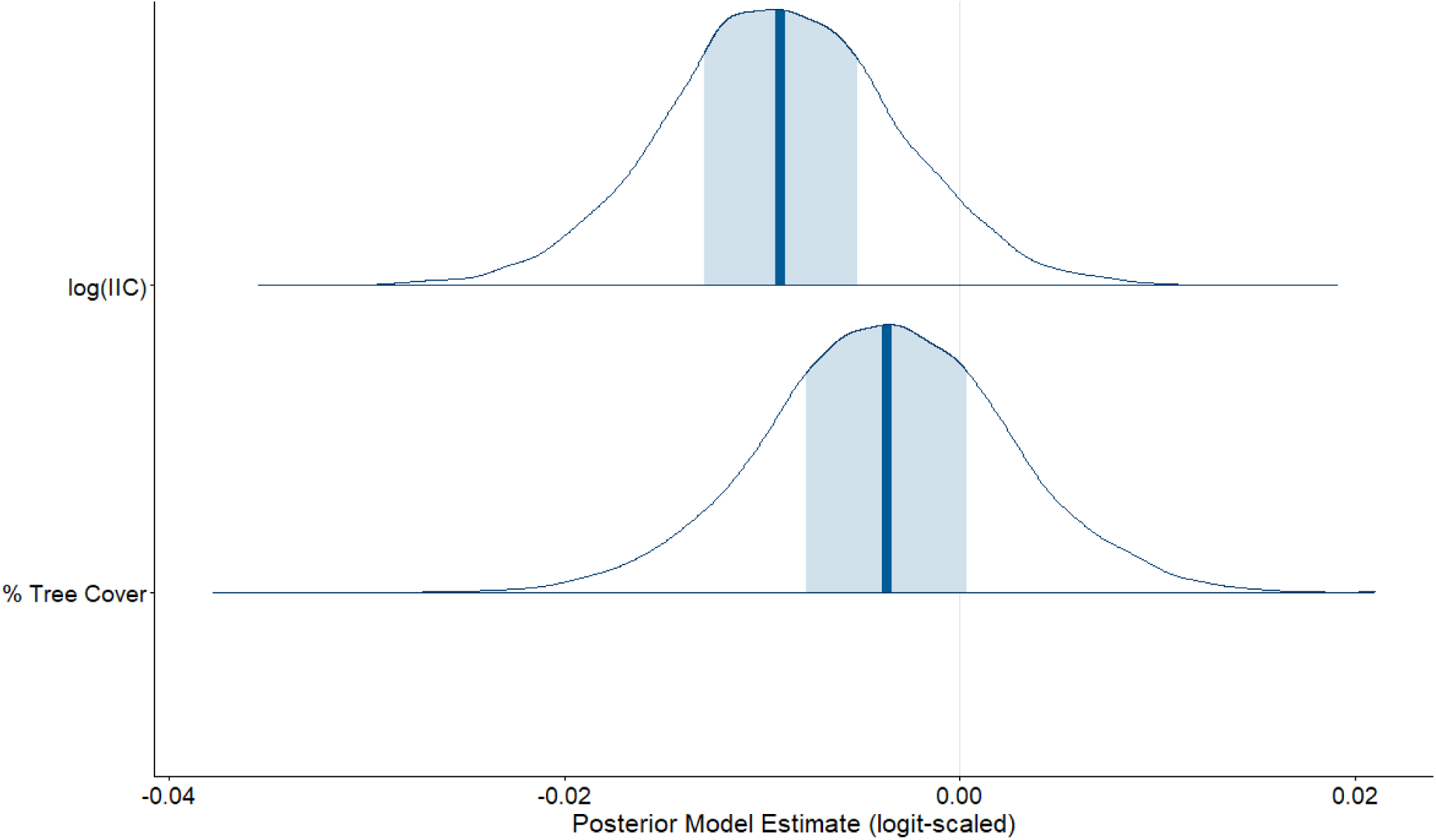
Posterior densities of parameter estimates for predictors of the contributions of species turnover to the Jaccard Index of Community Dissimilarity. Dark blue bars indicate the median estimate, shaded blue areas indicate the 50% central quantiles of the posterior distributions and unshaded areas show the entire posterior distribution. In the final model, we only included parameters that had a clear directional effect on the mean value of the response (i.e., where at least the middle 50% BCI did not overlap with zero).

**Supplemental Figure S24:**
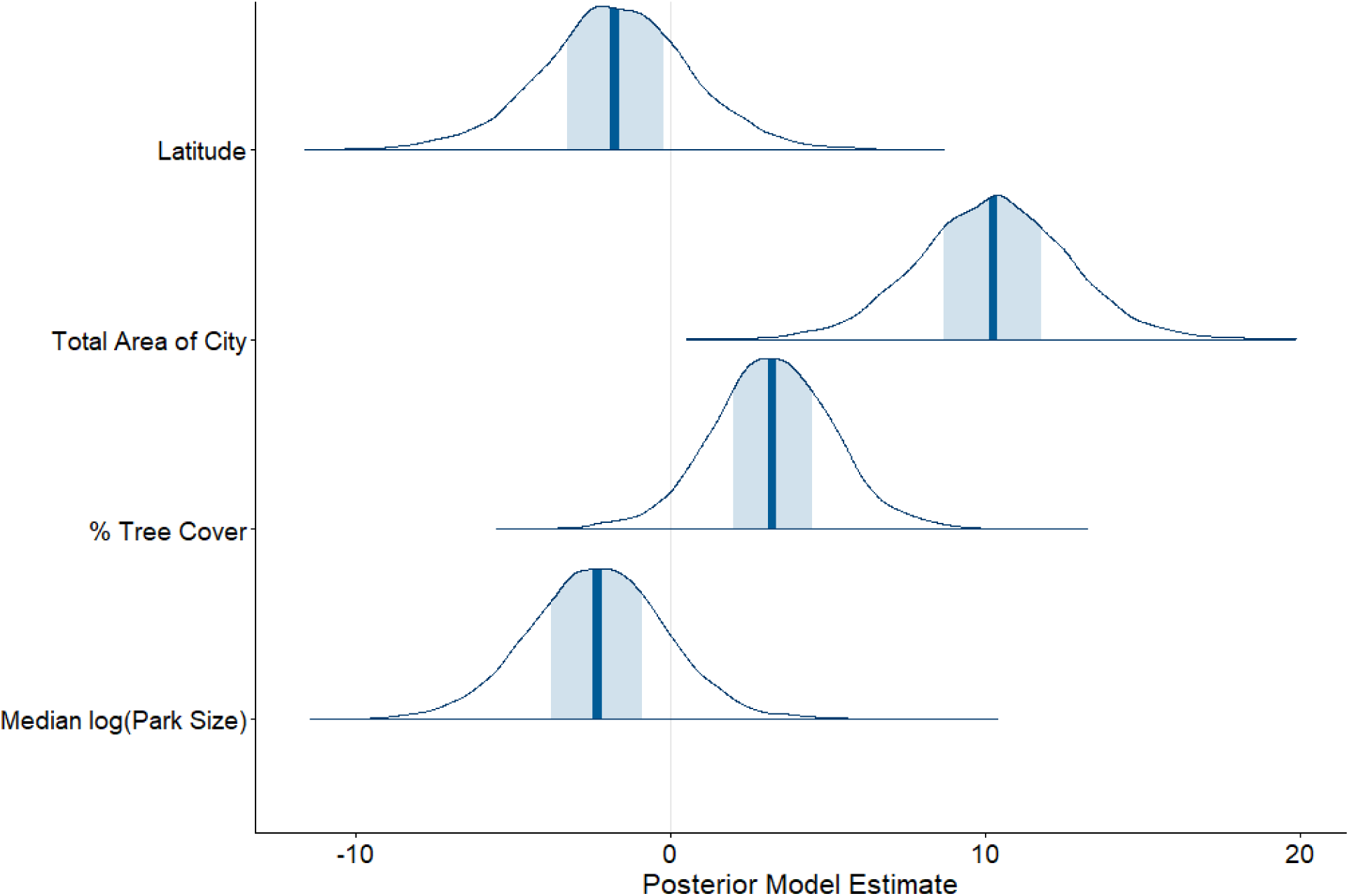
Posterior densities of parameter estimates for predictors of city-wide species richness. Dark blue bars indicate the median estimate, shaded blue areas indicate the 50% central quantiles of the posterior distributions and unshaded areas show the entire posterior distribution. In the final model, we only included parameters that had a clear directional effect on the mean value of the response (i.e., where at least the middle 50% BCI did not overlap with zero).

**Supplemental Figure S25:**
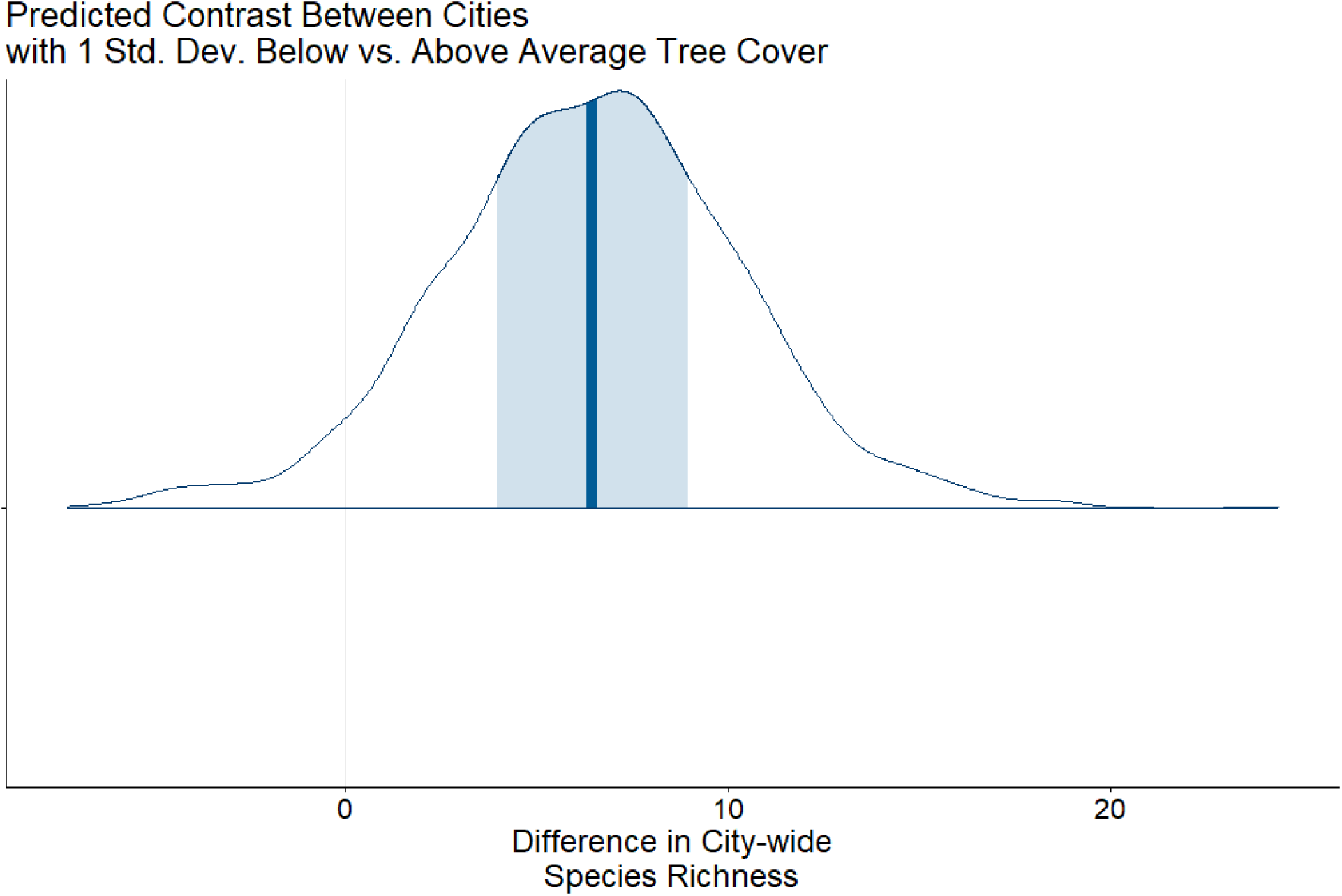
Posterior density of the predicted contrast between the predicted city-wide species richness in a city with 1 standard deviation above average tree cover versus for a city with 1 standard deviation below average tree cover. Dark blue bars indicate the median estimated contrast, shaded blue areas indicate the 50% central quantiles of the posterior distribution of contrasts and unshaded areas show the entire posterior distribution of contrasts. Values greater than zero indicate a prediction of greater city-wide species richness in a city with higher than average tree cover.

**Supplemental Figure S26:**
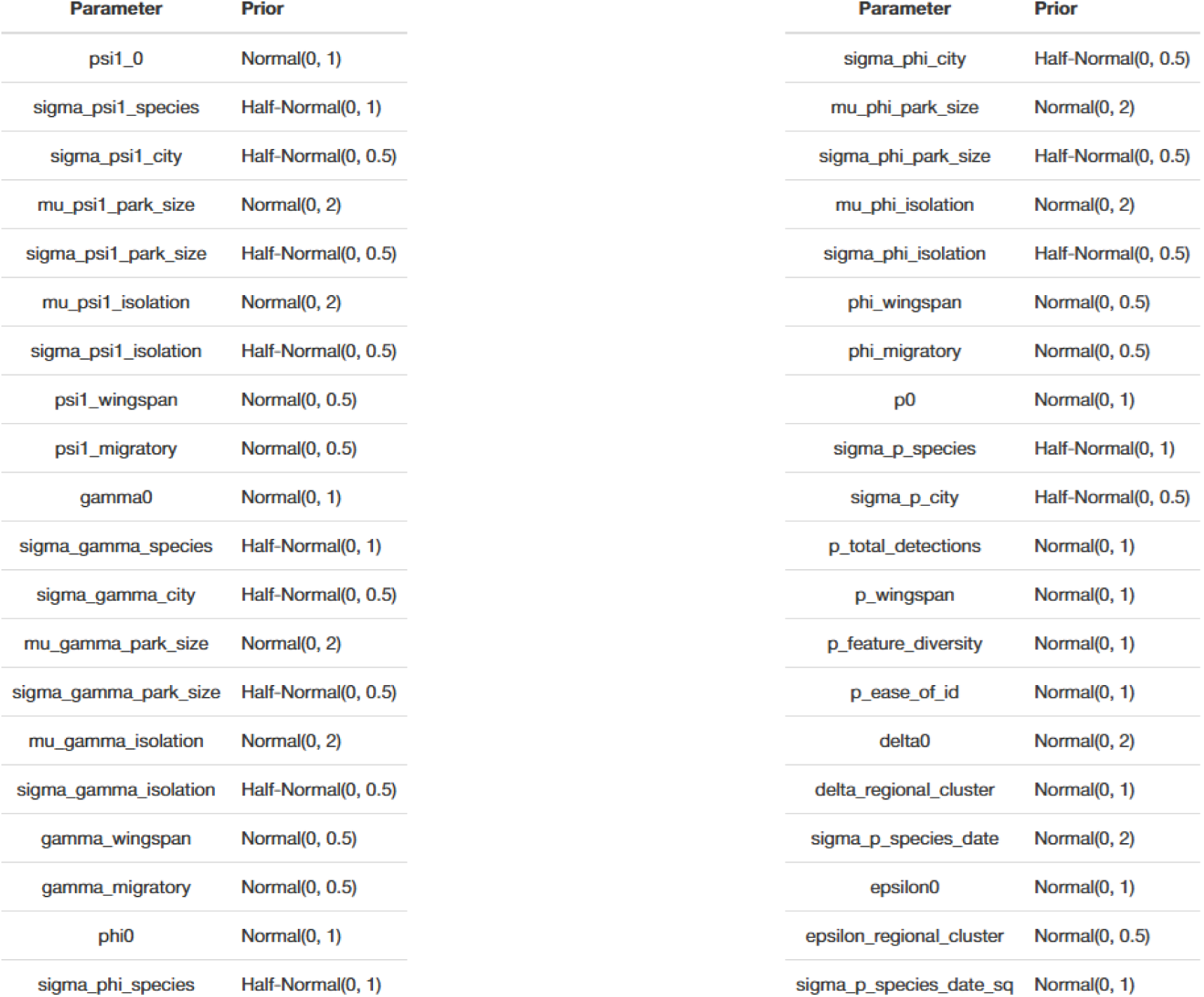
Prior probability distributions for dynamic occupancy model.

**Supplemental Figure S27:**
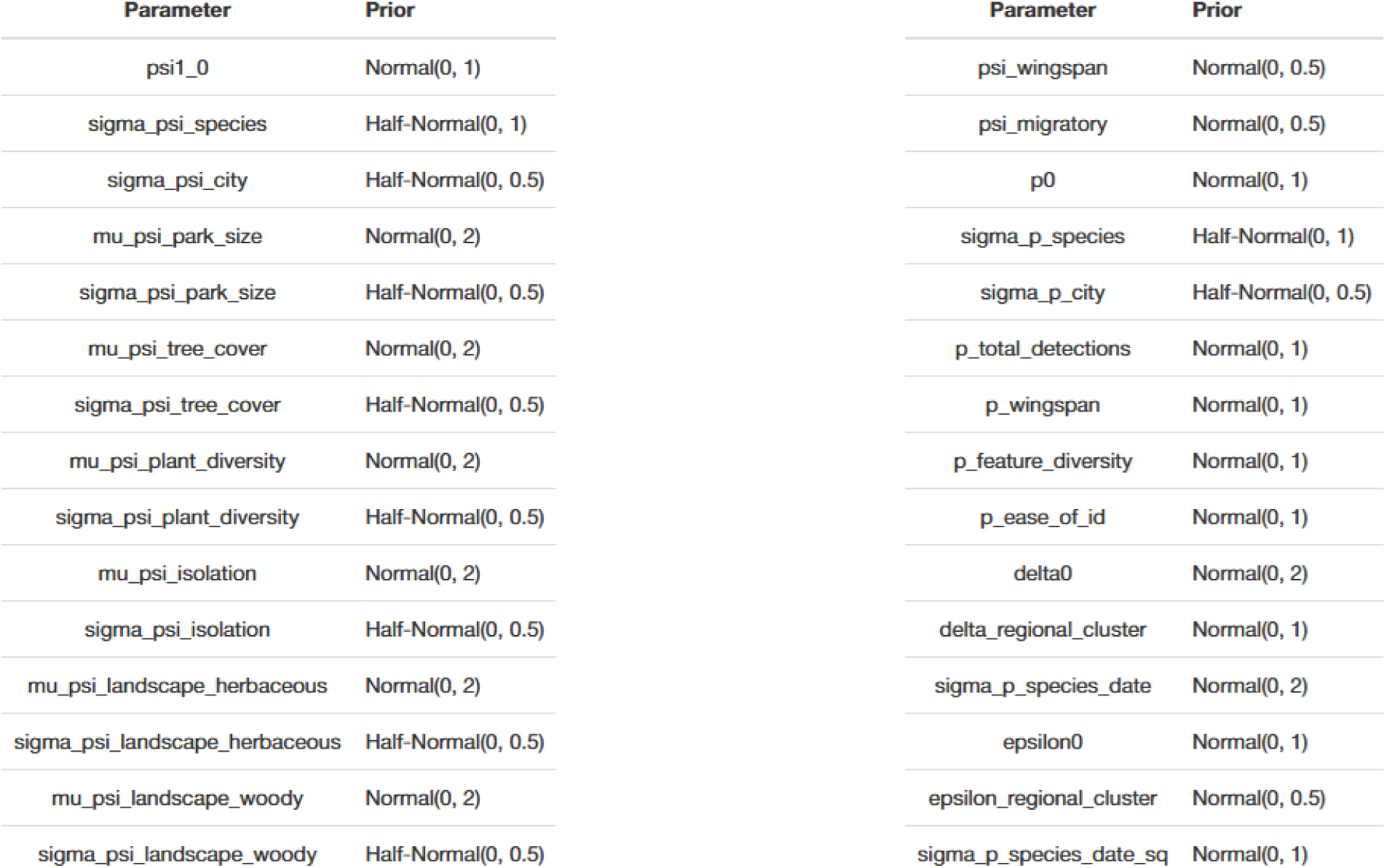
Prior probability distributions for static occupancy model.

**Supplemental Figure S28:**
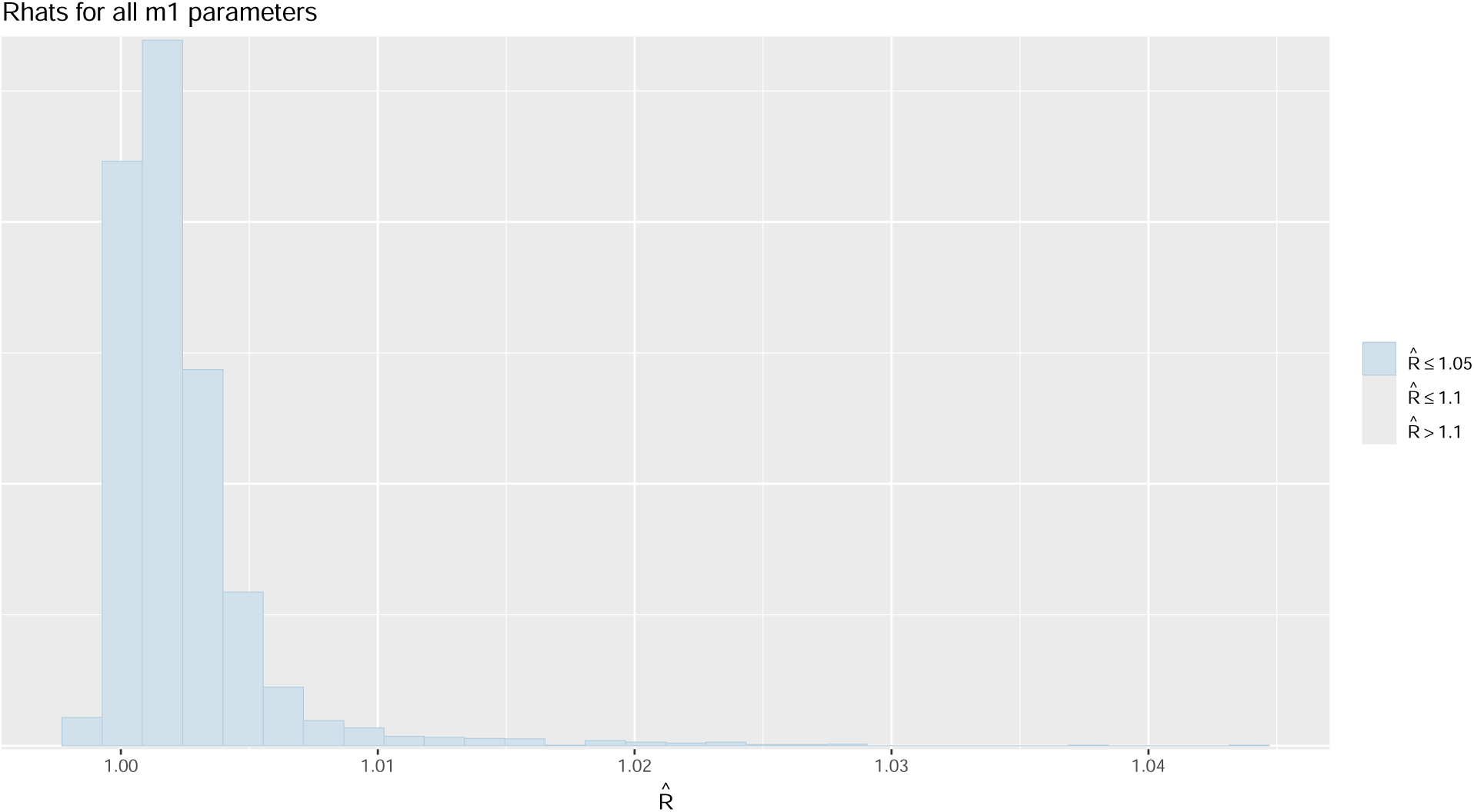
Histogram of Rhat values for all parameters for the dynamic occupancy model. Rhats less than 1.05 generally represent convergence of the HMC chains.

**Supplemental Figure S29:**
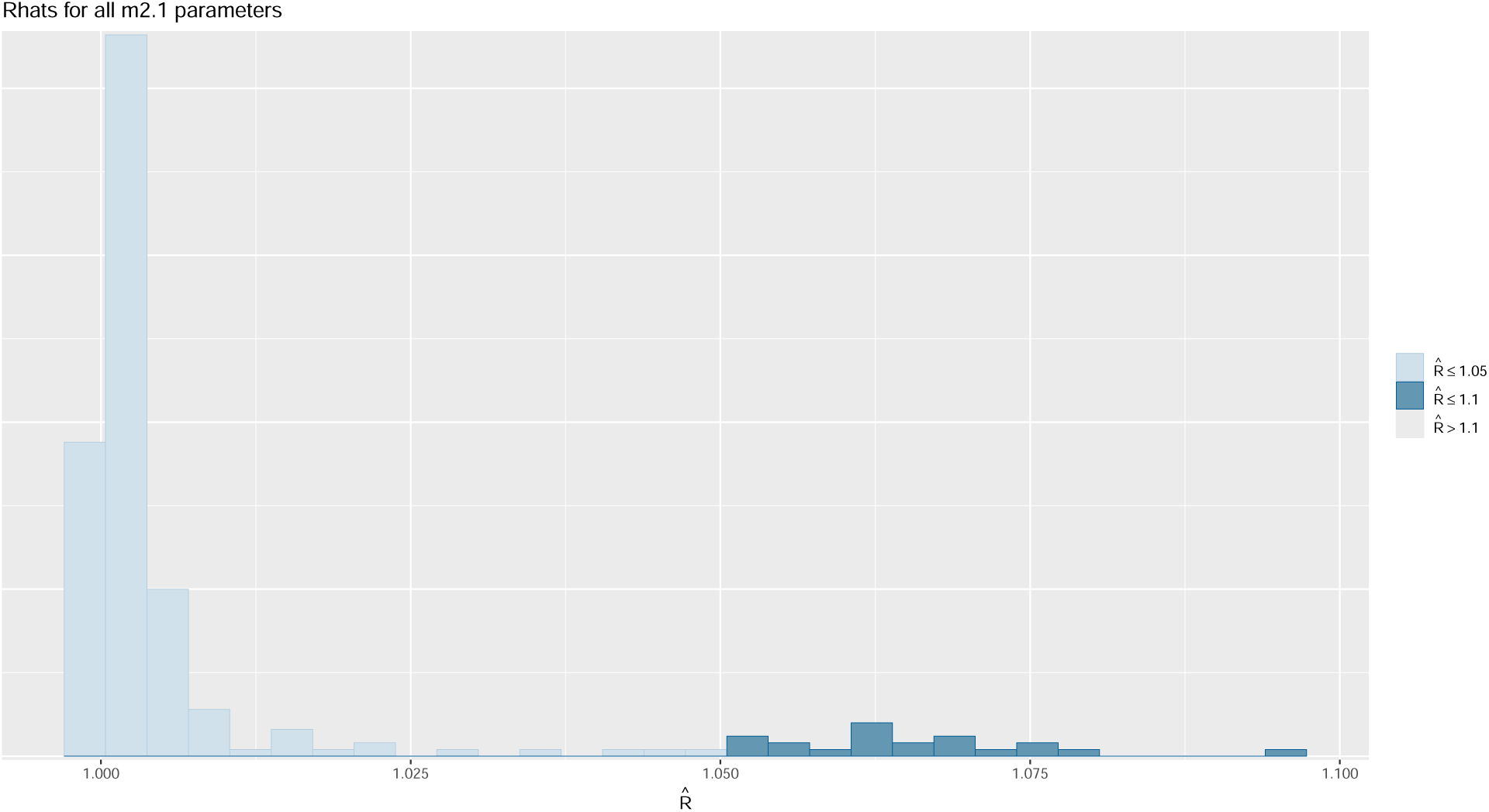
Histogram of Rhat values for all parameters for the static occupancy model. Rhats less than 1.05 generally represent convergence of the HMC chains.

**Supplemental Figure S30:**
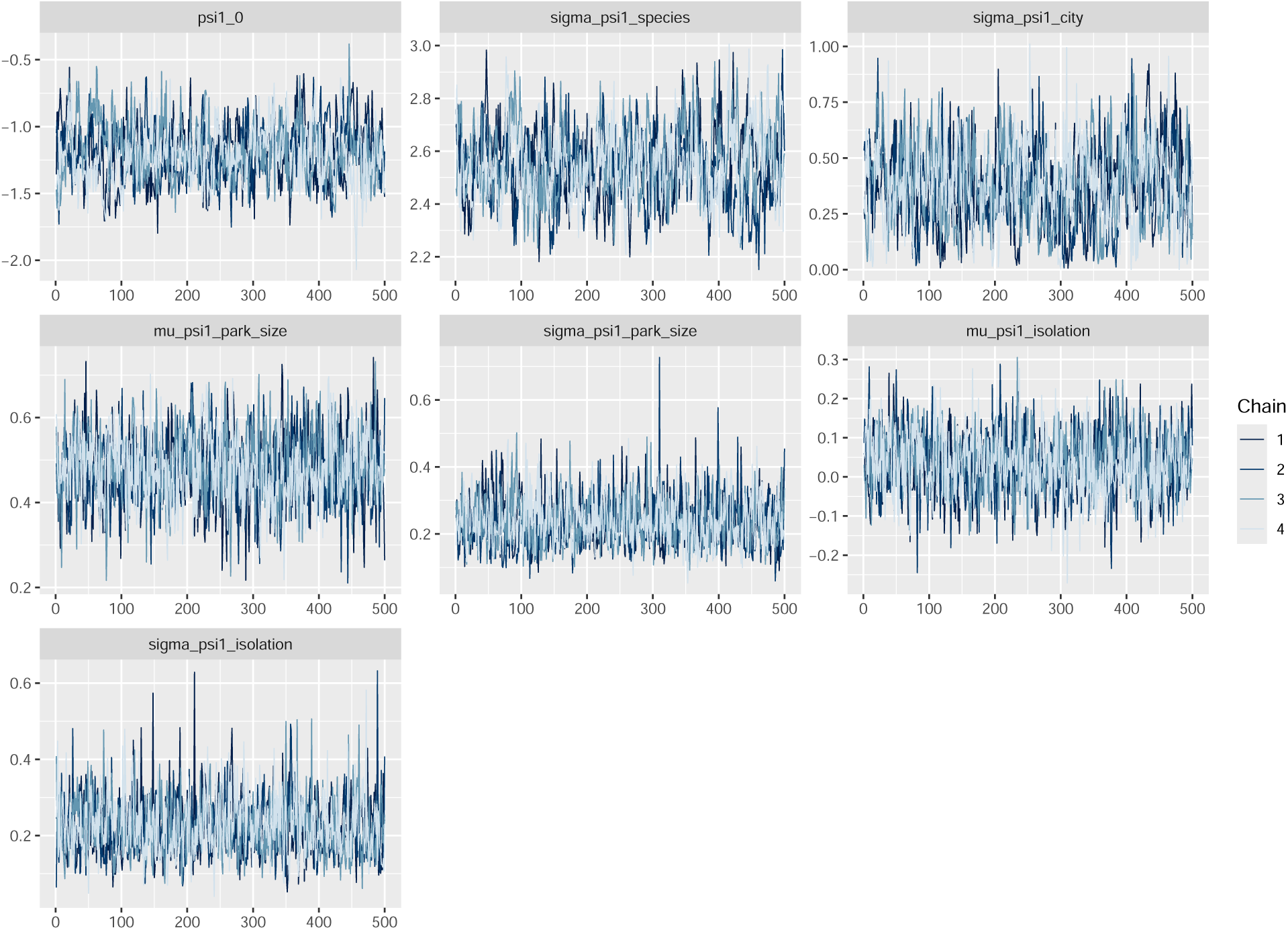
Traceplots for some sample parameters from the dynamic occupancy model. We used the traceplots to visually assess stationarity, mixing, and convergence of the HMC chains.

**Supplemental Figure S31:**
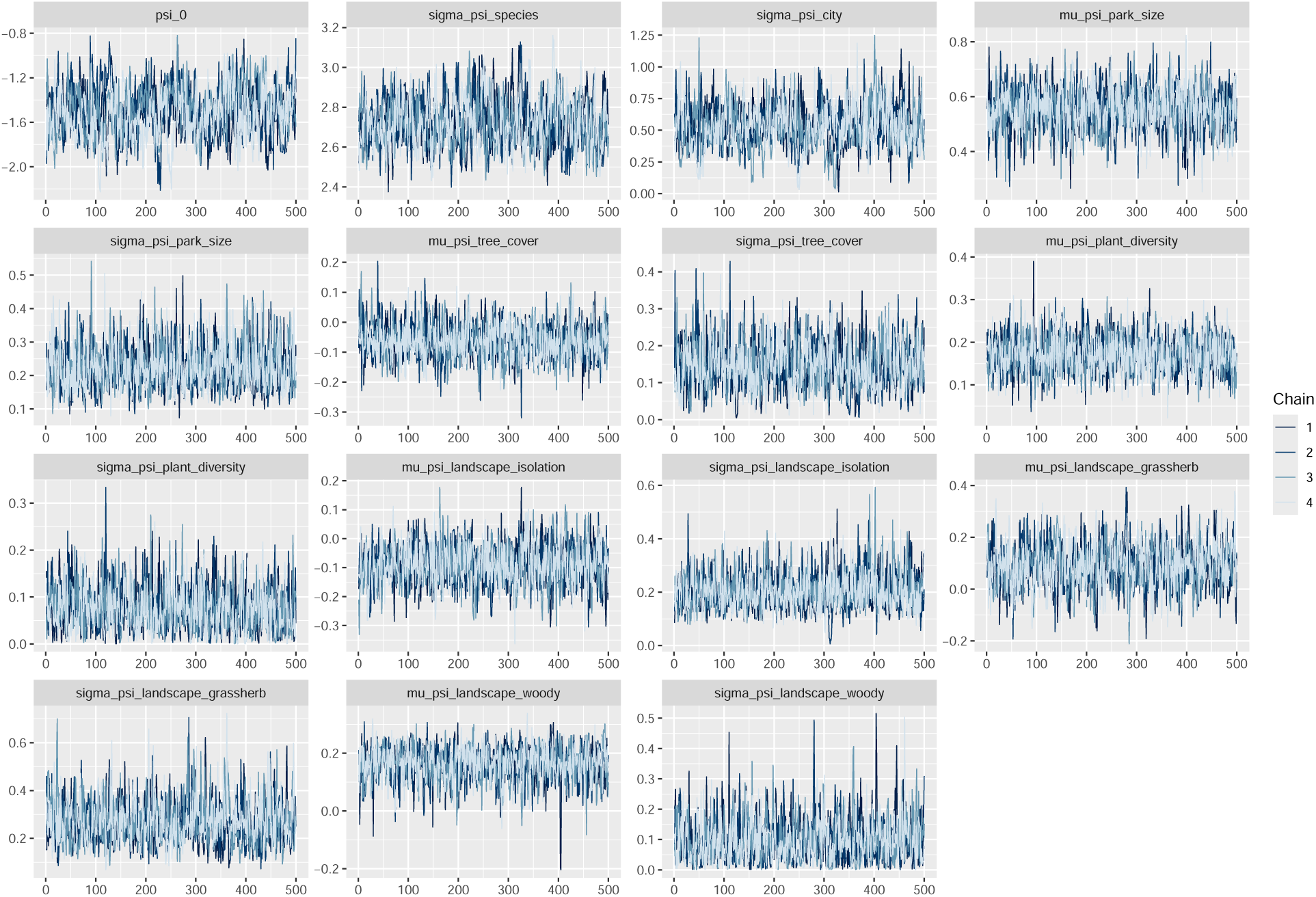
Traceplots for some sample parameters from the static occupancy model. We used the traceplots to visually assess stationarity, mixing, and convergence of the HMC chains.

**Supplemental Figure S32:**
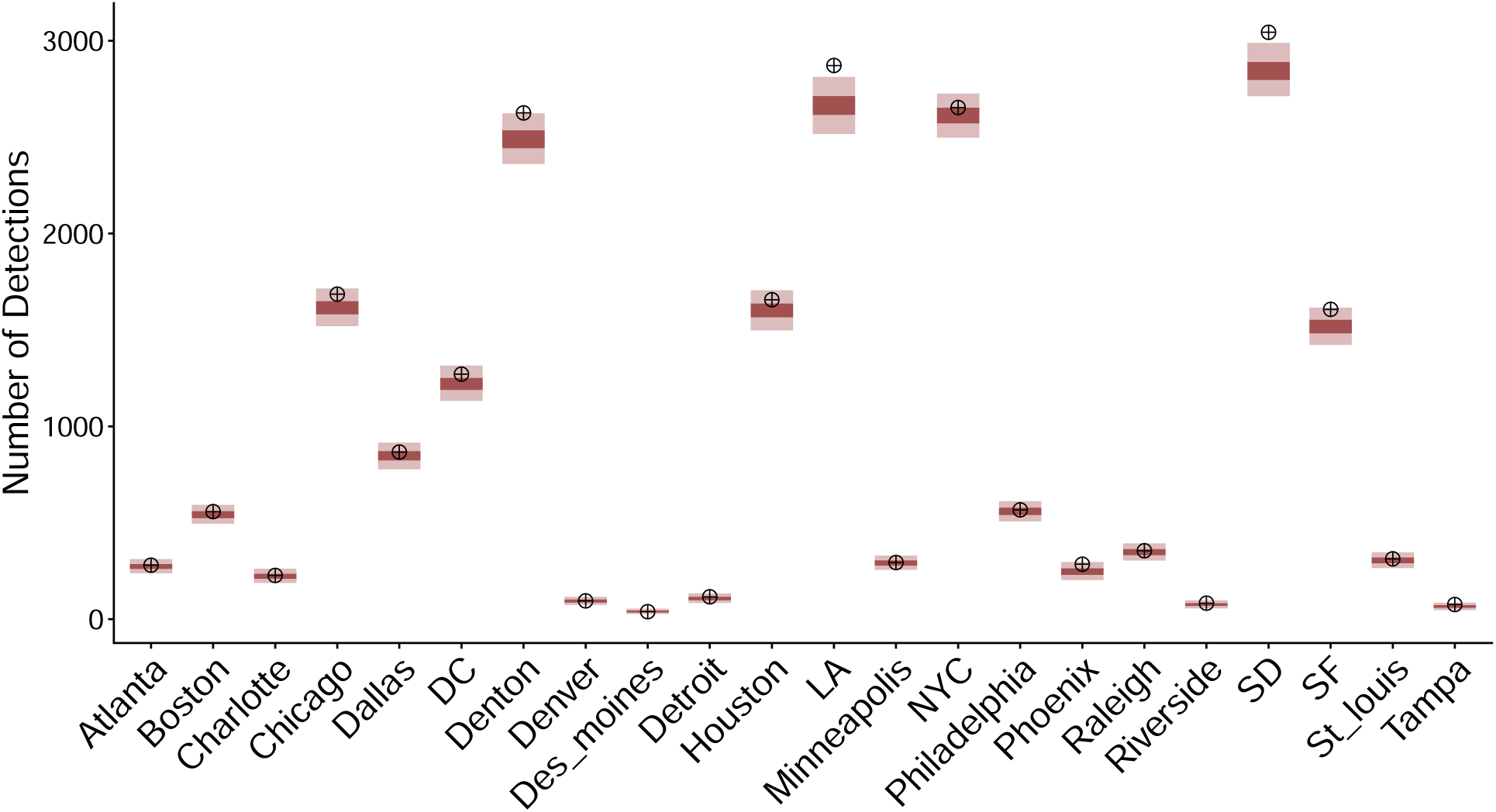
Posterior predictive checks for dynamic occupancy model. Targets represent the real number of detections observed in each city. Red 50 and 95% BCI’s for model predicted number of detections are shown in dark and light red.

**Supplemental Figure S33:**
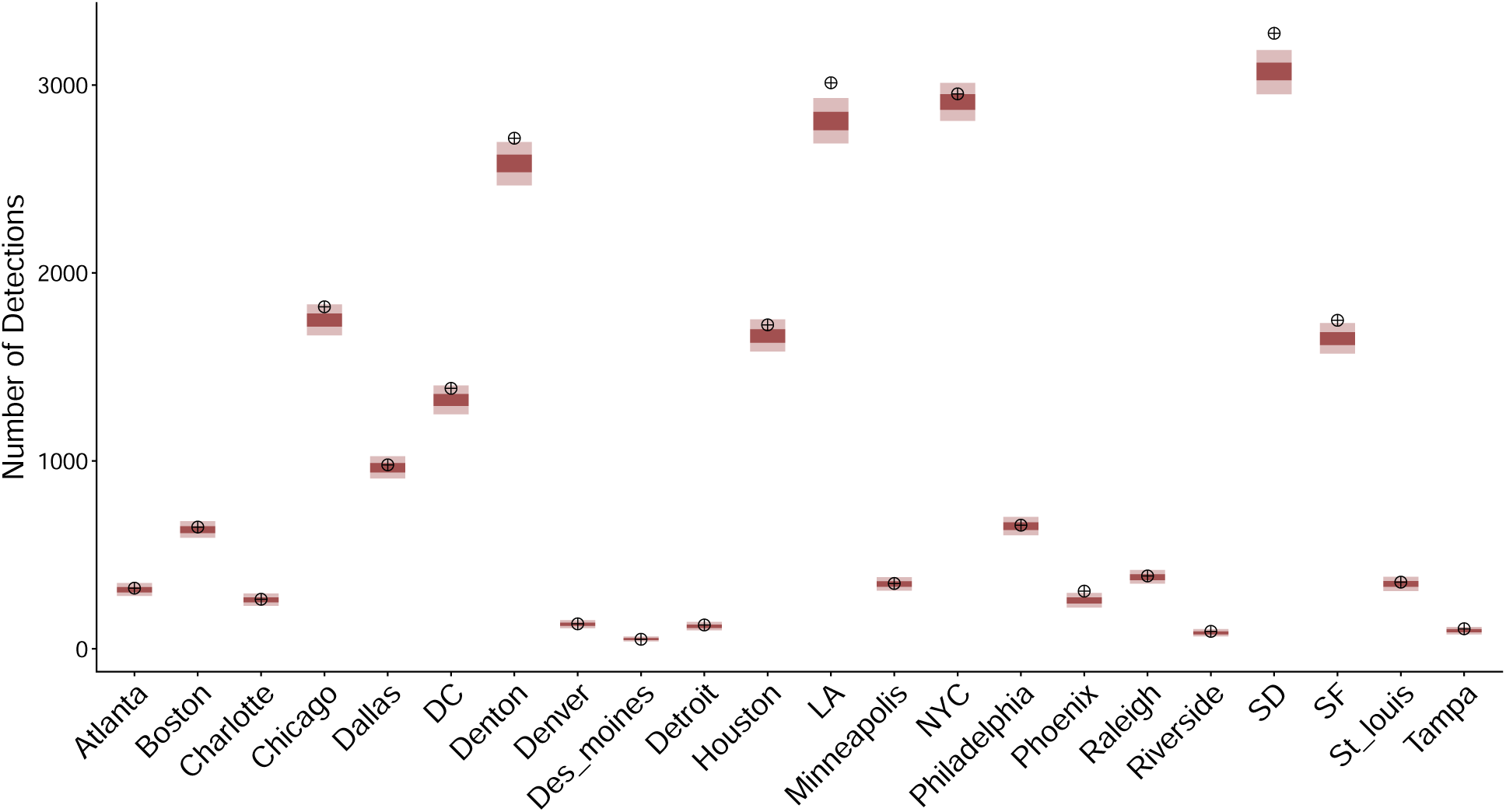
Posterior predictive checks for static occupancy model. Targets represent the real number of detections observed in each city. Red 50 and 95% BCI’s for model predicted number of detections are shown in dark and light red.

**Supplemental Table S1:** Total number of community sampling events and number of com-munity sampling events by butterfly family for each city.

| City | Community Sampling Events | Hesperiidae | Lycaenidae | Nymphalidae | Papilionidae | Pieridae |
| --- | --- | --- | --- | --- | --- | --- |
| Atlanta | 162 | 37 | 14 | 56 | 34 | 21 |
| Boston | 413 | 61 | 33 | 176 | 45 | 98 |
| Charlotte | 161 | 37 | 16 | 71 | 28 | 9 |
| Chicago | 755 | 119 | 46 | 341 | 153 | 96 |
| Washington D.C. | 449 | 85 | 52 | 141 | 90 | 81 |
| Dallas | 467 | 69 | 64 | 182 | 65 | 87 |
| Denton | 601 | 92 | 91 | 216 | 83 | 119 |
| Denver | 99 | 15 | 2 | 42 | 23 | 17 |
| Des Moines | 45 | 8 | 3 | 24 | 9 | 1 |
| Detroit | 49 | 10 | 5 | 19 | 10 | 5 |
| Houston | 414 | 75 | 52 | 150 | 79 | 58 |
| Los Angeles | 1009 | 205 | 192 | 325 | 122 | 165 |
| Minneapolis | 199 | 30 | 10 | 91 | 42 | 26 |
| New York City | 1465 | 237 | 166 | 576 | 227 | 259 |
| Philadelphia | 372 | 71 | 46 | 111 | 84 | 60 |
| Phoenix | 98 | 15 | 18 | 43 | 6 | 16 |
| Raleigh | 238 | 54 | 23 | 80 | 58 | 23 |
| Riverside | 64 | 11 | 13 | 19 | 9 | 12 |
| San Diego | 1375 | 253 | 288 | 451 | 191 | 192 |
| San Francisco | 788 | 140 | 68 | 33 | 158 | 89 |
| St. Louis | 121 | 19 | 8 | 48 | 24 | 22 |
| Tampa | 64 | 9 | 3 | 33 | 11 | 8 |

