## Supplemental Materials: Species Detections Table for "Large parks and city-wide tree cover boost butterfly diversity across 22 major U.S. cities"

**Butterfly Species Included in Occupancy Models**. Our analyses included 183 different butterfly species from five different butterfly families. Species are listed by alphabetical order within family. We modeled butterfly species occupancy within cities that were considered to be within the geographical range. We considered a city to be within range if the species was detected by iNaturalist users at least once in a park in the city, or if the species was detected by iNaturalist users at least once anywhere within a 20 km buffer of the city administrative boundary. Cities determined to be within range of each species are listed under 'Cities' column. *n* indicates the number of unique (park by month by year) detections of each species in our dataset. Duplicate detections of the same species from the same time and place were not counted. The *n* detection counts are coloured from lowest (dark colours) to highest (light colours).

| Family | Species | Cities | n |
|---|---|---|---|
| Hesperiidae | *Ancyloxypha numitor* | Atlanta, Boston, Charlotte, Chicago, Dallas, Washington D.C., Denton, Denver, Des Moines, Detroit, Houston, Minneapolis, New York City, Philadelphia, Raleigh, St. Louis, Tampa | 110 |
| Hesperiidae | *Atalopedes campestris* | Los Angeles, Riverside, San Francisco | 1 |
| Hesperiidae | *Atrytone delaware* | Boston, Charlotte, Chicago, Dallas, Washington D.C., Denton, Denver, Des Moines, Detroit, Houston, Minneapolis, New York City, St. Louis, Tampa | 12 |
| Hesperiidae | *Burnsius albezens* | Charlotte, Washington D.C., Denton, Houston, Los Angeles, Phoenix, Riverside, San Diego, Tampa | 94 |
| Hesperiidae | *Burnsius communis* | Atlanta, Charlotte, Chicago, Dallas, Washington D.C., Denton, Denver, Des Moines, Detroit, Houston, Los Angeles, Minneapolis, New York City, Philadelphia, Phoenix, Raleigh, San Francisco, St. Louis | 121 |
| Hesperiidae | *Calpodes ethlius* | Atlanta, Charlotte, Washington D.C., Denton, Houston, New York City, Raleigh, Tampa | 32 |
| Hesperiidae | *Chioides catillus* | Houston | 49 |
| Hesperiidae | *Cymaenes tripunctus* | Tampa | 1 |
| Hesperiidae | *Epargyreus clarus* | Atlanta, Boston, Charlotte, Chicago, Dallas, Washington D.C., Denton, Denver, Des Moines, Detroit, Houston, Minneapolis, New York City, Philadelphia, Raleigh, San Francisco, St. Louis, Tampa | 410 |
| Hesperiidae | *Erynnis baptisiae* | Boston, Chicago, Dallas, Washington D.C., Denton, Detroit, Houston, New York City, Philadelphia, Raleigh, St. Louis | 59 |
| Hesperiidae | *Erynnis brizo* | Washington D.C., Denton, Denver | 1 |
| Hesperiidae | *Erynnis funeralis* | Chicago, Washington D.C., Denton, Houston, Los Angeles, Minneapolis, Phoenix, Riverside, San Diego, St. Louis | 111 |

| Family | Species | Cities | n |
|---|---|---|---|
| Hesperiidae | *Erynnis horatius* | Atlanta, Boston, Charlotte, Chicago, Dallas, Washington D.C., Denton, Denver, Detroit, Houston, Minneapolis, New York City, Philadelphia, Raleigh, St. Louis, Tampa | 75 |
| Hesperiidae | *Erynnis juvenalis* | Atlanta, Boston, Charlotte, Chicago, Dallas, Washington D.C., Denton, Detroit, Houston, New York City, Philadelphia, Raleigh, St. Louis | 12 |
| Hesperiidae | *Erynnis tristis* | Washington D.C., Denton, Los Angeles, Phoenix, Riverside, San Diego, San Francisco | 78 |
| Hesperiidae | *Euphyes dion* | Charlotte, Chicago, Dallas, Washington D.C., Denton, Detroit, Minneapolis, Raleigh, St. Louis | 8 |
| Hesperiidae | *Euphyes vestris* | Atlanta, Boston, Charlotte, Chicago, Dallas, Washington D.C., Denton, Denver, Des Moines, Detroit, Houston, Minneapolis, New York City, Philadelphia, Raleigh, Riverside, San Diego, St. Louis, Tampa | 42 |
| Hesperiidae | *Heliopetes ericetorum* | Los Angeles, Phoenix, Riverside, San Diego, San Francisco | 46 |
| Hesperiidae | *Heliopetes laviana* | Houston | 1 |
| Hesperiidae | *Heliopetes macaira* | Houston | 1 |
| Hesperiidae | *Hylephila phyleus* | Atlanta, Boston, Charlotte, Chicago, Dallas, Washington D.C., Denton, Denver, Des Moines, Detroit, Houston, Los Angeles, Minneapolis, New York City, Philadelphia, Phoenix, Raleigh, Riverside, San Diego, San Francisco, St. Louis, Tampa | 1081 |
| Hesperiidae | *Lerema accius* | Atlanta, Charlotte, Dallas, Washington D.C., Denton, Houston, Raleigh, St. Louis, Tampa | 92 |
| Hesperiidae | *Lerodea arabus* | Phoenix | 1 |
| Hesperiidae | *Lerodea eufala* | Atlanta, Charlotte, Washington D.C., Denton, Houston, Los Angeles, Phoenix, Raleigh, Riverside, San Diego, San Francisco, Tampa | 103 |
| Hesperiidae | *Lon hobomok* | Boston, Chicago, Dallas, Detroit, Minneapolis, New York City, Philadelphia | 19 |
| Hesperiidae | *Lon melane* | Los Angeles, Riverside, San Diego, San Francisco | 397 |
| Hesperiidae | *Lon taxiles* | Denver | 3 |
| Hesperiidae | *Lon zabulon* | Atlanta, Boston, Charlotte, Chicago, Dallas, Washington D.C., Denton, Des Moines, Detroit, New York City, Philadelphia, Raleigh, St. Louis | 392 |
| Hesperiidae | *Mastor celia* | Washington D.C., Denton, Houston | 3 |
| Hesperiidae | *Nastra lherminier* | Atlanta, Charlotte, Dallas, Houston, New York City, Philadelphia | 1 |
| Hesperiidae | *Oarisma aurantiaca* | Washington D.C., Denton, Houston, Los Angeles, Phoenix, San Diego | 4 |
| Hesperiidae | *Oarisma minima* | Atlanta, Charlotte, Washington D.C., Denton, Houston, Raleigh, Tampa | 21 |
| Hesperiidae | *Ochlodes agricola* | Los Angeles, Riverside, San Diego, San Francisco | 12 |
| Hesperiidae | *Ochlodes sylvanoides* | Denver, Los Angeles, Riverside, San Diego, San Francisco | 118 |
| Hesperiidae | *Panoquina errans* | Los Angeles, San Diego | 33 |
| Hesperiidae | *Panoquina ocola* | Atlanta, Boston, Charlotte, Chicago, Dallas, Washington D.C., Denton, Houston, New York City, Philadelphia, Raleigh, St. Louis, Tampa | 34 |
| Hesperiidae | *Pholisora catullus* | Boston, Charlotte, Chicago, Dallas, Denver, Des Moines, Detroit, New York City, Philadelphia, Phoenix, Raleigh, St. Louis | 62 |

| Family | Species | Cities | n |
|---|---|---|---|
| Hesperiidae | *Poanes viator* | Boston, Chicago, Dallas, Washington D.C., Denton, Detroit, Houston, Minneapolis, New York City, Philadelphia, St. Louis | 39 |
| Hesperiidae | *Polites coras* | Boston, Chicago, Dallas, Denver, Des Moines, Detroit, Minneapolis, New York City, Philadelphia, St. Louis | 161 |
| Hesperiidae | *Polites egeremet* | Boston, Charlotte, Chicago, Dallas, Detroit, Houston, Minneapolis, New York City, Philadelphia, St. Louis | 3 |
| Hesperiidae | *Polites otho* | Dallas, Washington D.C., Denton, Houston, Tampa | 6 |
| Hesperiidae | *Polites sabuleti* | Los Angeles, Riverside, San Diego, San Francisco | 18 |
| Hesperiidae | *Polites themistocles* | Boston, Chicago, Dallas, Denver, Des Moines, Detroit, Minneapolis, New York City, Philadelphia, St. Louis, Tampa | 23 |
| Hesperiidae | *Polites vibex* | Atlanta, Charlotte, Washington D.C., Denton, Houston, Tampa | 2 |
| Hesperiidae | *Polygonus leo* | Phoenix | 1 |
| Hesperiidae | *Pyrgus oileus* | Atlanta, Washington D.C., Denton, Houston, Raleigh, Tampa | 17 |
| Hesperiidae | *Staphylus mazans* | Washington D.C., Denton, St. Louis | 5 |
| Hesperiidae | *Systasea zampa* | Phoenix | 1 |
| Hesperiidae | *Thorybes daunus* | Atlanta, Washington D.C., Denton, Houston, St. Louis | 1 |
| Hesperiidae | *Thorybes lyciades* | Atlanta, Charlotte, Denton, Raleigh, St. Louis | 3 |
| Hesperiidae | *Thorybes pylades* | Boston, Charlotte, Dallas, Washington D.C., Denton, Denver, Detroit, Houston, Minneapolis, New York City, Philadelphia, Raleigh, San Francisco, Tampa | 10 |
| Hesperiidae | *Thymelicus lineola* | Boston, Chicago, Des Moines, Detroit, Minneapolis | 16 |
| Hesperiidae | *Urbanus proteus* | Atlanta, Boston, Charlotte, Dallas, Washington D.C., Denton, Houston, Los Angeles, New York City, Philadelphia, Raleigh, St. Louis, Tampa | 45 |
| Hesperiidae | *Vernia verna* | Atlanta, Boston, Charlotte, Chicago, Dallas, Detroit, Houston, Minneapolis, New York City, Philadelphia, Raleigh, St. Louis | 12 |
| Lycaenidae | *Apodemia virgulti* | Los Angeles, Riverside, San Diego | 190 |
| Lycaenidae | *Atlides halesus* | Atlanta, Charlotte, Washington D.C., Denton, Houston, Los Angeles, Phoenix, Raleigh, Riverside, San Diego, Tampa | 39 |
| Lycaenidae | *Brephidium exilis* | Washington D.C., Denton, Denver, Houston, Los Angeles, Phoenix, Riverside, San Diego, San Francisco, Tampa | 190 |
| Lycaenidae | *Callicista columella* | Washington D.C., Denton, Houston, Los Angeles, Phoenix, Riverside, San Diego, Tampa | 13 |
| Lycaenidae | *Callophrys dumetorum* | Los Angeles, Riverside, San Diego, San Francisco | 27 |
| Lycaenidae | *Callophrys viridis* | San Francisco | 19 |
| Lycaenidae | *Calycopis cecrops* | Atlanta, Boston, Charlotte, Dallas, Washington D.C., Denton, Houston, New York City, Philadelphia, Raleigh, St. Louis, Tampa | 159 |
| Lycaenidae | *Calycopis isobeon* | Washington D.C., Denton, Houston | 14 |
| Lycaenidae | *Celastrina ladon* | Atlanta, Boston, Charlotte, Chicago, Dallas, Denver, Des Moines, Los Angeles, New York City, Philadelphia, Phoenix, Raleigh, Riverside, San Diego, San Francisco, St. Louis | 54 |

| Family | Species | Cities | n |
|---|---|---|---|
| Lycaenidae | *Cyaniris neglecta* | Atlanta, Boston, Charlotte, Chicago, Dallas, Washington D.C., Denton, Des Moines, Detroit, Minneapolis, New York City, Philadelphia, Raleigh, St. Louis | 156 |
| Lycaenidae | *Echinargus isola* | Washington D.C., Denton, Denver, Des Moines, Houston, Los Angeles, Phoenix, Riverside, San Diego | 73 |
| Lycaenidae | *Elkalyce amyntula* | Denver, Los Angeles, San Diego, San Francisco | 20 |
| Lycaenidae | *Elkalyce comyntas* | Atlanta, Boston, Charlotte, Chicago, Dallas, Washington D.C., Denton, Des Moines, Detroit, Houston, Minneapolis, New York City, Philadelphia, Raleigh, San Francisco, St. Louis | 261 |
| Lycaenidae | *Eumaeus atala* | Tampa | 1 |
| Lycaenidae | *Feniseca tarquinius* | Atlanta, Charlotte, Chicago, Dallas, Detroit, Minneapolis, New York City, Philadelphia, Raleigh, St. Louis | 21 |
| Lycaenidae | *Fixsenia favonius* | Atlanta, Dallas, Washington D.C., Denton, Houston, New York City, Philadelphia, Tampa | 5 |
| Lycaenidae | *Glaucopsyche lygdamus* | Denver, Los Angeles, Minneapolis, Riverside, San Diego, San Francisco | 43 |
| Lycaenidae | *Harkenclenus titus* | Boston, Chicago, Denver, Des Moines, Detroit, Minneapolis, New York City, Philadelphia | 1 |
| Lycaenidae | *Hemiargus ceraunus* | Washington D.C., Denton, Houston, Los Angeles, Phoenix, Riverside, San Diego, Tampa | 98 |
| Lycaenidae | *Icaricia acmon* | Los Angeles, Phoenix, Riverside, San Diego, San Francisco | 126 |
| Lycaenidae | *Icaricia lupini* | Denver, Los Angeles, Phoenix, San Diego | 1 |
| Lycaenidae | *Incisalia henrici* | Boston, Dallas, Washington D.C., Denton, Philadelphia, St. Louis, Tampa | 1 |
| Lycaenidae | *Incisalia irioides* | Boston, Denver, Los Angeles, Phoenix, Riverside, San Diego, San Francisco | 18 |
| Lycaenidae | *Leptotes marina* | Chicago, Washington D.C., Denton, Denver, Los Angeles, Phoenix, Riverside, San Diego | 382 |
| Lycaenidae | *Lycaena hypophlaeas* | Boston, Detroit, Minneapolis, New York City, Philadelphia | 38 |
| Lycaenidae | *Ministrymon leda* | Los Angeles, Phoenix, San Diego | 7 |
| Lycaenidae | *Mitoura gryneus* | Atlanta, Boston, Charlotte, Dallas, Washington D.C., Denton, Denver, Minneapolis, Philadelphia, Phoenix, Raleigh | 5 |
| Lycaenidae | *Parrhasius m-album* | Atlanta, Boston, Charlotte, Chicago, Dallas, Washington D.C., Denton, Houston, New York City, Philadelphia, Raleigh, St. Louis, Tampa | 13 |
| Lycaenidae | *Phaeostrymon alcestis* | Washington D.C., Denton | 2 |
| Lycaenidae | *Philotes bernardino* | Los Angeles, Riverside, San Diego | 49 |
| Lycaenidae | *Satyrium calanus* | Atlanta, Boston, Charlotte, Chicago, Dallas, Washington D.C., Denton, Denver, Detroit, Houston, Los Angeles, Minneapolis, New York City, Philadelphia, Raleigh, St. Louis | 28 |
| Lycaenidae | *Satyrium edwardsii* | Boston, Detroit, Minneapolis | 1 |

| Family | Species | Cities | n |
|---|---|---|---|
| Lycaenidae | *Strymon melinus* | Atlanta, Boston, Charlotte, Chicago, Dallas, Washington D.C., Denton, Denver, Des Moines, Houston, Los Angeles, New York City, Philadelphia, Phoenix, Raleigh, Riverside, San Diego, San Francisco, St. Louis, Tampa | 641 |
| Lycaenidae | *Strymon saepium* | Denver, Los Angeles, Riverside, San Diego, San Francisco | 10 |
| Lycaenidae | *Strymon sylvinus* | Los Angeles, Riverside, San Diego | 43 |
| Lycaenidae | *Tharsalea hyllus* | Chicago, Denver, Des Moines, Detroit, Minneapolis, New York City, St. Louis | 7 |
| Lycaenidae | *Thecla iroides* | Los Angeles, San Diego, San Francisco | 2 |
| Nymphalidae | *Anaea andria* | Washington D.C., Denton, Denver, Houston, St. Louis | 77 |
| Nymphalidae | *Anartia jatrophae* | Boston, Denton, Tampa | 13 |
| Nymphalidae | *Anthanassa texana* | Washington D.C., Denton, Houston, Phoenix | 32 |
| Nymphalidae | *Asterocampa celtis* | Atlanta, Charlotte, Chicago, Dallas, Washington D.C., Denton, Denver, Des Moines, Detroit, Houston, Minneapolis, New York City, Philadelphia, Raleigh, St. Louis, Tampa | 294 |
| Nymphalidae | *Asterocampa clyton* | Atlanta, Charlotte, Chicago, Dallas, Washington D.C., Denton, Detroit, Houston, Minneapolis, New York City, Philadelphia, Raleigh, St. Louis, Tampa | 119 |
| Nymphalidae | *Asterocampa leilia* | Phoenix | 6 |
| Nymphalidae | *Cercyonis pegala* | Atlanta, Boston, Charlotte, Chicago, Dallas, Washington D.C., Denton, Denver, Des Moines, Detroit, Minneapolis, New York City, Philadelphia, Raleigh, San Francisco, St. Louis | 8 |
| Nymphalidae | *Chlosyne californica* | Los Angeles, Phoenix, Riverside, San Diego | 7 |
| Nymphalidae | *Chlosyne gabbii* | Los Angeles, Riverside, San Diego | 10 |
| Nymphalidae | *Chlosyne gorgone* | Washington D.C., Denton, Denver, Des Moines | 2 |
| Nymphalidae | *Chlosyne lacinia* | Washington D.C., Denton, Phoenix | 12 |
| Nymphalidae | *Chlosyne nycteis* | Atlanta, Charlotte, Chicago, Dallas, Washington D.C., Denton, Des Moines, Detroit, Houston, Minneapolis, New York City, Raleigh, St. Louis | 9 |
| Nymphalidae | *Coenonympha california* | Boston, Denver, Detroit, Los Angeles, Minneapolis, New York City, San Diego, San Francisco | 44 |
| Nymphalidae | *Danaus gilippus* | Charlotte, Dallas, Washington D.C., Denton, Denver, Houston, Los Angeles, Phoenix, Riverside, San Diego, San Francisco, Tampa | 238 |
| Nymphalidae | *Danaus plexippus* | Atlanta, Boston, Charlotte, Chicago, Dallas, Washington D.C., Denton, Denver, Des Moines, Detroit, Houston, Los Angeles, Minneapolis, New York City, Philadelphia, Phoenix, Raleigh, Riverside, San Diego, San Francisco, St. Louis, Tampa | 2290 |
| Nymphalidae | *Dione vanillae* | Atlanta, Charlotte, Washington D.C., Denton, Denver, Houston, Los Angeles, New York City, Phoenix, Raleigh, Riverside, San Diego, San Francisco, St. Louis, Tampa | 728 |
| Nymphalidae | *Enodia portlandia* | Charlotte, Houston, Raleigh | 1 |
| Nymphalidae | *Eresia aveyrona* | Phoenix, Riverside, San Diego, San Francisco | 7 |
| Nymphalidae | *Euphydryas phaeton* | Boston, Chicago, Detroit, Minneapolis | 1 |

| Family | Species | Cities | n |
|---|---|---|---|
| Nymphalidae | *Euptoieta claudia* | Atlanta, Boston, Charlotte, Chicago, Dallas, Washington D.C., Denton, Denver, Des Moines, Detroit, Houston, Los Angeles, Minneapolis, New York City, Philadelphia, Phoenix, Raleigh, Riverside, San Diego, St. Louis, Tampa | 350 |
| Nymphalidae | *Euptychia cornelius* | Atlanta, Charlotte, Washington D.C., Denton, Houston, Raleigh, St. Louis | 2 |
| Nymphalidae | *Euptychia cymela* | Atlanta, Boston, Charlotte, Chicago, Dallas, Washington D.C., Denton, Des Moines, Detroit, Houston, Minneapolis, New York City, Philadelphia, Raleigh, St. Louis | 48 |
| Nymphalidae | *Heliconius charithonia* | Atlanta, Charlotte, Washington D.C., Denton, Houston, Los Angeles, Phoenix, Tampa | 13 |
| Nymphalidae | *Hermeuptychia hermes* | Atlanta, Charlotte, Houston, Raleigh, Tampa | 10 |
| Nymphalidae | *Junonia coenia* | Atlanta, Boston, Charlotte, Chicago, Dallas, Washington D.C., Denton, Denver, Des Moines, Detroit, Houston, Minneapolis, New York City, Philadelphia, Raleigh, St. Louis, Tampa | 677 |
| Nymphalidae | *Junonia grisea* | Denver, Los Angeles, Phoenix, Riverside, San Diego, San Francisco | 280 |
| Nymphalidae | *Junonia neildi* | Tampa | 2 |
| Nymphalidae | *Lethe anthedon* | Atlanta, Boston, Charlotte, Chicago, Dallas, Des Moines, Detroit, Minneapolis, Raleigh, St. Louis | 15 |
| Nymphalidae | *Lethe creola* | Atlanta, Charlotte, Raleigh | 3 |
| Nymphalidae | *Lethe eurydice* | Atlanta, Boston, Charlotte, Chicago, Dallas, Detroit, Minneapolis, New York City, Philadelphia, Raleigh, St. Louis | 28 |
| Nymphalidae | *Libytheana carinenta* | Atlanta, Boston, Charlotte, Chicago, Dallas, Washington D.C., Denton, Denver, Des Moines, Detroit, Houston, Los Angeles, Minneapolis, New York City, Philadelphia, Phoenix, Raleigh, Riverside, San Diego, St. Louis | 180 |
| Nymphalidae | *Limenitis archippus* | Atlanta, Boston, Charlotte, Chicago, Dallas, Washington D.C., Denton, Denver, Des Moines, Detroit, Houston, Los Angeles, Minneapolis, New York City, Philadelphia, Phoenix, Raleigh, St. Louis, Tampa | 129 |
| Nymphalidae | *Limenitis arthemis* | Atlanta, Boston, Charlotte, Chicago, Dallas, Washington D.C., Denton, Des Moines, Detroit, Houston, Minneapolis, New York City, Philadelphia, Phoenix, Raleigh, St. Louis, Tampa | 76 |
| Nymphalidae | *Limenitis astyanax* | Atlanta, Boston, Charlotte, Chicago, Dallas, Washington D.C., Denton, Des Moines, Detroit, Houston, Minneapolis, New York City, Philadelphia, Raleigh, St. Louis, Tampa | 159 |
| Nymphalidae | *Limenitis bredowii* | Los Angeles, Phoenix, Riverside, San Diego, San Francisco | 83 |
| Nymphalidae | *Limenitis lorquini* | Los Angeles, Riverside, San Diego, San Francisco | 178 |
| Nymphalidae | *Mestra amymone* | Washington D.C., Denton | 2 |
| Nymphalidae | *Nymphalis antiopa* | Atlanta, Boston, Charlotte, Chicago, Dallas, Washington D.C., Denton, Denver, Des Moines, Detroit, Los Angeles, Minneapolis, New York City, Philadelphia, Phoenix, Raleigh, Riverside, San Diego, San Francisco, St. Louis | 484 |
| Nymphalidae | *Nymphalis californica* | Denver, Los Angeles, Riverside, San Diego, San Francisco | 21 |

| Family | Species | Cities | n |
|---|---|---|---|
| Nymphalidae | *Occidryas chalcedona* | Los Angeles, Phoenix, Riverside, San Diego, San Francisco | 93 |
| Nymphalidae | *Phyciodes phaon* | Washington D.C., Denton, Houston, Tampa | 132 |
| Nymphalidae | *Phyciodes tharos* | Atlanta, Boston, Charlotte, Chicago, Dallas, Washington D.C., Denton, Denver, Des Moines, Detroit, Houston, Minneapolis, New York City, Philadelphia, Raleigh, San Francisco, St. Louis, Tampa | 561 |
| Nymphalidae | *Polygonia comma* | Atlanta, Boston, Charlotte, Chicago, Dallas, Washington D.C., Denton, Des Moines, Detroit, Minneapolis, New York City, Philadelphia, Raleigh, St. Louis | 110 |
| Nymphalidae | *Polygonia interrogationis* | Atlanta, Boston, Charlotte, Chicago, Dallas, Washington D.C., Denton, Denver, Des Moines, Detroit, Houston, Los Angeles, Minneapolis, New York City, Philadelphia, Phoenix, Raleigh, St. Louis, Tampa | 459 |
| Nymphalidae | *Polygonia progne* | Chicago, Des Moines, Minneapolis | 1 |
| Nymphalidae | *Polygonia satyrus* | Denver, Los Angeles, Phoenix, Riverside, San Diego, San Francisco | 8 |
| Nymphalidae | *Polygonia vaualbum* | Boston, Chicago, Minneapolis | 3 |
| Nymphalidae | *Speyeria aphrodite* | Denver, Minneapolis, Philadelphia | 1 |
| Nymphalidae | *Speyeria callippe* | Denver, Los Angeles, San Diego, San Francisco | 3 |
| Nymphalidae | *Speyeria cybele* | Atlanta, Boston, Charlotte, Chicago, Dallas, Denton, Des Moines, Detroit, Minneapolis, New York City, Philadelphia, Raleigh, St. Louis | 28 |
| Nymphalidae | *Speyeria idalia* | Denver, Des Moines | 1 |
| Nymphalidae | *Vanessa annabella* | Washington D.C., Denton, Denver, Houston, Los Angeles, Phoenix, Riverside, San Diego, San Francisco | 379 |
| Nymphalidae | *Vanessa atalanta* | Atlanta, Boston, Charlotte, Chicago, Dallas, Washington D.C., Denton, Denver, Des Moines, Detroit, Houston, Los Angeles, Minneapolis, New York City, Philadelphia, Phoenix, Raleigh, Riverside, San Diego, San Francisco, St. Louis, Tampa | 1395 |
| Nymphalidae | *Vanessa cardui* | Atlanta, Boston, Charlotte, Chicago, Dallas, Washington D.C., Denton, Denver, Des Moines, Detroit, Houston, Los Angeles, Minneapolis, New York City, Philadelphia, Phoenix, Raleigh, Riverside, San Diego, San Francisco, St. Louis, Tampa | 587 |
| Nymphalidae | *Vanessa virginiensis* | Atlanta, Boston, Charlotte, Chicago, Dallas, Washington D.C., Denton, Denver, Des Moines, Detroit, Houston, Los Angeles, Minneapolis, New York City, Philadelphia, Phoenix, Raleigh, Riverside, San Diego, San Francisco, St. Louis, Tampa | 505 |
| Papilionidae | *Battus philenor* | Atlanta, Boston, Charlotte, Chicago, Dallas, Washington D.C., Denton, Denver, Detroit, Houston, Los Angeles, New York City, Philadelphia, Phoenix, Raleigh, San Diego, San Francisco, St. Louis, Tampa | 290 |
| Papilionidae | *Battus polydamas* | Houston, San Diego, Tampa | 2 |
| Papilionidae | *Papilio canadensis* | Boston | 1 |
| Papilionidae | *Papilio cresphontes* | Atlanta, Boston, Charlotte, Chicago, Dallas, Washington D.C., Denton, Des Moines, Detroit, Houston, Minneapolis, New York City, Philadelphia, St. Louis, Tampa | 86 |

| Family | Species | Cities | n |
|---|---|---|---|
| Papilionidae | *Papilio eurymedon* | Denver, Los Angeles, Riverside, San Diego, San Francisco | 69 |
| Papilionidae | *Papilio glaucus* | Atlanta, Boston, Charlotte, Chicago, Dallas, Washington D.C., Denton, Denver, Des Moines, Detroit, Houston, Minneapolis, New York City, Philadelphia, Raleigh, St. Louis, Tampa | 644 |
| Papilionidae | *Papilio multicaudata* | Denver, Phoenix | 19 |
| Papilionidae | *Papilio palamedes* | Houston, Raleigh, Tampa | 3 |
| Papilionidae | *Papilio polyxenes* | Atlanta, Boston, Charlotte, Chicago, Dallas, Washington D.C., Denton, Denver, Des Moines, Detroit, Houston, Los Angeles, Minneapolis, New York City, Philadelphia, Phoenix, Raleigh, Riverside, San Diego, St. Louis, Tampa | 771 |
| Papilionidae | *Papilio rumiko* | Washington D.C., Denton, Houston, Los Angeles, Phoenix, Riverside, San Diego | 143 |
| Papilionidae | *Papilio rutulus* | Denver, Los Angeles, Phoenix, Riverside, San Diego, San Francisco | 287 |
| Papilionidae | *Papilio troilus* | Atlanta, Boston, Charlotte, Chicago, Dallas, Washington D.C., Denton, Detroit, Houston, New York City, Philadelphia, Raleigh, St. Louis, Tampa | 234 |
| Papilionidae | *Papilio zelicaon* | Denver, Los Angeles, Riverside, San Diego, San Francisco | 317 |
| Papilionidae | *Protographium marcellus* | Atlanta, Charlotte, Chicago, Dallas, Houston, Raleigh, St. Louis, Tampa | 18 |
| Pieridae | *Abaeis mexicana* | Washington D.C., Denton, Denver, Houston, Phoenix | 1 |
| Pieridae | *Abaeis nicippe* | Atlanta, Charlotte, Chicago, Dallas, Washington D.C., Denton, Denver, Houston, Los Angeles, Philadelphia, Phoenix, Raleigh, Riverside, San Diego, St. Louis, Tampa | 77 |
| Pieridae | *Anteos clorinde* | Houston | 1 |
| Pieridae | *Anteos maerula* | Houston | 1 |
| Pieridae | *Anthocharis cethura* | Phoenix, Riverside, San Diego | 3 |
| Pieridae | *Anthocharis midea* | Atlanta, Charlotte, Dallas, Washington D.C., Denton, Houston, Philadelphia, Raleigh | 6 |
| Pieridae | *Anthocharis sara* | Los Angeles, Riverside, San Diego, San Francisco | 86 |
| Pieridae | *Ascia monuste* | Denton, Houston, Phoenix, Tampa | 2 |
| Pieridae | *Colias eurytheme* | Atlanta, Boston, Charlotte, Chicago, Dallas, Washington D.C., Denton, Denver, Des Moines, Detroit, Houston, Los Angeles, Minneapolis, New York City, Philadelphia, Phoenix, Raleigh, Riverside, San Diego, San Francisco, St. Louis | 446 |
| Pieridae | *Colias philodice* | Boston, Charlotte, Chicago, Dallas, Denver, Des Moines, Detroit, Minneapolis, New York City, Philadelphia, Raleigh, St. Louis | 67 |
| Pieridae | *Euchloe ausonides* | Denver, San Francisco | 1 |
| Pieridae | *Eurema daira* | Tampa | 3 |
| Pieridae | *Kricogonia lyside* | Washington D.C., Denton, Houston, Phoenix | 1 |

| Family | Species | Cities | n |
|---|---|---|---|
| Pieridae | *Nathalis iole* | Washington D.C., Denton, Denver, Des Moines, Houston, Los Angeles, Minneapolis, New York City, Phoenix, Riverside, San Diego, St. Louis, Tampa | 143 |
| Pieridae | *Phoebis agarithe* | Washington D.C., Denton, Houston, Los Angeles, Phoenix, Riverside, San Diego | 13 |
| Pieridae | *Phoebis marcellina* | Denver, Los Angeles, Riverside, San Diego | 6 |
| Pieridae | *Phoebis philea* | Washington D.C., Denton, Houston, Tampa | 2 |
| Pieridae | *Phoebis sennae* | Atlanta, Charlotte, Chicago, Dallas, Washington D.C., Denton, Des Moines, Detroit, Houston, Los Angeles, New York City, Philadelphia, Phoenix, Raleigh, Riverside, San Diego, St. Louis, Tampa | 234 |
| Pieridae | *Pieris rapae* | Atlanta, Boston, Charlotte, Chicago, Dallas, Washington D.C., Denton, Denver, Des Moines, Detroit, Los Angeles, Minneapolis, New York City, Philadelphia, Phoenix, Raleigh, Riverside, San Diego, San Francisco, St. Louis | 1378 |
| Pieridae | *Pontia beckerii* | Los Angeles, Riverside, San Diego | 1 |
| Pieridae | *Pontia protodice* | Atlanta, Washington D.C., Denton, Denver, Des Moines, Houston, Los Angeles, Minneapolis, Philadelphia, Phoenix, Riverside, San Diego, San Francisco, St. Louis, Tampa | 310 |
| Pieridae | *Pyrisitia lisa* | Atlanta, Charlotte, Chicago, Dallas, Washington D.C., Denton, Des Moines, Houston, Minneapolis, Philadelphia, Raleigh, St. Louis, Tampa | 126 |
| Pieridae | *Zerene cesonia* | Chicago, Washington D.C., Denton, Houston, Los Angeles, Phoenix, Riverside, St. Louis | 19 |
